# Accumulated Cytotoxicity Induced by Islet Amyloid Polypeptide Oligomers in Type 2 Diabetes

**DOI:** 10.64898/2026.06.26.734712

**Authors:** Andrey V. Kuznetsov

## Abstract

Type 2 diabetes is characterized by the progressive aggregation of islet amyloid polypeptide (IAPP) within the islets of Langerhans, a process strongly implicated in beta-cell dysfunction and loss. Although oligomeric IAPP intermediates are widely considered the principal cytotoxic species, the relative contributions of the numerous biological and kinetic processes governing their formation, clearance, and conversion into fibrils remain poorly quantified. Here, a mathematical model of IAPP aggregation is developed that incorporates the physiology of beta-cell secretion and the microanatomy of the islet, including capillary-mediated clearance, enzymatic degradation, and the kinetics of oligomer and fibril formation within a well-mixed control volume. Building on the hypothesis that oligomers are the major cytotoxic species, the concept of accumulated cytotoxicity is introduced, defined as the time integral of the oligomer concentration, and a systematic sensitivity analysis of this quantity with respect to all model parameters is conducted. The results reveal a clear hierarchy: only two parameters, the basal rate of IAPP monomer secretion and the rate constant for spontaneous oligomer dissociation, exert a first-order influence on long-term accumulated cytotoxicity, with dimensionless sensitivities approaching +1 and −1, respectively, while the effects of all other parameters remain subordinate and diminish at long times. The model further shows that capillary clearance, owing to the physical exclusion of oligomers from fenestrated capillaries, selectively reduces fibril accumulation and amyloid deposition without affecting oligomer-mediated cytotoxicity, indicating that amyloid area fraction, a commonly used histological metric of disease severity, may not be a reliable surrogate for cytotoxic burden. At the highest secretion rate, the model predicts approximately 48% replacement of the islet area by amyloid after 30 years, substantially below the more than 85% reported in some end-stage histological studies; this gap likely reflects the omission of late-stage processes such as beta-cell death and release of intracellular IAPP from the present model. These findings identify monomer secretion and oligomer dissociation as potentially promising therapeutic targets for limiting cytotoxic damage in type 2 diabetes and provide a quantitative framework for evaluating candidate intervention strategies.

## 1. Introduction

Type 2 diabetes is characterized not only by insulin resistance and progressive beta-cell dysfunction, but also by a distinctive pathological hallmark: the deposition of islet amyloid within the islets of Langerhans. Post-mortem histopathological studies have identified these deposits in more than 90% of individuals with type 2 diabetes (Clark *et al*., 1996), (Jaikaran & Clark, 2001), (Shome *et al*., 2025), making islet amyloidosis one of the most consistent pathological correlates of the disease. The principal constituent of these deposits is islet amyloid polypeptide (IAPP), also known as amylin, a 37-residue peptide hormone that is co-synthesized, co-packaged, and co-secreted with insulin from pancreatic beta cells (Westermark, 2011).

Under healthy physiological conditions, IAPP is maintained in a soluble, monomeric form within beta-cell secretory granules, a stability that is maintained by the low pH and high local concentrations of insulin and zinc within the granule (Westermark & Westermark, 2008), (Clark & Nilsson, 2004). Insulin, in particular, is a potent inhibitor of IAPP fibrillization in vitro (Westermark & Westermark, 2008), and this protective heterodimerization is thought to suppress aggregation despite the intrinsically high amyloidogenic propensity of human IAPP.

In type 2 diabetes, this protective balance is disrupted: chronic hypersecretion driven by insulin resistance, together with impaired proIAPP processing within beta-cell secretory granules, increases the proportion of incompletely processed proIAPP relative to mature IAPP while reducing the local insulin-to-IAPP ratio, thereby promoting IAPP misfolding and aggregation (Clark & Nilsson, 2004), (Kanatsuka *et al*., 2018).

IAPP aggregation into amyloid fibrils is generally understood to proceed through three kinetically distinct phases: a lag phase, during which small soluble oligomers form from monomeric peptide via primary nucleation; a growth phase, during which protofibrils elongate and additional aggregates form via fibril-surface-catalyzed secondary nucleation; and a plateau phase, during which mature amyloid fibrils dominate (Hassan *et al*., 2022). Although this nucleation-dependent polymerization mechanism has been extensively characterized in vitro (Buchanan *et al*., 2013), (Buell, 2019), typically using a fixed, non-replenished pool of monomers, the situation modeled in the present study is fundamentally different: IAPP monomers are continuously secreted by pancreatic beta cells into the interstitial space of the islets of Langerhans, meaning that the total monomer pool is not conserved but is instead continuously replenished and depleted through production and clearance. This continuous monomer supply is a central feature of the present model and distinguishes it from closed-system in vitro frameworks. Furthermore, increasing evidence indicates that the earliest steps of IAPP aggregation in vivo occur intracellularly, within the secretory granules themselves. ProIAPP-containing fibrillar deposits have been observed in the granule halo of beta cells in both human IAPP transgenic mice and human islet transplants (Paulsson *et al*., 2006), and it has been proposed that these intracellular aggregates can puncture granule membranes, trigger apoptosis, and, upon cell death, act as extracellular seeds that template further amyloid formation from IAPP secreted by neighboring beta cells (Westermark *et al*., 2000), (Paulsson *et al*., 2006).

A major development in this field over the past two decades has been the recognition that mature amyloid fibrils, despite their conspicuous presence in islet histology, may not be the principal agents of beta-cell toxicity. Instead, small, soluble oligomeric intermediates formed transiently during aggregation are now widely regarded as the dominant cytotoxic species (Haataja *et al*., 2008), (Gurlo *et al*., 2010), (Raleigh *et al*., 2017). These toxic oligomers have been reported to form within the endoplasmic reticulum, Golgi apparatus, and secretory granules of beta cells (Gurlo *et al*., 2010), where they compromise membrane integrity through pore formation and disruption of membrane fluidity, ultimately triggering oxidative stress, mitochondrial dysfunction, and apoptosis (Engel *et al*., 2008), (Raleigh *et al*., 2017). This “toxic oligomer hypothesis” parallels analogous developments in Alzheimer’s disease (AD) research, where soluble amyloid beta (Aβ) oligomers, rather than amyloid plaques, are now considered the principal drivers of synaptic dysfunction and neurodegeneration (McLean *et al*., 1999), (Benilova *et al*., 2012), (Viola & Klein, 2015), a parallel that is reinforced by the substantial sequence and conformational similarity between amylin and Aβ (Mizgalska *et al*., 2025).

A recent series of studies has established quantitative frameworks linking Aβ aggregation kinetics to neurotoxicity and biological aging in AD, including models of senile plaque growth (Kuznetsov, 2024) and the concept of accumulated neurotoxicity, defined as the time integral of the oligomer concentration, which is proposed as a marker of cumulative neuronal damage and biological age (Kuznetsov, 2025), (Kuznetsov, 2026a), (Kuznetsov, 2026b). The demonstrated utility of this framework in the neurodegenerative context, together with the well-documented mechanistic parallels between Aβ and IAPP aggregation, motivates extending this framework to islet amyloidosis.

This paper develops a mathematical model of IAPP aggregation in type 2 diabetes. The model equations are derived by adapting the framework for Aβ plaque formation developed in (Dear *et al*., 2020) and extended in (Kuznetsov, 2026b). The key modifications introduced relative to these models are IAPP monomer production by beta cells, clearance of IAPP monomers via fenestrated capillaries, and the release of preassembled oligomers by stressed beta cells, a mechanism that allows the aggregation process to bypass the lag phase (Gurlo *et al*., 2010). Adopting the hypothesis that oligomeric intermediates are the principal cytotoxic species (Haataja *et al*., 2008), (Raleigh *et al*., 2017), (Gurlo *et al*., 2010), the concept of accumulated cytotoxicity is introduced as the time integral of the IAPP oligomer concentration, thereby quantifying the cumulative damage inflicted on beta cells. A systematic sensitivity analysis of this quantity with respect to all model parameters is then performed, with the aim of identifying which physiological and kinetic processes exert the strongest influence on the long-term cytotoxic burden, and thereby highlighting potentially promising targets for therapeutic intervention.

## 2. Materials and models

### 2.1. Mathematical model for IAPP aggregation

Conservation equations for IAPP monomers, oligomers, fibrillar species of all lengths, and total fibril mass are formulated within the interstitial space of the islet of Langerhans, which is treated as the control volume (CV) (Fig. 1). IAPP monomers enter the CV upon release from secretory granules and are removed by clearance through the fenestrated capillaries. In type 2 diabetes, a small additional influx of preassembled IAPP oligomers enters the CV from dysfunctional secretory granules. Applying conservation of monomers within the CV and normalizing both sides by the interstitial volume of the islet of Langerhans gives the following governing equation:

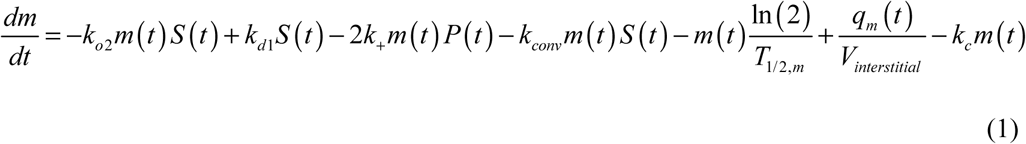

**Fig. 1.**
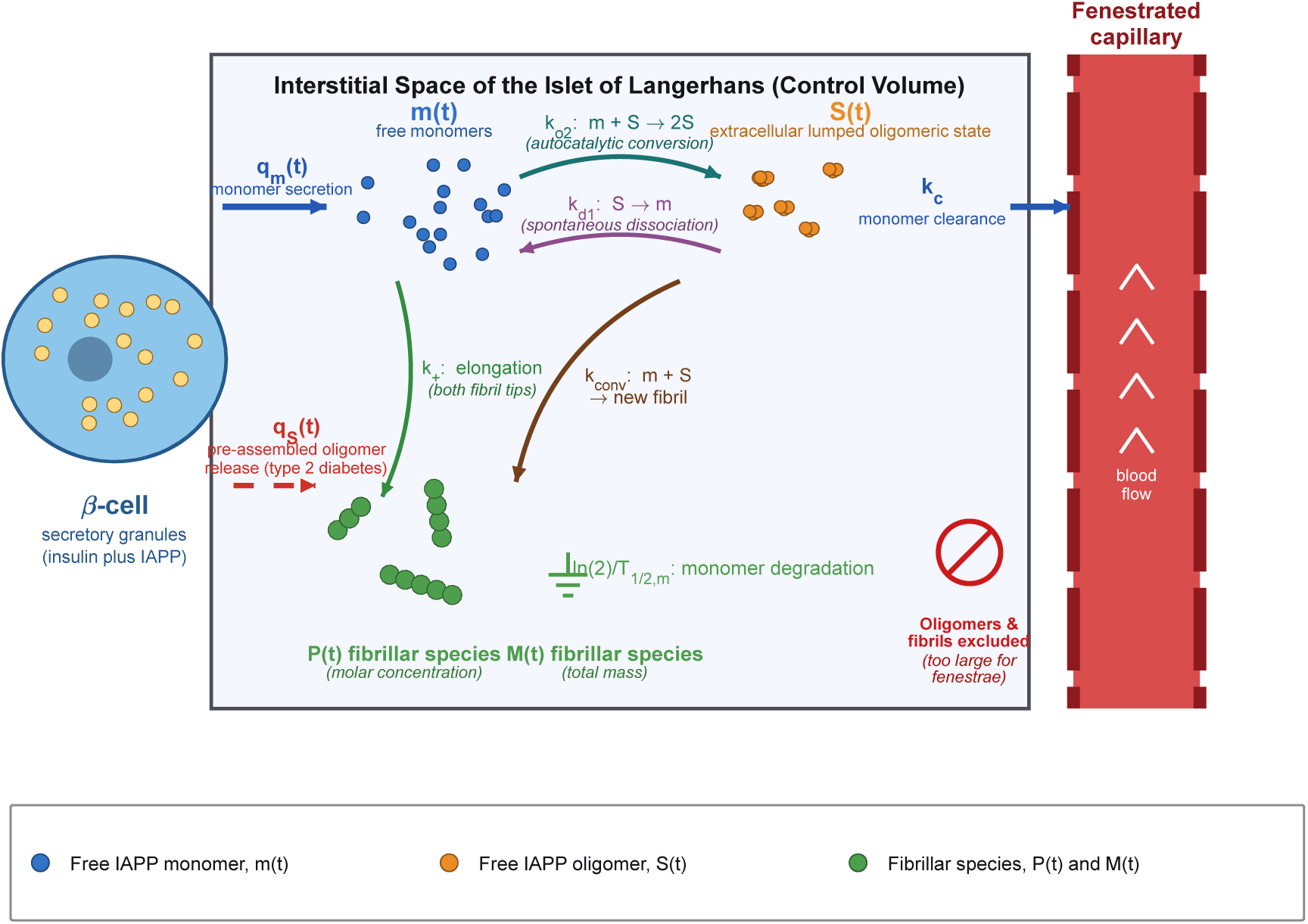
Schematic representation of the mathematical model of IAPP aggregation in the interstitial space of an islet of Langerhans. The model accounts for IAPP secretion by beta cells, monomer and oligomer transport and kinetics, aggregate formation and growth, clearance through fenestrated capillaries and proteolytic degradation, amyloid deposition, and the resulting cytotoxic effects on beta cells.

where *t* is the time elapsed since diabetes onset. Each term on the right-hand side of Eq. (1) represents a distinct physical process. The first term accounts for the utilization of IAPP monomers through secondary, or autocatalytic, conversion, a process in which existing oligomers catalyze the formation of new ones. Monomer conversion via nucleation is assumed to be negligible compared to oligomer-catalyzed conversion, because in type 2 diabetes oligomers can form by misfolding within secretory granules and subsequently be released into the interstitial space, where they can seed autocatalytic oligomer formation. The second term describes monomer production arising from the spontaneous dissociation of oligomers into monomers; the corresponding kinetic constant is taken from (Pillay & Govender, 2014). Oligomer dissociation catalyzed by fibril surfaces was neglected, as no data for this process were reported in that study. The third term describes monomer utilization due to fibril elongation. Because each fibril has two free tips to which monomers can attach, a factor of 2 appears in this term. The fourth term represents the conversion of oligomers into elongation-competent fibrils. The product of *m* (*t* ) and *S* (*t* ) reflects the assumption that this conversion proceeds via monomer addition, consistent with (Dear *et al*., 2020). The fifth term describes the removal of monomers through proteolytic degradation, with the rate of this process governed by the half-life of IAPP monomers, *T*_1/ 2,*m*_ . The sixth term represents the volumetric generation rate of monomers in the interstitial space of the islet of Langerhans (the CV), where *q_m_* (*t* ) denotes the rate of monomer secretion by beta cells within a single islet. The seventh term represents the loss of IAPP monomers driven by diffusion through the interstitial space into the fenestrated capillaries, where they are subsequently removed from the islet by the bloodstream.

For the case with oscillating *q_m_* (*t* )

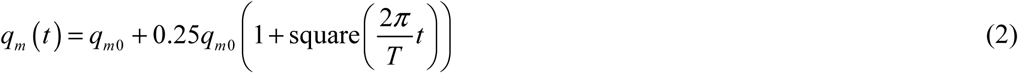

see footnote “e” after Table 2. In Eq. (2), square(*t*) denotes a unit square wave of period 2π, defined analogously to the sine function in that it oscillates between −1 and +1 with the same period, but differs in that its transitions between states are instantaneous rather than continuous. This produces a piecewise-constant waveform that alternates between two fixed values at regular intervals.

**Table 1.** Dependent variables utilized within the computational model.

| Symbol | Definition | Units |
| --- | --- | --- |
| $m(t)$ | Molar concentration of free IAPP monomers<br>(number of free IAPP monomers per unit<br>volume) | $\mu\text{M}$ |
| $M(t)$ | Fibril mass concentration: concentration of<br>IAPP monomer equivalents incorporated into | $\mu\text{M}$ monomer<br>equivalents |
|  | all fibrils, used as a measure of total fibril mass per unit volume |  |
| $P(t)$ | Fibril particle concentration: molar concentration of fibril particles, irrespective of fibril length | $\mu\text{M}$ |
| $S(t)$ | Extracellular concentration of the lumped on-pathway IAPP oligomeric/activated state within the interstitial CV, expressed in monomer-equivalent molar units | $\mu\text{M}$ monomer equivalents |
| $\Xi(t)$ | Accumulated oligomer exposure, used as the model measure of accumulated cytotoxicity,<br>$\Xi(t) = \int_0^t S(\hat{t}) d\hat{t}$ | $\mu\text{M s}$ |

**Table 2.**
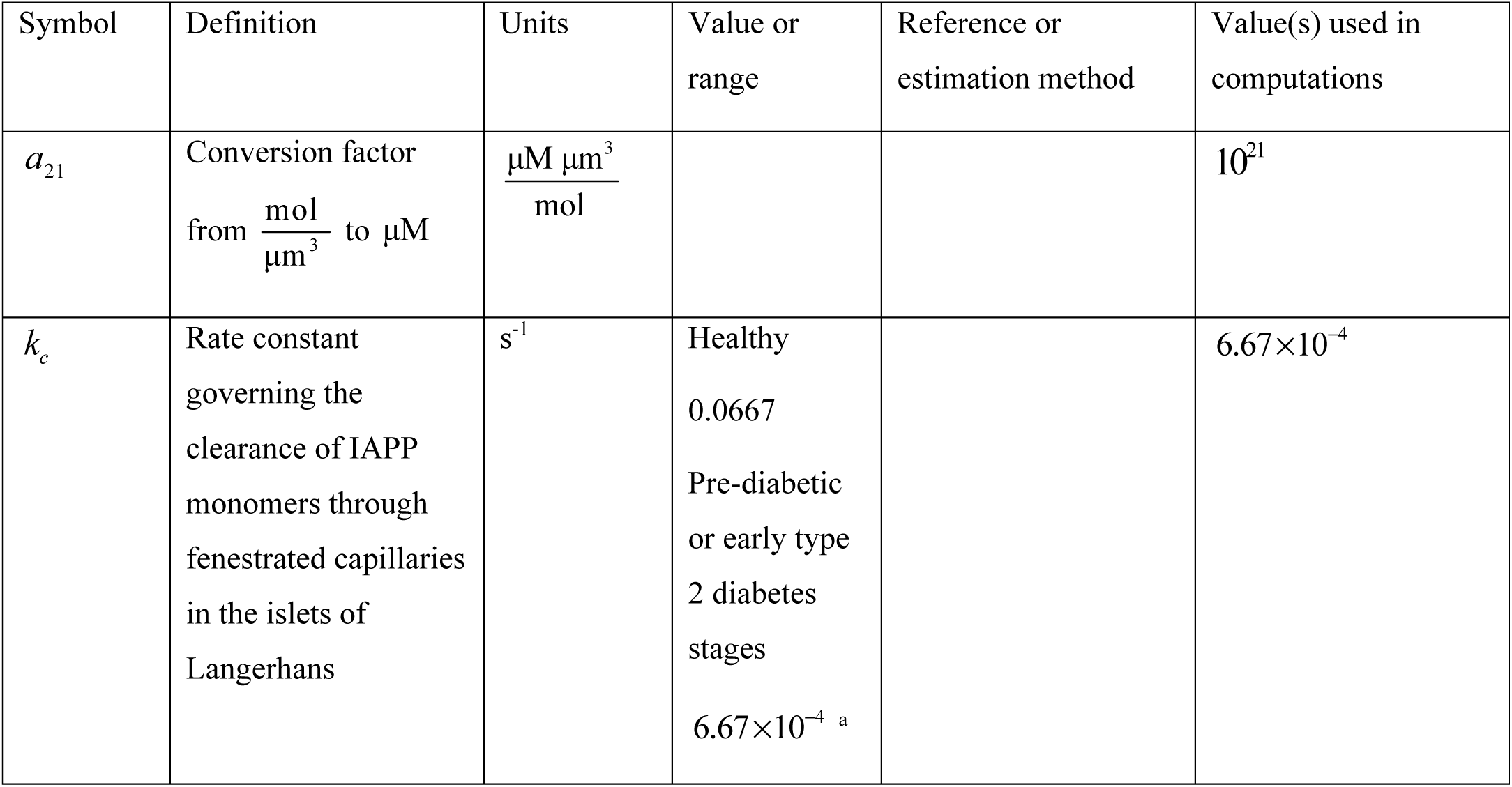
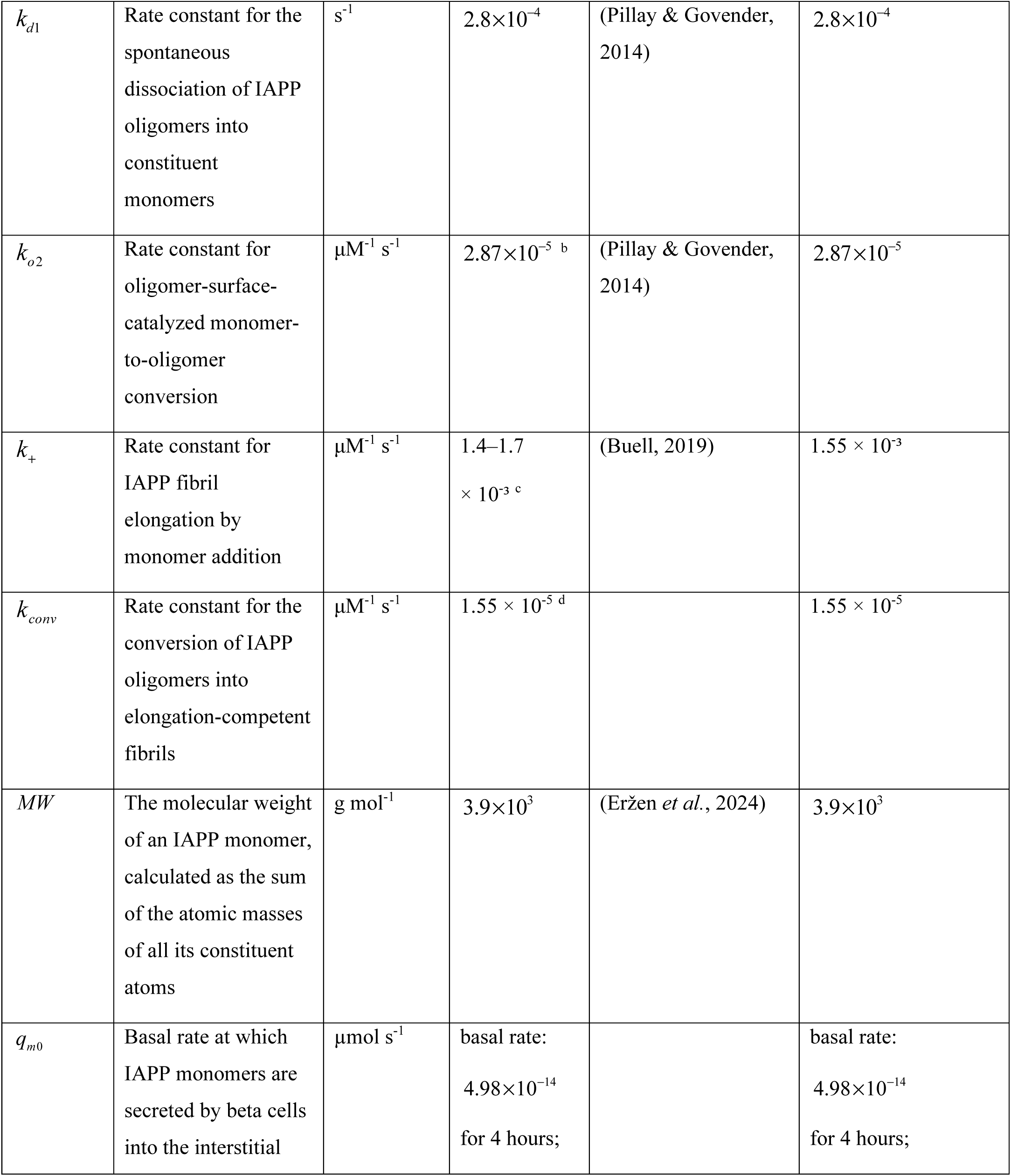
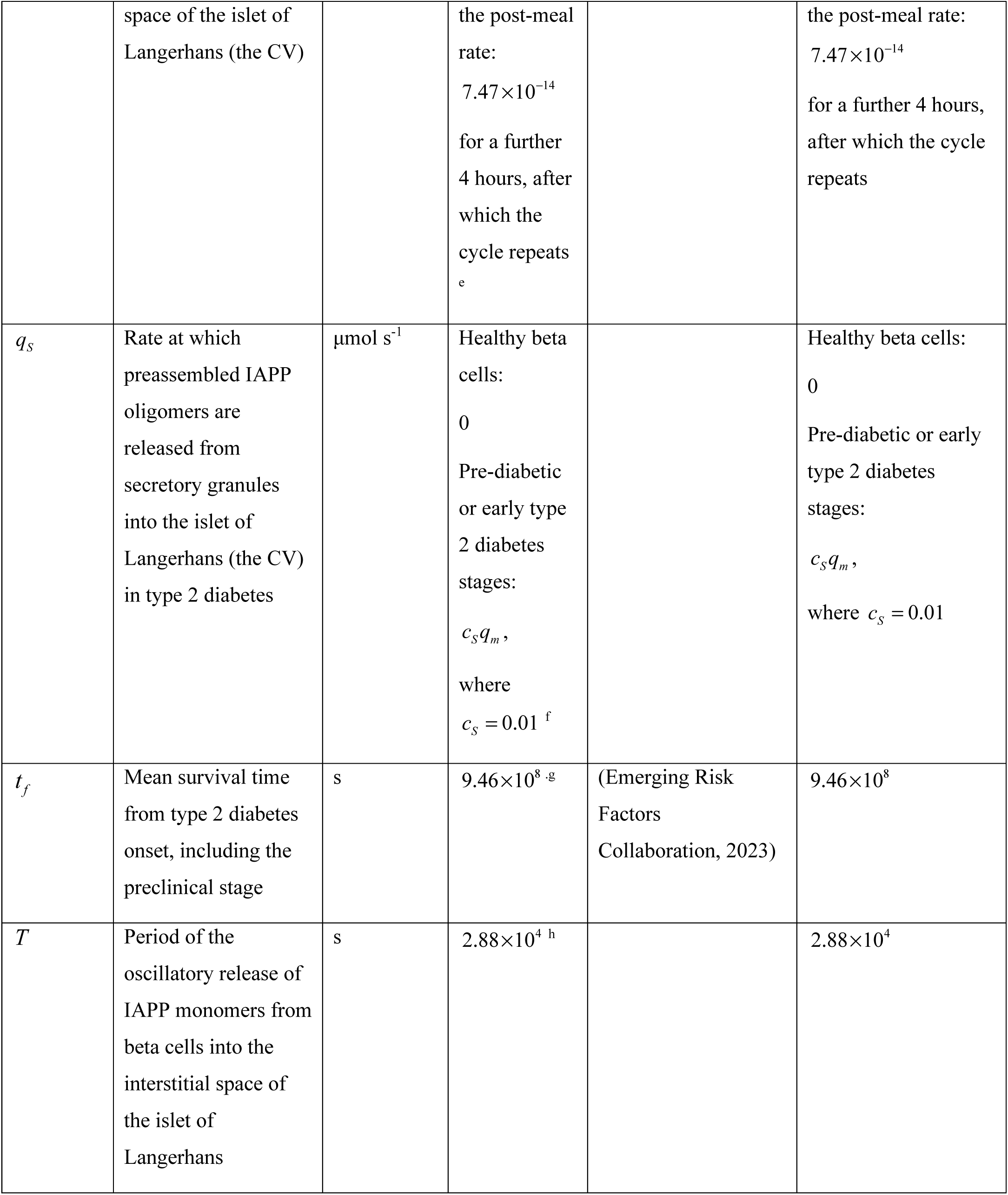
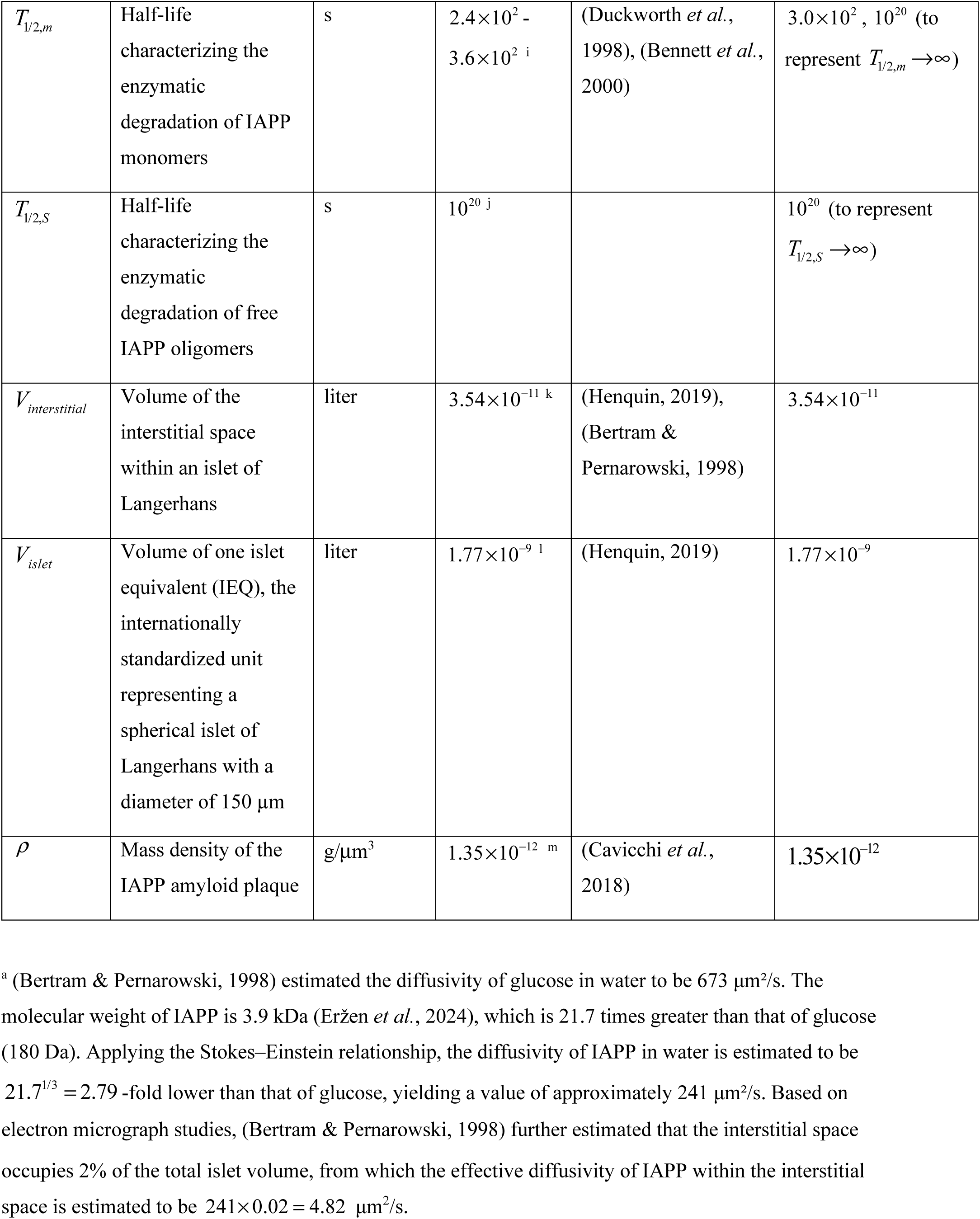
Model parameters.

For the constant-release formulation, *q_m_* is defined as the temporal mean of the oscillatory release rate given by Eq. (2), averaged over one full oscillation period *T*, ensuring that the total IAPP monomer input is the same in the two formulations.

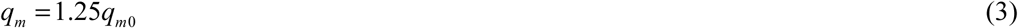

Adopting the Finke-Watzky framework, in which an oligomer is modeled as an activated monomer with identical mass (Morris *et al*., 2008), (Iashchishyn *et al*., 2017), the stepwise assembly of oligomeric intermediates is represented by a single lumped activation step. This is a deliberate simplification that reduces the complexity of the model while preserving the essential features of the nucleation pathway.

In the present lumped formulation, *S* (*t* ) represents the extracellular concentration of on-pathway IAPP oligomers within the interstitial CV (Fig. 1). In keeping with the Finke–Watzky framework adopted here, oligomers are assumed to carry the same mass as monomers, and their concentration is accordingly expressed in monomer-equivalent molar units. Intracellular IAPP oligomers are not represented by an independent state variable; rather, their release from dysfunctional secretory granules is captured by the source term *q_S_* (*t* ) , after which the released oligomers contribute to the extracellular pool described by *S* (*t* ) . The conservation equation for IAPP oligomers within the CV, normalized by the interstitial volume within an islet of Langerhans, takes the form:

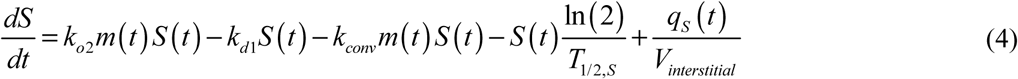

The first term on the right-hand side of Eq. (4) represents the gain of IAPP oligomers through autocatalytic conversion from monomers. This term has the same magnitude but the opposite sign as the first term on the right-hand side of Eq. (1). The second term describes oligomer loss due to spontaneous dissociation into monomers, and is likewise equal in magnitude but opposite in sign to the corresponding term in Eq. (1). The third term represents oligomer loss arising from their conversion into elongation-competent fibrils. This term has the same magnitude and sign as the fourth term on the right-hand side of Eq. (1), consistent with the assumption that one oligomer and one monomer combine to form an elongation-competent fibril (Dear *et al*., 2020). The fourth term describes the removal of oligomers through proteolytic degradation, with the rate of this process governed by the half-life of IAPP oligomers, *T*_1/2,*S*_ . The fifth term describes the volumetric generation rate of oligomers in the CV, where *q_S_* (*t* ) denotes the rate of oligomer secretion by beta cells within a single islet. Oligomer secretion arises from the fact that in type 2 diabetes, oligomers can form through misfolding within secretory granules and subsequently be released into the interstitial space.

Applying the conservation principle to IAPP fibrillar species of all lengths and normalizing by the volume of the CV yields the governing equation for the total fibril concentration, irrespective of fibril length:

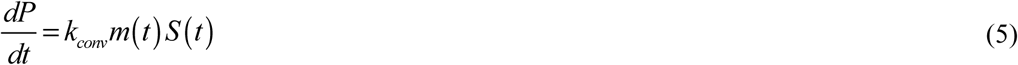

The right-hand side of Eq. (5) corresponds to the fourth term in Eq. (1) and the third term in Eq. (4), with opposite signs in the respective equations, reflecting the fact that oligomer-to-fibril conversion simultaneously depletes both the monomer and oligomer pools. The conversion process is modeled as proceeding through monomer addition: a bimolecular reaction in which one monomer and one oligomer associate to yield a newly formed, elongation-competent fibril. Fibril fragmentation, which is accounted for in the equations of (Dear *et al*., 2020), is not included in Eq. (5) owing to insufficient experimental evidence that this process plays a significant role in IAPP aggregation.

The fibril population within the CV is also described in terms of its total mass, expressed as the number of IAPP monomers incorporated into all fibrils. This moment-based representation eliminates the need to resolve the full fibril-length distribution. Applying the conservation principle to this quantity and normalizing by the CV volume yields:

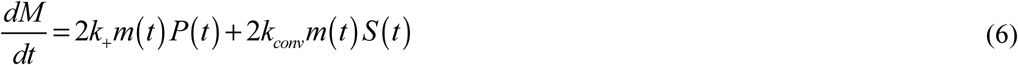

In Eq. (6), the first term on the right-hand side describes elongation of existing fibrils through the addition of free monomers at either fibril tip. The second term accounts for the creation of new fibril mass via oligomer-to-fibril conversion: because each event involves the simultaneous incorporation of one free monomer and one oligomer into the fibrillar phase, the total mass gained per nucleation event is equivalent to two monomers, which is reflected in the factor of 2.

Before the onset of diabetes and severe insulin resistance, healthy physiological mechanisms are thought to prevent the accumulation of toxic IAPP oligomers (Haataja *et al*., 2008). Consistent with this assumption, the initial concentrations of IAPP oligomers and fibrils are assumed to be zero. The initial monomer concentration is obtained from Eq. (1) by assuming that monomer secretion by beta cells is exactly balanced by capillary clearance at the onset of the simulation:

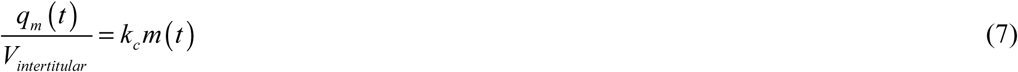

Solving for *m*(0) yields the following initial conditions for Eqs. (1), (4)–(6):

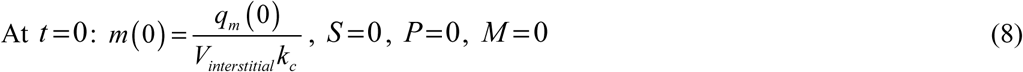

^a^(Bertram & Pernarowski, 1998) estimated the diffusivity of glucose in water to be 673 μm²/s. The molecular weight of IAPP is 3.9 kDa (Eržen *et al*., 2024), which is 21.7 times greater than that of glucose (180 Da). Applying the Stokes–Einstein relationship, the diffusivity of IAPP in water is estimated to be 21.7^1/3^ = 2.79 -fold lower than that of glucose, yielding a value of approximately 241 μm²/s. Based on electron micrograph studies, (Bertram & Pernarowski, 1998) further estimated that the interstitial space occupies 2% of the total islet volume, from which the effective diffusivity of IAPP within the interstitial space is estimated to be 241× 0.02 = 4.82 μm^2^/s.

The characteristic diffusion time is given by *x*^2^ / (2*D*) (Crank, 1975), (Dorrigiv *et al*., 2021), where *x* is the diffusion distance and *D* is the diffusivity. Analysis of the spatial distribution of beta cells relative to the capillaries suggests that the majority of cells lie within 10 μm of the nearest capillary (Cottle *et al*., 2021), yielding a characteristic diffusion time for IAPP of 10.4 s. Taking this diffusion time as an approximation of the IAPP half-life within the islets of Langerhans, the rate constant governing the clearance of IAPP monomers through the fenestrated capillaries of the islets can be estimated as ln (2) / *t*_1/2_ , giving a value of 0.0667 s⁻¹.

In type 2 diabetes, the islet microvasculature undergoes progressive structural remodeling, including thickening of endothelial cells (Brissova *et al*., 2015), (Hogan & Hull, 2017) and a reduction in capillary endothelial fenestration (Li *et al*., 2006). These changes are expected to reduce the rate of IAPP monomer clearance from the interstitial space. Compounding this effect, increased extracellular trapping and deposition of IAPP monomers have been reported in type 2 diabetic islets (Hull *et al*., 2004). It is assumed that the combined effects of reduced microvascular permeability and increased diffusion resistance prolong the interstitial residence time of IAPP monomers in type 2 diabetes by a factor of 100 relative to the healthy state. This assumption translates directly into a 100-fold reduction in the rate constant characterizing the clearance of IAPP monomers from the interstitial space via the fenestrated capillaries.

^b^It is assumed that primary nucleation plays a negligible role in IAPP aggregation relative to secondary nucleation. This is because, in the context of type 2 diabetes, IAPP oligomers may form within secretory granules and subsequently be released into the interstitial space, where they catalyze further IAPP aggregation (Gurlo *et al*., 2010). The secondary nucleation rate constant is assumed to be 28.7 M^-1^ s^-1^ (Pillay & Govender, 2014).

^c^Experimental measurements of the IAPP fibril elongation rate constant, compiled in (Buell, 2019) from single-particle AFM and TIRF studies, yield values of approximately 1.4–1.7 × 10³ M⁻¹ s⁻¹.

^d^Although the oligomer-to-fibril conversion rate constant is difficult to measure directly, because of the transient and heterogeneous nature of the oligomeric intermediate (Dear *et al*., 2020), indirect evidence for the assumption that *k_conv_* << *k*_+_ comes from the pronounced lag phase observed in IAPP aggregation kinetics. If oligomer-to-fibril conversion were as fast as fibril elongation, no lag phase would be expected, as fibrils would begin accumulating immediately upon the appearance of oligomers (Buchanan *et al*., 2013). Accordingly, *k_conv_* = 0.01*k*_+_ is adopted as a modeling assumption in the present work.

^e^The number of insulin molecules per secretory granule has been estimated at 300,000 (Rutter *et al*., 2015). IAPP is co-synthesized and co-packaged with insulin into secretory granules, and is co-secreted at a ratio to insulin of approximately 1:100 (Germanos *et al*., 2021). Dividing 300,000 by 100 yields an estimate of 3,000 IAPP molecules per secretory granule.

A typical islet of Langerhans contains approximately 1,000 beta cells (Henquin, 2019). According to the data reported in (Rorsman & Renström, 2003), beta cells in mouse islets release approximately 15 granules per minute during the first phase of insulin secretion and 5 granules per minute during the sustained second phase. (Low *et al*., 2013) reported that granule fusion events per beta cell increase ninefold as glucose rises from basal (3 mmol/l) to maximally stimulating (20 mmol/l) concentrations. Taking the second-phase rate as the reference and dividing it by this ninefold factor yields a basal exocytosis rate of approximately 0.6 granules per beta cell per minute.

(Alcazar & Buchwald, 2019) reported that the maximal insulin secretion rate of human islets is approximately threefold lower than that of murine islets. Applying this threefold scaling factor to the mouse-derived basal estimate yields an approximate basal exocytosis rate of 0.2 granules per beta cell per minute in human islets. The same scaling applied to the second-phase rate gives approximately 1.67 granules per beta cell per minute.

The total basal secretion rate for an islet containing approximately 1,000 beta cells is thus estimated at 200 granules per minute, increasing to approximately 1,670 granules per minute during the second phase. With approximately 3,000 IAPP molecules per secretory granule, the total basal IAPP release rate amounts to approximately 600,000 molecules per minute per islet (10,000 molecules per second, or approximately 1.66 ×10_−14_ µmol/s). During the second phase, this rate rises to approximately 5,010,000 molecules per minute per islet (83,500 molecules per second, or approximately 1.39 ×10_−13_ µmol/s).

Basal fasting plasma insulin concentrations in patients with prediabetes or early-stage type 2 diabetes are two to four times greater than in healthy subjects (Solis-Herrera *et al*., 2025). Since IAPP is co-packaged and co-secreted with insulin in the secretory granules of pancreatic beta cells, the basal rate of IAPP release in prediabetic or early-stage type 2 diabetic patients is assumed to be three times that of healthy individuals, giving a value of 4.98 ×10_−14_ µmol/s per islet. Because beta cells in patients with prediabetes and early-stage type 2 diabetes are assumed to be functionally impaired, with depleted secretory granules and a completely abolished first-phase insulin spike (Polonsky *et al*., 1988), (DeFronzo, 2009), they are assumed to exhibit only a modest 1.5-fold increase in IAPP release following a meal, bringing the post-meal release rate to 7.47 ×10_−14_ µmol/s per islet. It is thus assumed that in pre-diabetic and early-stage type 2 diabetic patients, beta cells release IAPP at the basal rate of 4.98 ×10_−14_ µmol/s per islet for 4 hours, followed by the post-meal rate of 7.47 ×10_−14_ µmol/s per islet for a further 4 hours, after which the cycle repeats.

^f^ The acidic, insulin-rich, zinc-containing environment of beta-cell secretory granules suppresses IAPP self-association (Nedumpully-Govindan & Ding, 2015), making the formation of preassembled oligomers unlikely under healthy conditions. For this reason, *q_S_* is assumed to be zero in healthy beta cells.

In type 2 diabetes, impaired processing of proIAPP, combined with reduced insulin co-packaging, which normally suppresses IAPP fibrillogenesis, and a loss of the protective acidic granular environment, allows IAPP to misfold and assemble into toxic oligomers intracellularly within the secretory granules (Clark & Nilsson, 2004), (Kanatsuka *et al*., 2018). These oligomers disrupt the membranes of secretory granules and, upon escaping into the cytoplasm, attack mitochondrial membranes, ultimately leading to beta-cell death, while a secreted fraction seeds extracellular amyloid deposits (Gurlo *et al*., 2010). For modelling purposes, and in the absence of direct experimental measurements, the rate at which preassembled oligomers are released from beta cells under type 2 diabetes conditions is assumed to be a small fraction of the monomer release rate,

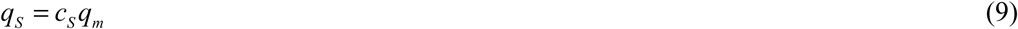

where the proportionality factor, *c_S_* , is assumed to be 0.01.

^g^Among individuals surviving to age 50, the reduction in life expectancy associated with a type 2 diabetes diagnosis was approximately 14, 10, and 6 years for those diagnosed at ages 30, 40, and 50 years, respectively, relative to age-matched individuals without diabetes (Emerging Risk Factors Collaboration, 2023). Assuming an average life expectancy of 80 years, the duration of living with type 2 diabetes may therefore be estimated to be approximately 30 years.

^h^8 hours, consistent with a three-meal-per-day feeding pattern in which meals are approximately equally spaced throughout the day.

^i^To estimate the catabolic half-life of IAPP monomers, insulin was used as a kinetic proxy. The principal protease responsible for degrading IAPP in the interstitial space is insulin-degrading enzyme (IDE), which has been shown in vitro to degrade IAPP with comparable kinetics (Bennett *et al*., 2000). Given that IAPP is co-secreted alongside insulin into the same interstitial compartment and competes with insulin for the same active site on IDE, it is reasonable to assume that both peptides share a similar catabolic half-life. The plasma half-life of insulin has been measured at 4 to 6 minutes (Duckworth *et al*., 1998), and this value is accordingly adopted for IAPP.

^j^IDE, the primary enzyme responsible for the catabolism of IAPP monomers (Bennett *et al*., 2000), contains an enclosed catalytic chamber that imposes stringent size constraints on its substrates (Shen *et al*., 2006). This structural limitation renders IDE incapable of degrading IAPP oligomers and fibrils, providing a mechanistic rationale for treating their catabolic half-lives as effectively infinite in the present model.

^k^By international standards, one islet equivalent (IEQ) is defined as a perfect sphere with a diameter of 150 µm (Henquin, 2019), corresponding to a volume of 1.77 ×10_−9_ liters. Based on electron micrograph studies, the extracellular space accounts for only 1–2% of the total islet volume (Bertram & Pernarowski, 1998). Adopting the upper estimate of 2%, the interstitial volume of a single islet of Langerhans is estimated at approximately 3.54 ×10_−11_ liters.

^l^The volume of a perfect sphere with a diameter of 150 µm, which approximately equals 1.77 ×10_−9_ liters (Henquin, 2019).

^m^The average density of proteins, regardless of their specific type, is commonly approximated as 1.35 g cm⁻³ (Cavicchi *et al*., 2018).

### 2.2. Modeling amyloid deposit volume in pancreatic islets

The temporal evolution of the volume occupied by amyloid deposits is modeled following the framework of (Kuznetsov, 2024). To track the growth of IAPP plaques (Fig. 1), the total number of IAPP monomers, *N*, incorporated into the plaques as a function of time *t* was monitored. Adopting the approach of (Watzky *et al*., 2008), this quantity is derived from the mean concentration of IAPP monomers deposited into the plaque:

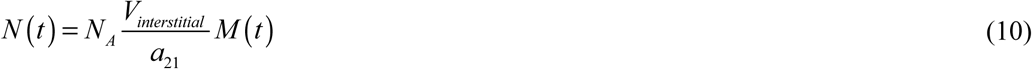

where *N_A_* is Avogadro’s number. The total number of deposited monomers can also be related to the volume of the growing plaques. Following (Watzky *et al*., 2008), *N* is expressed in terms of the volume *V_IAD_* occupied by the IAPP plaques within the islet of Langerhans:

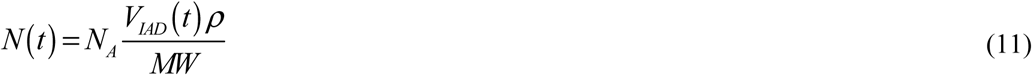

where *MW* is the mean molecular weight of an IAPP monomer. Combining Eqs. (10) and (11) by equating their right-hand sides and solving for *V_IAD_* yields:

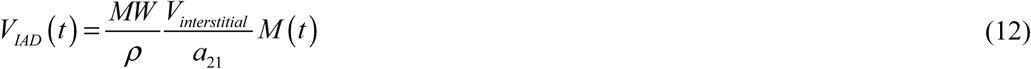

Where 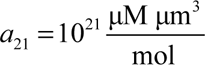 is a unit conversion factor that converts from 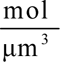 to μM .

Because *M* (*t*) is expressed as a molar concentration of monomer equivalents incorporated into fibrils, multiplication by *V_interstitial_* gives the corresponding amount of deposited IAPP within the interstitial control volume. The conversion factor *a*_21_ converts the resulting amount from the concentration units used in the model to moles. Multiplication by Avogadro’s number then gives the total number *N* (*t* ) of IAPP monomer equivalents incorporated into the amyloid deposits, as expressed by Eq. (10).

Alternatively, multiplication of the number of moles by the IAPP molecular weight gives the total fibrillar protein mass, and division by the assumed amyloid-protein density converts this mass into the amyloid-deposit volume *V_IAD_* , yielding Eqs. (11) and (12).

The volumetric fraction of the islet of Langerhans occupied by IAPP amyloid deposits is defined as:

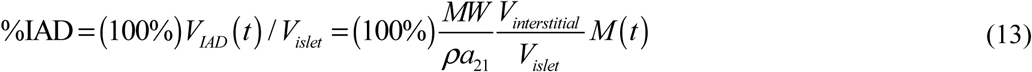

Eq. (13) gives the three-dimensional volume fraction of the islet occupied by amyloid. To compare this prediction with histological measurements reported as two-dimensional amyloid area fractions, the Delesse stereological principle (Weibel, 1979) is invoked: for representative, unbiased tissue sections, the expected area fraction occupied by a phase equals its volume fraction. Accordingly, *V_IAD_*/ *V_islet_* is interpreted here as the expected histological amyloid area fraction. This equivalence is a stereological approximation and need not hold exactly for an individual section, particularly when amyloid deposition is spatially heterogeneous.

### 2.3. Cytotoxicity arising from the accumulation of IAPP oligomers

There is growing evidence that IAPP oligomers are more cytotoxic than mature amyloid fibrils. These oligomers disrupt intracellular organelle membranes, including those of mitochondria, and are ultimately secreted into the extracellular space (Gurlo *et al*., 2010). The hypothesis that IAPP aggregates are the principal causative agents of beta-cell loss in type 2 diabetes is critically examined in (Haataja *et al*., 2008), (Zraika *et al*., 2010). The mechanisms by which IAPP oligomers exert their cytotoxic effects remain incompletely understood; however, two principal pathways have been proposed: the formation of pore-like structures in the lipid bilayer and the disruption of normal membrane fluidity, both of which are thought to compromise membrane integrity and ultimately trigger beta-cell apoptosis (Raleigh *et al*., 2017).

These membrane-disruptive mechanisms suggest that the damage inflicted by IAPP oligomers on a beta cell is cumulative: each pore-forming or membrane-destabilizing event represents an incremental insult that compromises cellular integrity, and the resulting stress, including organelle dysfunction and activation of apoptotic signaling, is unlikely to be fully repaired before subsequent exposure occurs. Under this view, it is not the instantaneous oligomer concentration at any single moment that determines a cell’s fate, but rather the total exposure accumulated over the course of disease progression. A beta cell exposed to a moderate oligomer concentration for a prolonged period may therefore sustain damage comparable to, or greater than, a cell exposed briefly to a much higher concentration. This motivates characterizing IAPP-induced cytotoxicity using the time integral of the oligomer concentration, rather than by any single instantaneous measurement, since only the integrated exposure captures the full history of membrane insult and apoptotic stress experienced by the cell.

Accumulated cytotoxicity of IAPP oligomers is therefore defined, following (Kuznetsov, 2025), as:

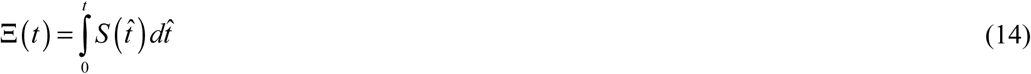

Thus, Ξ(*t* ) is a dimensional integrated-exposure measure rather than a dimensionless measure of cellular damage. Its units are concentration multiplied by time; when *S* is expressed in μM and time in seconds, Ξ has units of μM s. The term “accumulated cytotoxicity” is used here as shorthand for this cumulative exposure of beta cells to the modeled oligomeric state.

### 2.4. Approximate analytical solution for constant secretion rates of IAPP monomers and oligomers: the limiting case of *T*_1/ 2,*m*_ →∞ and *T*_1/ 2,*S*_ →∞

For this limiting case, summing Eqs. (1), (4), and (6) yields:

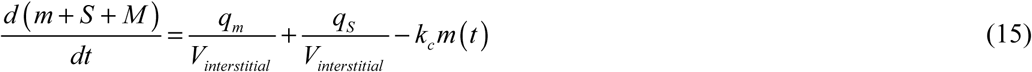

The reduction of Eq. (15) used below is a long-time asymptotic approximation rather than an exact identity. The analytical forms derived below show that *m* = *O* (*t* ^−1/2^ ) and *S* = *O* (1) , whereas the total fibril mass satisfies *M* = *O* (*t* ) . Consequently, (*m* + *S* ) / *M* → 0 as *t* →∞ , which provides the asymptotic justification for neglecting *m* + *S* relative to *M*. In addition, the capillary-clearance term satisfies *k m* = *O* (*t* ^−1/2^ ) and is therefore asymptotically smaller than the constant secretion source. This allows Eq. (15) to be integrated analytically to give:

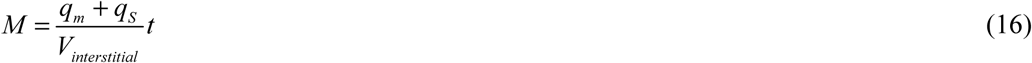

Eq. (16) should be interpreted as the leading-order long-time approximation for *M* (*t* ) , rather than as an exact solution of Eq. (15). Using the equations and parameter values stated in Table 2, (*m* + *S*)⁄*M* was estimated to be 1.96 ×10_−7_ , 4.08 ×10_−8_ , and 3.22 ×10_−8_ at 30 years for the low, baseline, and highest secretion rates, respectively.

Inspection of the numerical results in Figs. 2b, 3a, and 4a reveals that the time evolution of *m*, *S*, and *P* can be well approximated by:

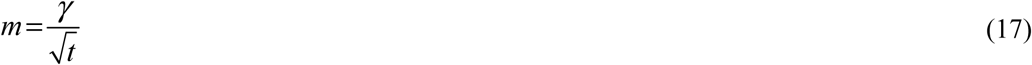

**Fig. 2.**
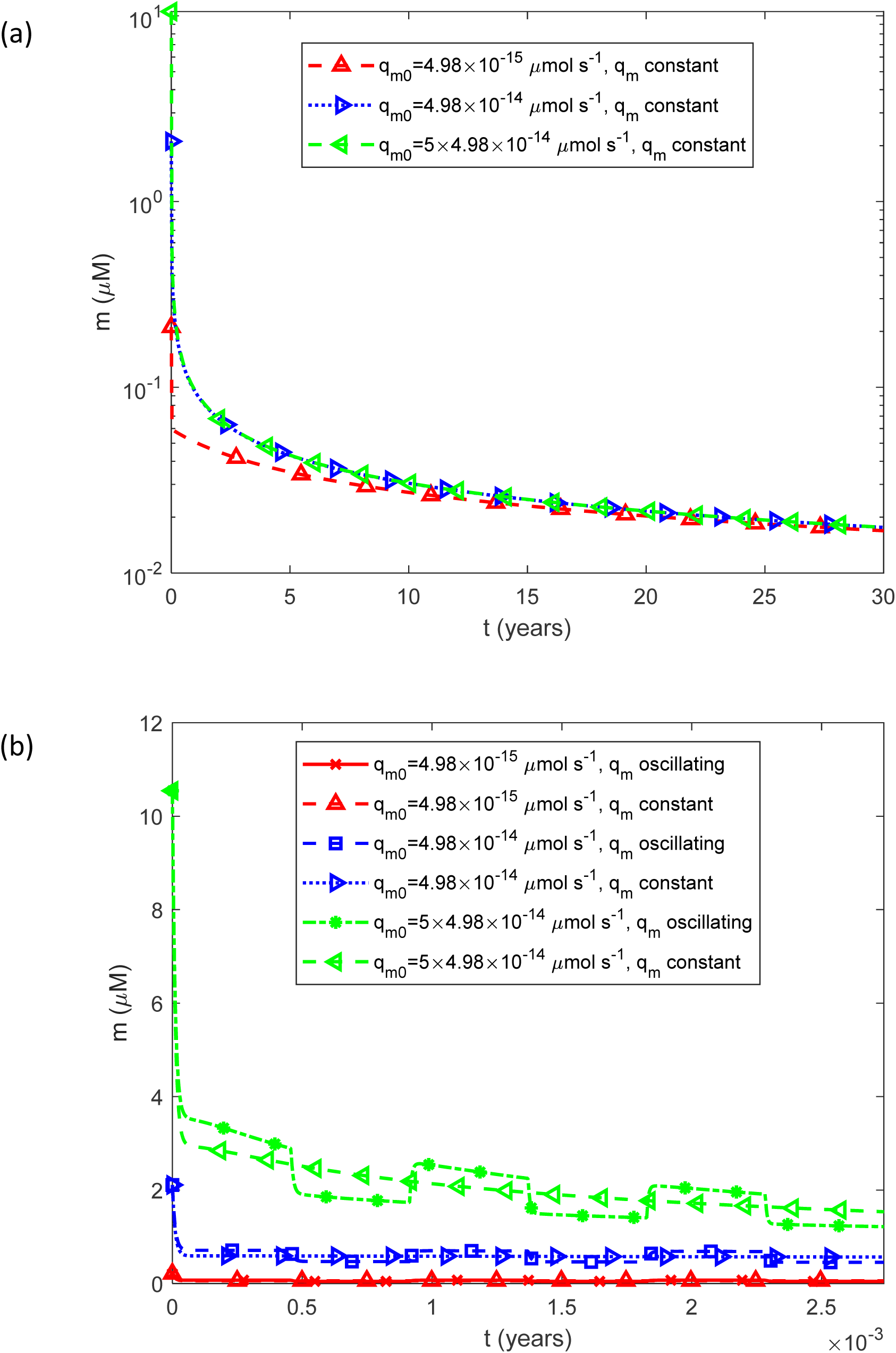
(a) Molar concentration of free IAPP monomers as a function of time, *m*(*t*) , for a constant (time-averaged) monomer release rate of 1.25*q_m_* _0_ . (b) As in (a), but restricted to the time interval [0, 0.0027 years], equivalent to 1 day, illustrating the effect of oscillatory monomer release on the short-term dynamics of the free monomer concentration. Results are shown for three representative values of the basal IAPP monomer secretion rate by beta cells, *q_m_* _0_ .

**Fig. 3.**
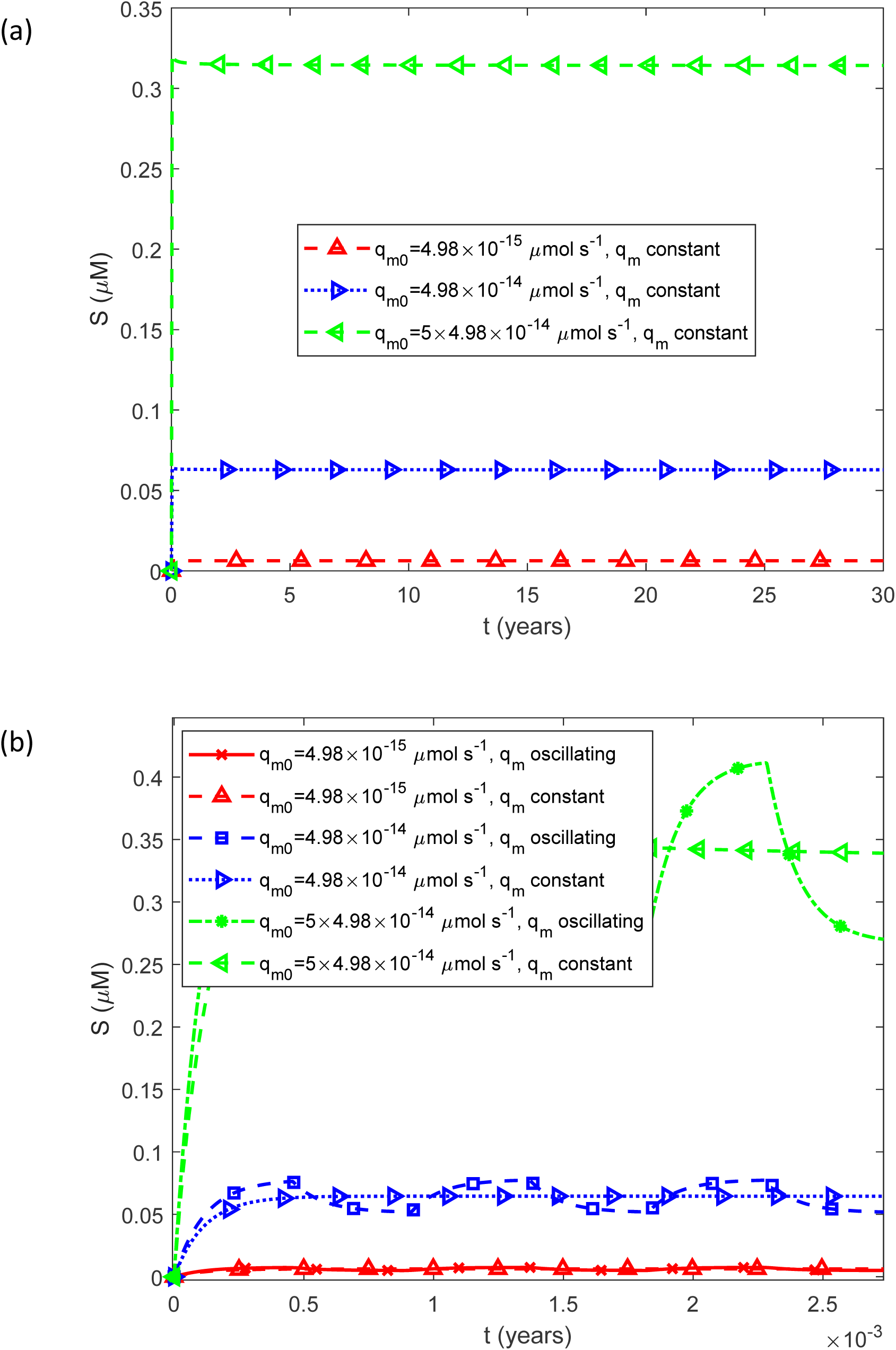
(a) Molar concentration of free IAPP oligomers as a function of time, *S*(*t*) , for a constant (time-averaged) monomer release rate of 1.25*q_m_* _0_ . (b) As in (a), but restricted to the time interval [0, 0.0027 years], equivalent to 1 day, illustrating the effect of oscillatory monomer release on the short-term dynamics of the free oligomer concentration. Results are shown for three representative values of the basal IAPP monomer secretion rate by beta cells, *q_m_* _0_ .

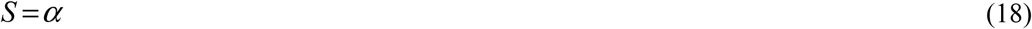

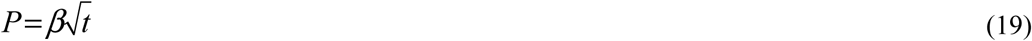

where *α* , *β* , and *γ* are constants to be determined. Substituting these expressions into Eq. (5) gives:

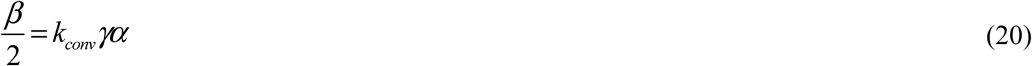

Inserting Eqs. (17) and (18) into Eq. (4) leads to:

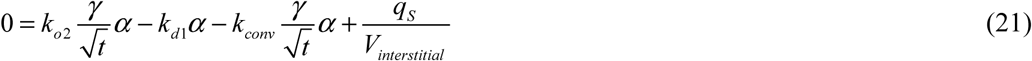

Seeking an approximate solution valid in the long-time limit ( *t* →∞ ) gives:

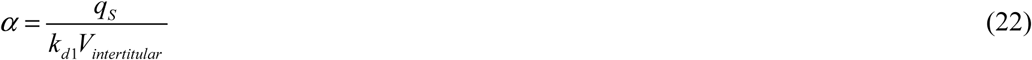

Substituting Eqs. (17)–(19) into Eq. (1) produces:

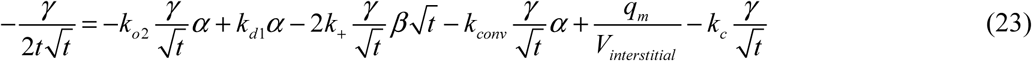

In the long-time limit ( *t* →∞ ), the approximate solution takes the form:

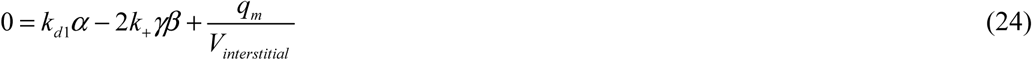

Inserting Eq. (22) into Eqs. (20) and (24) and solving simultaneously for *β* and *γ* gives:

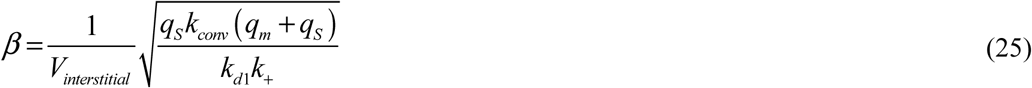

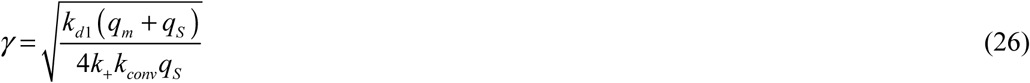

With *α* , *β* , and *γ* determined, substituting Eqs. (22), (25), and (26) into Eqs. (17)–(19) gives the approximate analytical solutions:

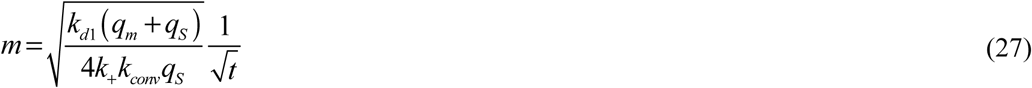

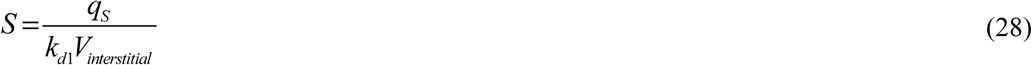

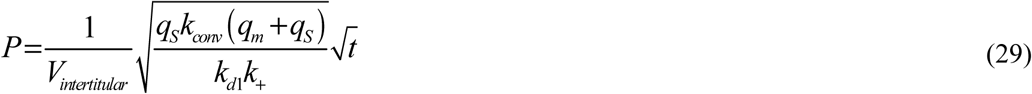

Inserting Eq. (16) into Eq. (13) yields:

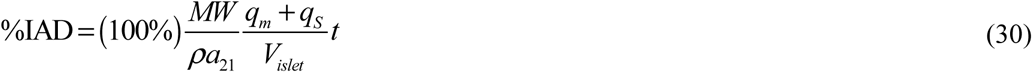

With Eq. (28) established, substituting it into Eq. (14) produces:

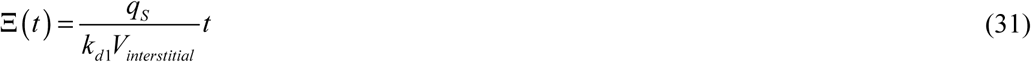

Solving Eq. (30) for *t* and substituting the resulting expression into Eq. (31) yields:

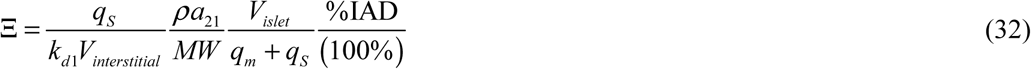

### 2.5. Sensitivity of accumulated cytotoxicity to model parameters

To characterize how each model parameter affects accumulated cytotoxicity, local sensitivity coefficients were computed as first-order partial derivatives of accumulated cytotoxicity with respect to each individual parameter (Beck & Arnold, 1977), (Sadegh Zadeh & Montas, 2010), (Zi, 2011), (Kuznetsov & Kuznetsov, 2019). For example, the sensitivity of accumulated cytotoxicity with respect to the basal rate of IAPP monomer secretion by beta cells was evaluated using a finite-difference approximation:

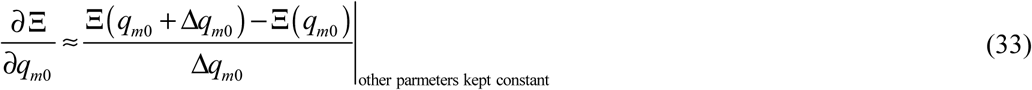

Here, Δ*q_mo_* = 10^−3^ *q_mo_* denotes the step size used in the numerical differentiation. To ensure independence from the choice of step size, several step sizes were tested, and the resulting sensitivity coefficients were found to be insensitive to this choice.

Because dimensional sensitivities cannot be directly compared across parameters, dimensionless relative sensitivity coefficients were also computed, following the approach of (Sadegh Zadeh & Montas, 2010), (Kuznetsov & Kuznetsov, 2019):

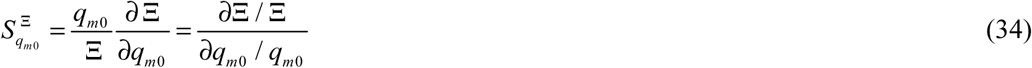

Eq. (34) shows that the dimensionless sensitivity 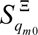 quantifies the fractional change in accumulated cytotoxicity induced by a fractional change in the corresponding parameter ( *q_m_*_0_ ). For example, a value of 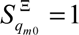 indicates that a 1% increase in the parameter *q_mo_* yields a corresponding 1% increase in accumulated cytotoxicity. It is important to distinguish between Ξ and its normalized sensitivity: accumulated cytotoxicity Ξ remains a dimensional quantity with units of concentration multiplied by time, whereas the relative sensitivity coefficient defined by Eq. (34) is dimensionless.

From Eqs. (9) and (31), it follows that

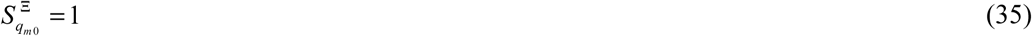

And

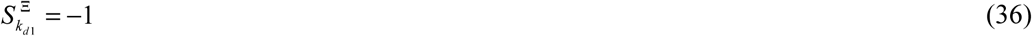

## 3. Results

### 3.1. Numerical implementation

The coupled system comprising Eqs. (1) and (4)–(6), subject to the initial conditions in Eq. (8), was integrated numerically in MATLAB R2024a using the built-in ODE45 solver. Both the relative and absolute error tolerances were set to 10^-10^. All figures in the main text (Figs. 2–11) and Supplementary Materials (Figs. S1–S39) were generated using the baseline parameter values listed in Table 2, except where an individual figure or its caption explicitly specifies alternative values. Additional numerical details are provided in Supplementary Section S1.

**Fig. 4.**
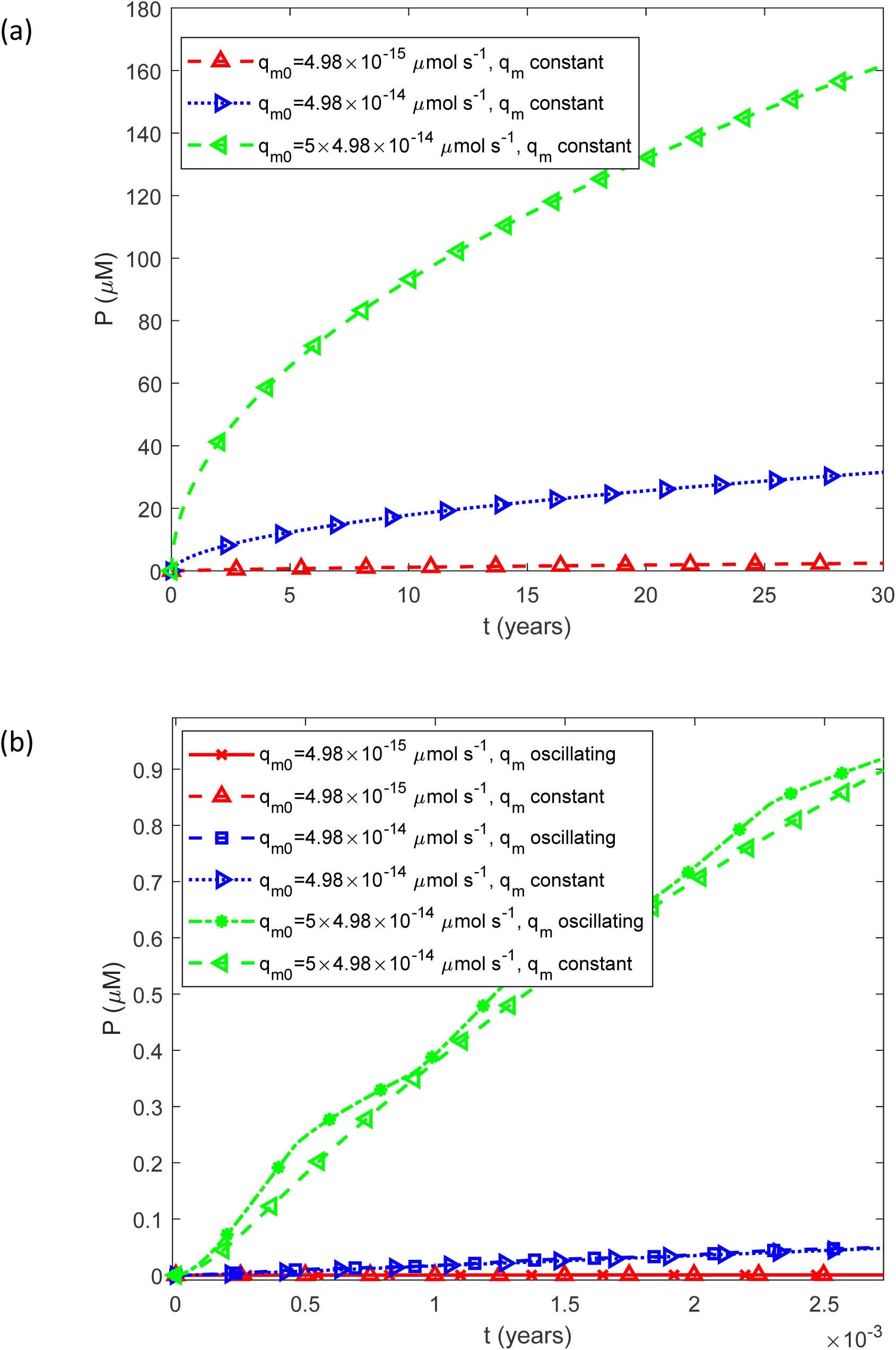
(a) Fibril particle concentration *P*(*t*) , i.e., the molar concentration of fibrils irrespective of length, as a function of time for a constant (time-averaged) monomer release rate of 1.25*q_m_* _0_ . (b) As in (a), but restricted to the time interval [0, 0.0027 years], equivalent to 1 day, illustrating the effect of oscillatory monomer release on the short-term dynamics of the fibril particle concentration. Results are shown for three representative values of the basal IAPP monomer secretion rate by beta cells, *q_m_* _0_ .

**Fig. 5.**
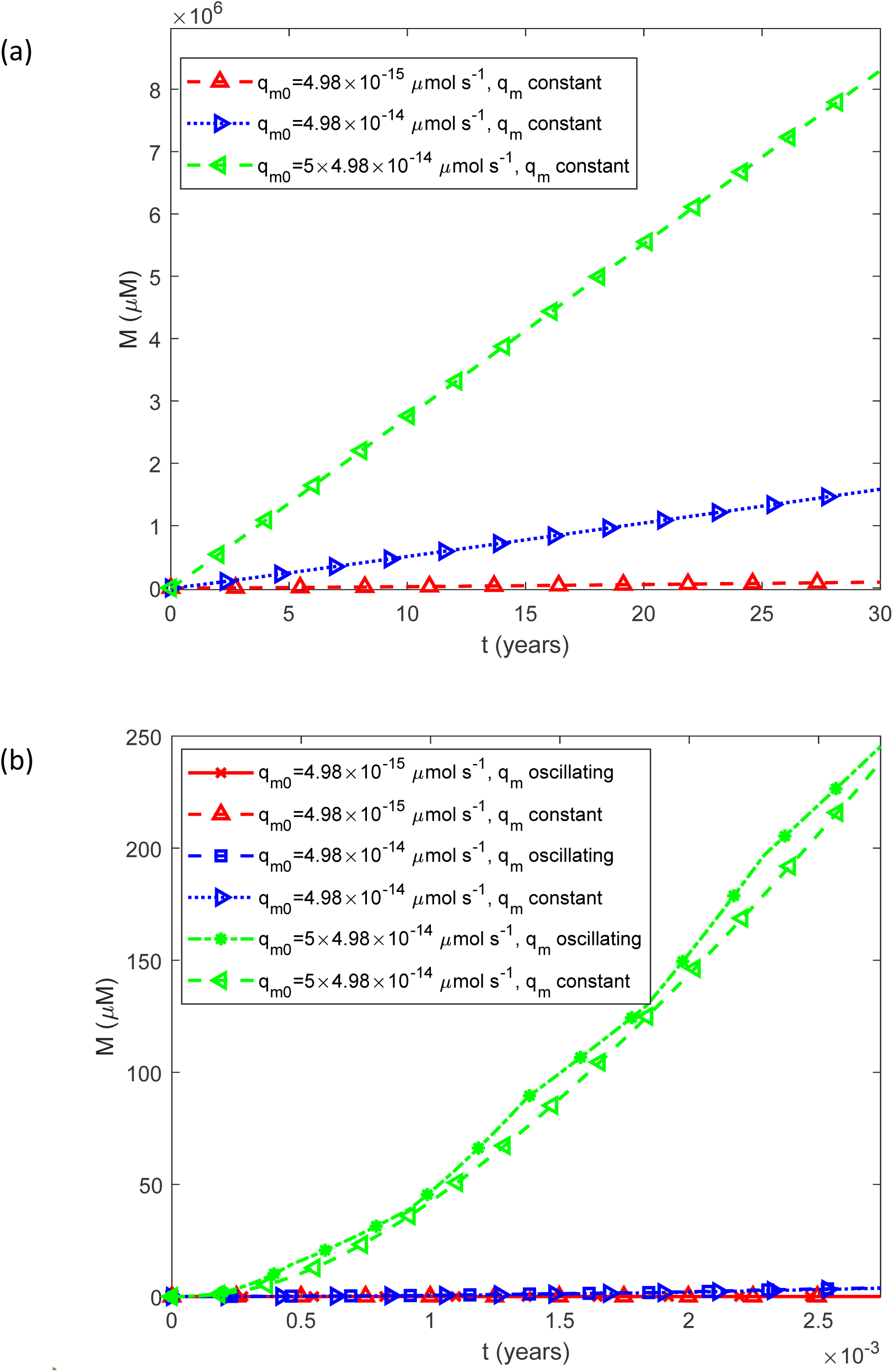
(a) Fibril mass concentration *M* (*t*) , expressed as the molar concentration of IAPP monomer equivalents incorporated into all fibrils, as a function of time for a constant (time-averaged) monomer release rate of 1.25*q_m_* _0_ . (b) As in (a), but restricted to the time interval [0, 0.0027 years], equivalent to 1 day, illustrating the effect of oscillatory monomer release on the short-term dynamics of total fibril mass. Results are shown for three representative values of the basal IAPP monomer secretion rate by beta cells, *q_m_* _0_ .

**Fig. 6.**
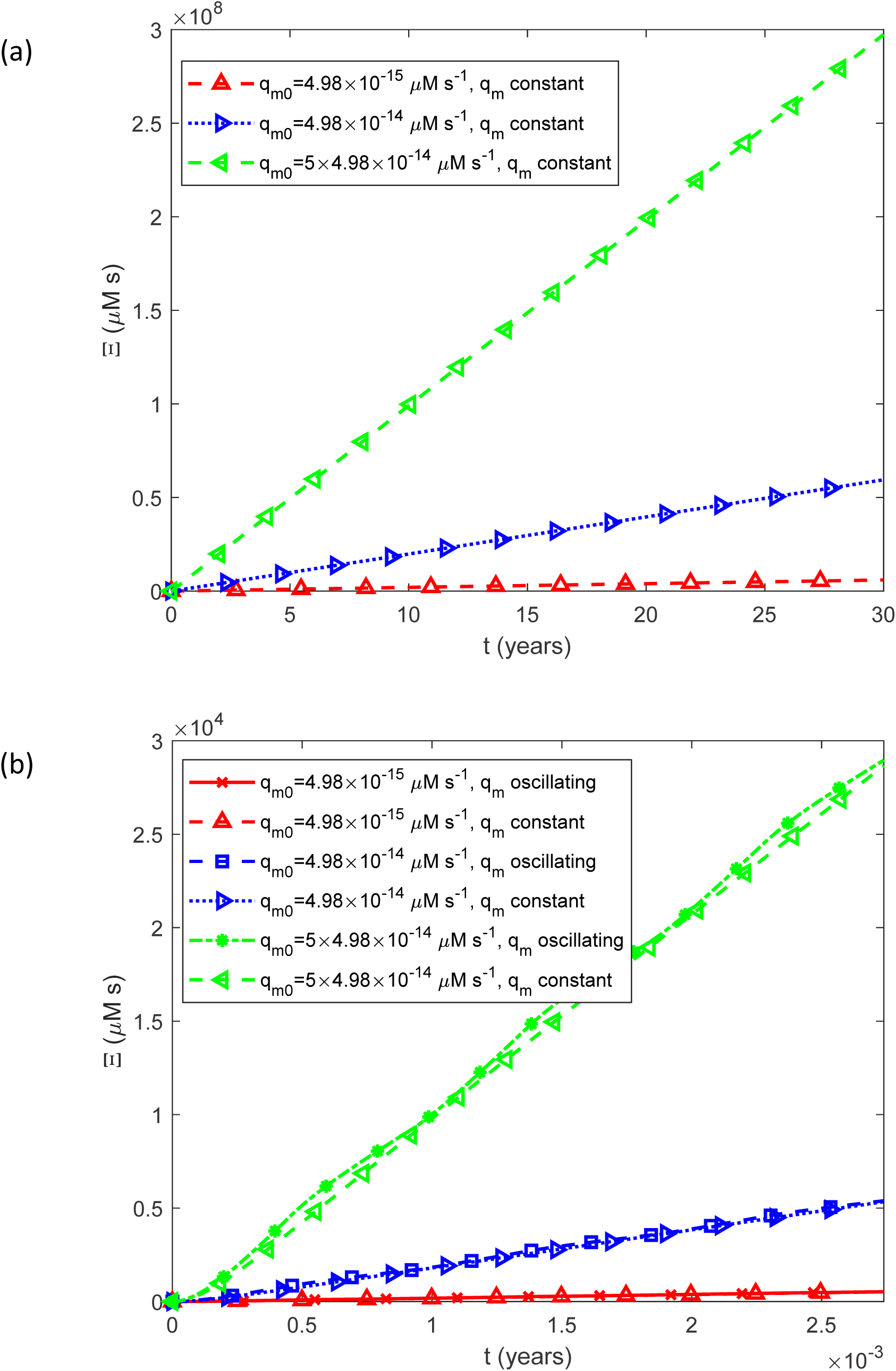
(a) Accumulated cytotoxicity (integrated IAPP oligomer exposure), Ξ(*t*) , as a function of time for a constant (time-averaged) monomer release rate of 1.25*q_m_* _0_ . (b) As in (a), but restricted to the time interval [0, 0.0027 years], equivalent to 1 day, illustrating the effect of oscillatory monomer release on the short-term dynamics of accumulated cytotoxicity. Results are shown for three representative values of the basal IAPP monomer secretion rate by beta cells, *q_m_*_0_ .

**Fig. 7.**
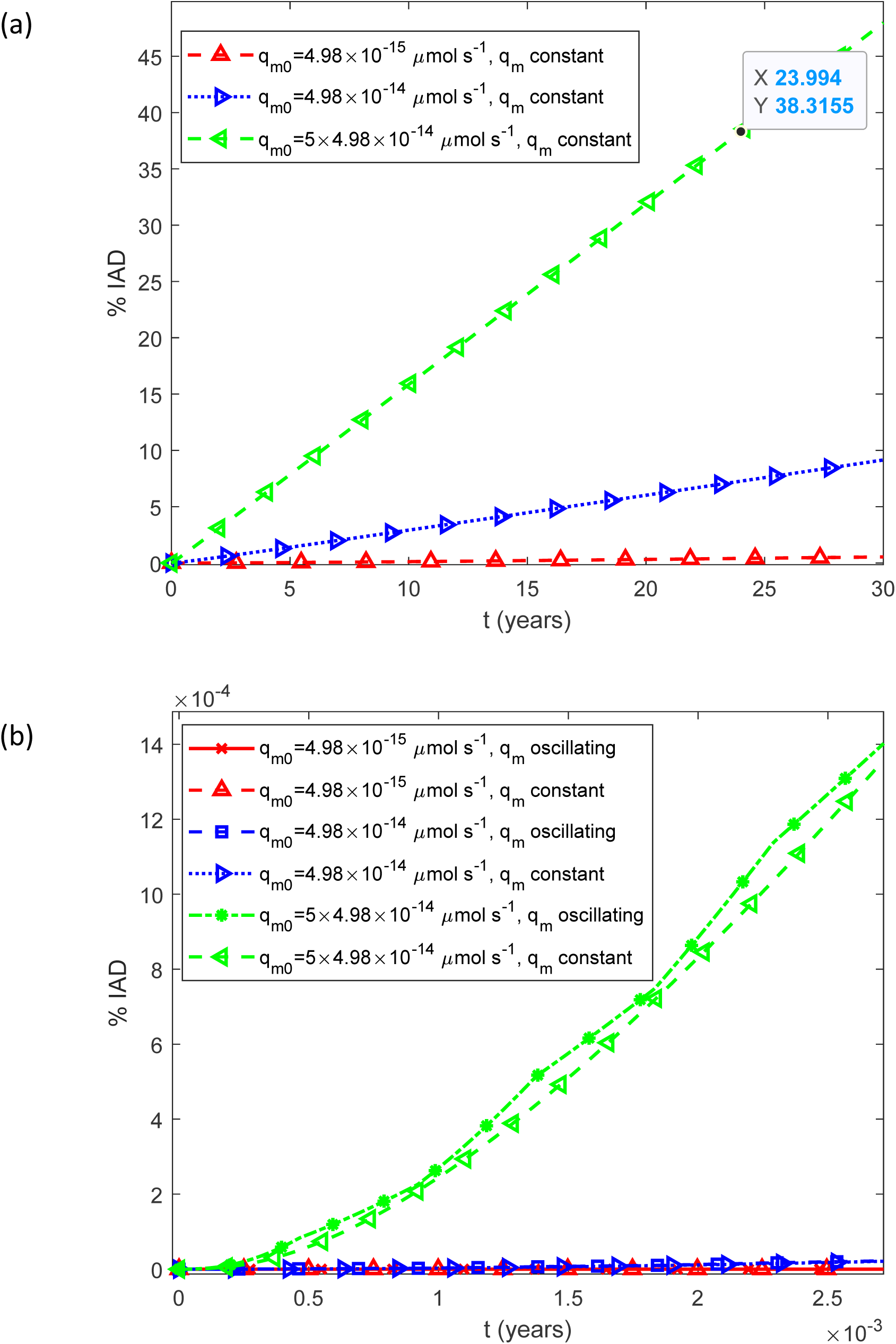
(a) Predicted percentage of the islet occupied by IADs, interpreted as the expected histological area fraction under the stereological assumption described in Section 2.2, as a function of time, %IADs, for a constant (time-averaged) monomer release rate of 1.25*q_m_* _0_ . (b) As in (a), but restricted to the time interval [0, 0.0027 years], equivalent to 1 day, illustrating the effect of oscillatory monomer release on the short-term dynamics of amyloid deposition. Results are shown for three representative values of the basal IAPP monomer secretion rate by beta cells, *q_m_* _0_ .

**Fig. 8.**
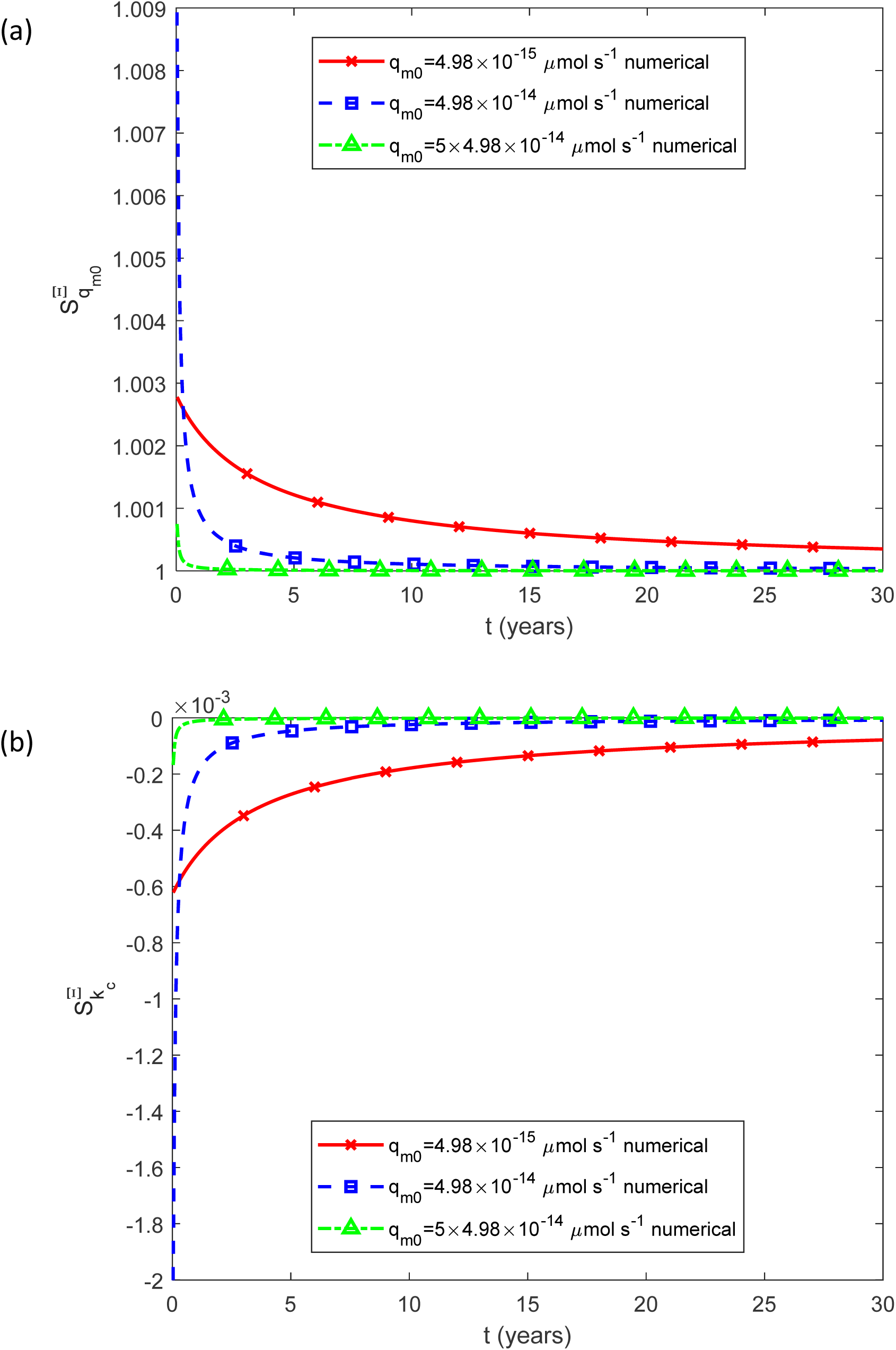
Dimensionless relative sensitivity coefficients of accumulated cytotoxicity with respect to (a) the basal rate of IAPP monomer secretion by beta cells, 1.25*q_m_*_0_ , and (b) the rate constant governing IAPP monomer clearance through fenestrated capillaries, *k_c_* , both shown as functions of time. Results are presented for three values of the basal monomer release rate.

**Fig. 9.**
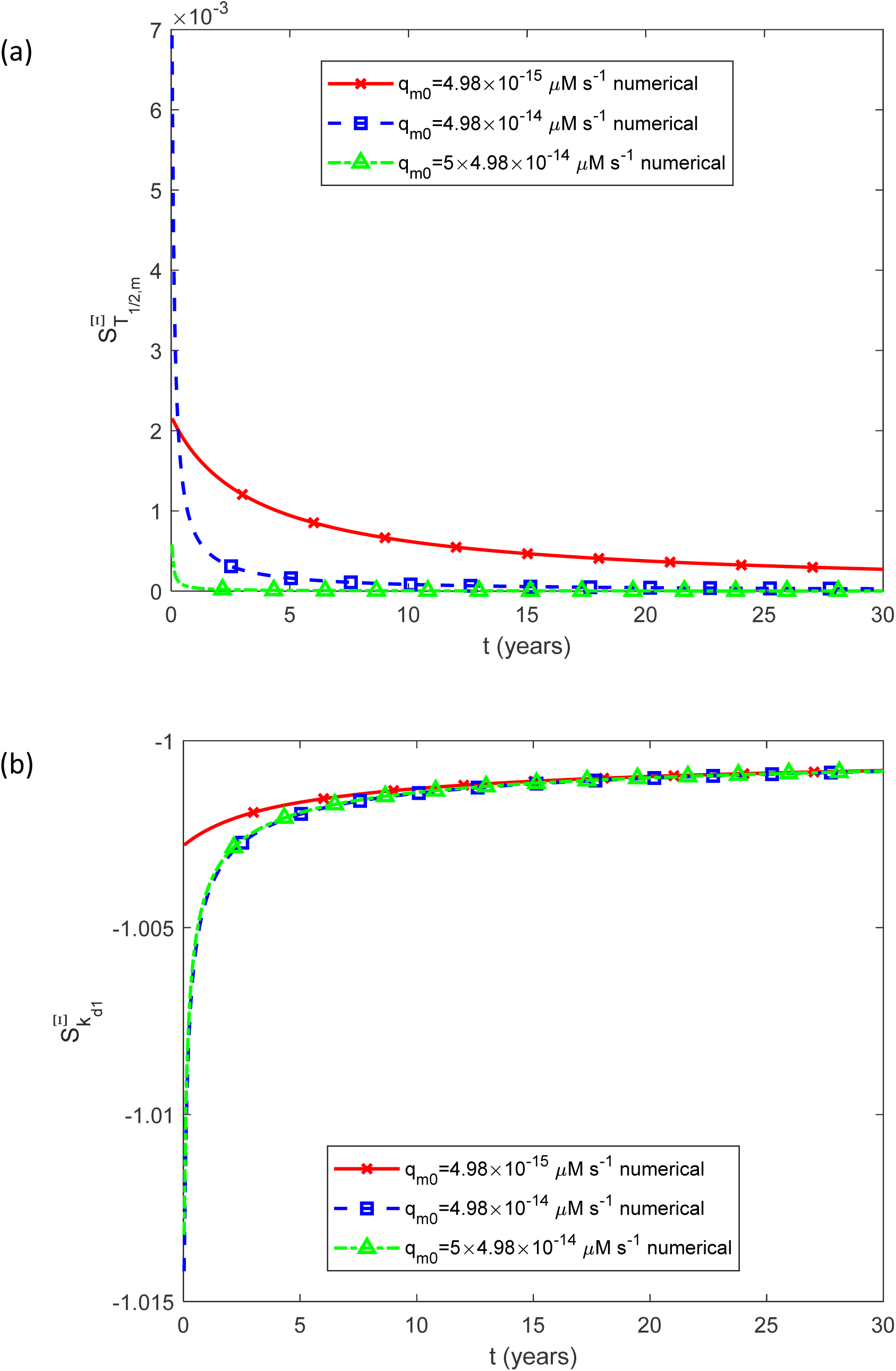
Dimensionless relative sensitivity coefficients of accumulated cytotoxicity with respect to (a) the half-life characterizing the enzymatic degradation of IAPP monomers, *T*_1/ 2,*m*_ , and (b) the rate constant for the spontaneous dissociation of IAPP oligomers into constituent monomers, *k_d_*_1_ , both shown as functions of time. Results are presented for three values of the basal monomer release rate.

**Fig. 10.**
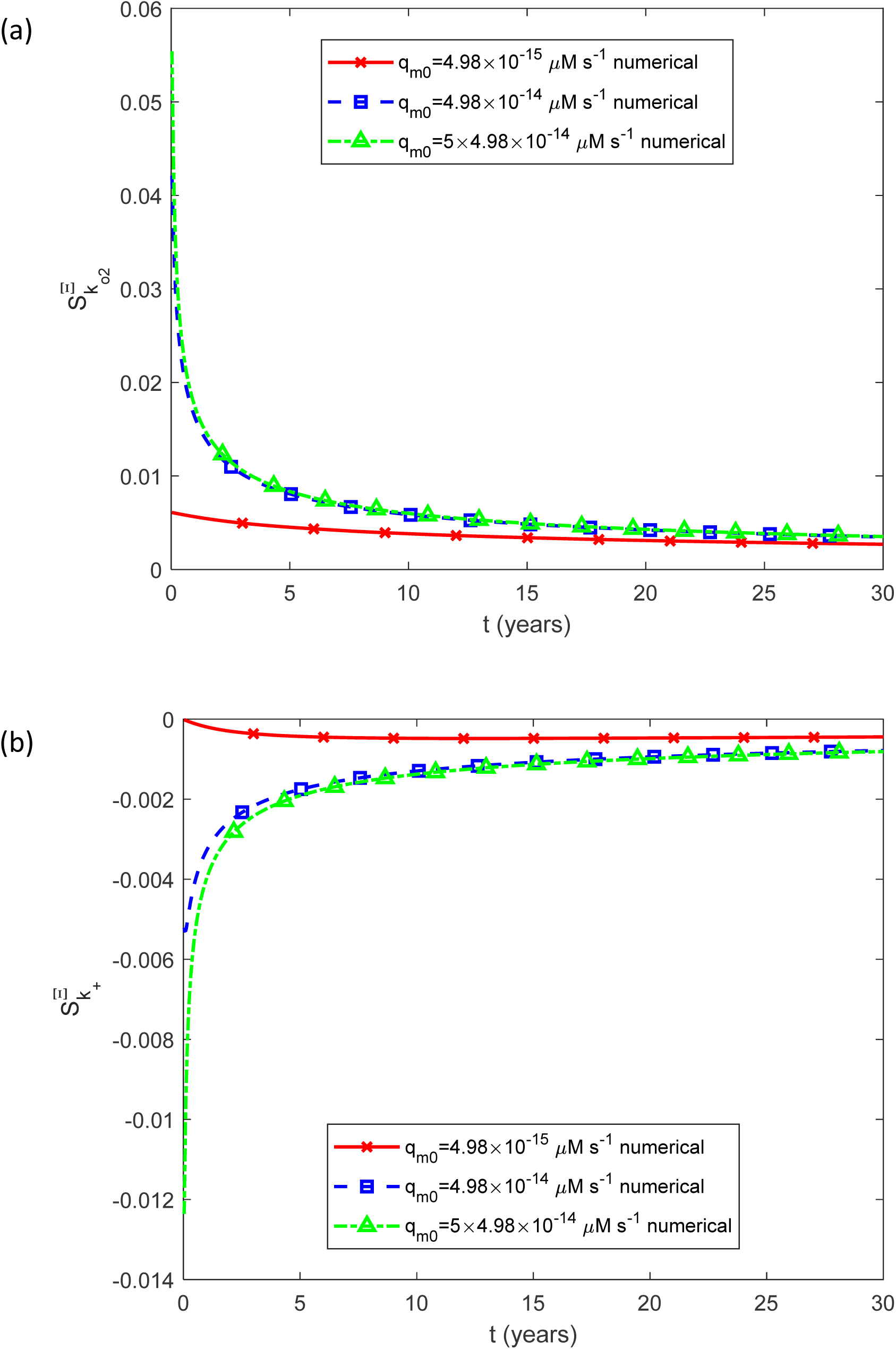
Dimensionless relative sensitivity coefficients of accumulated cytotoxicity with respect to (a) the rate constant for oligomer-surface-catalyzed monomer-to-oligomer conversion, *k_o_*_2_ , and (b) the rate constant for IAPP fibril elongation by monomer addition, *k*_+_ , both shown as functions of time. Results are presented for three values of the basal monomer release rate.

**Fig. 11.**
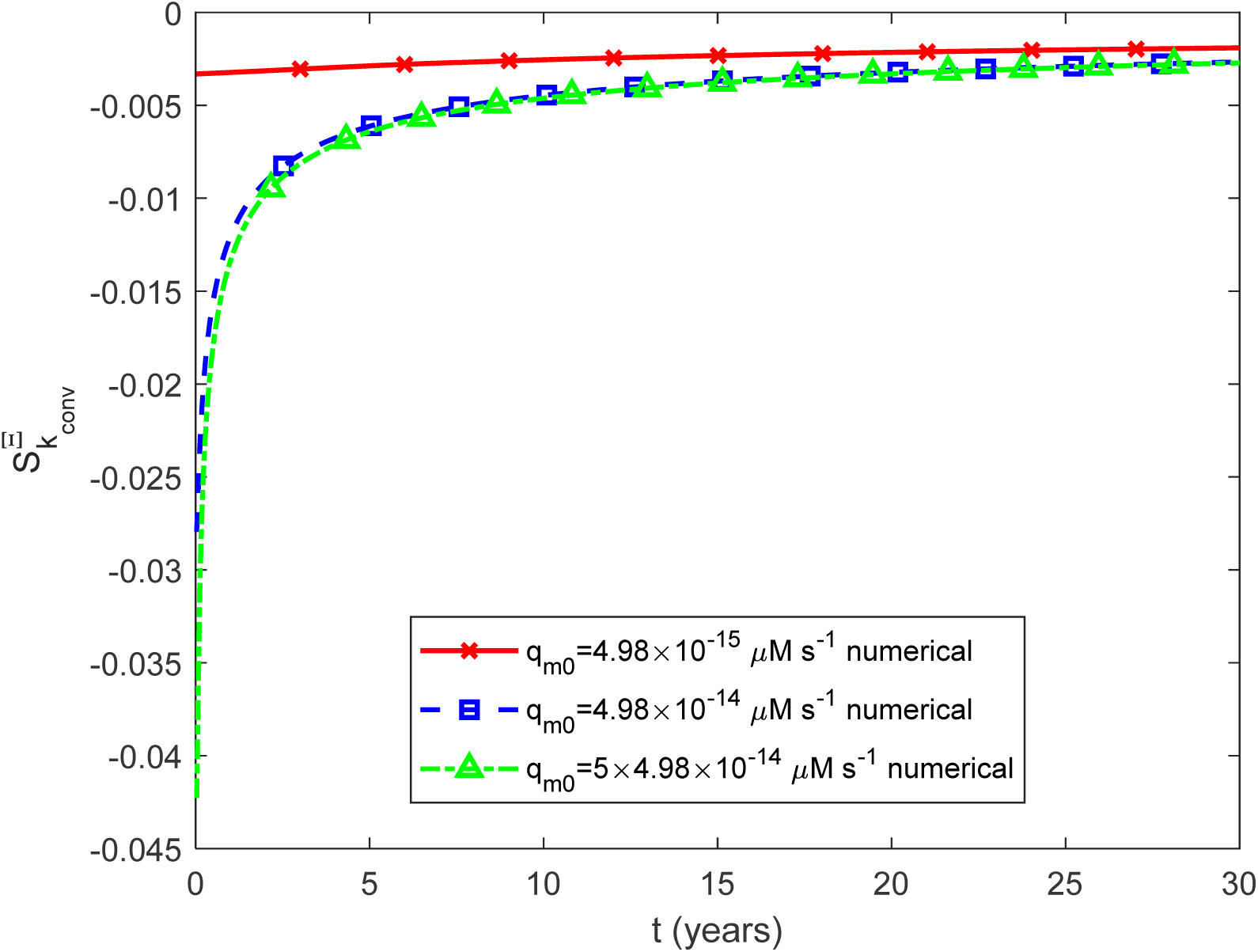
Dimensionless relative sensitivity coefficients of accumulated cytotoxicity with respect to the rate constant for the conversion of IAPP oligomers into elongation-competent fibrils, *k_conv_*, shown as a function of time. Results are presented for three values of the basal monomer release rate.

### 3.2. Baseline dynamics

The model captures the expected sequence of events in IAPP aggregation. The free monomer concentration drops rapidly from its initial value and remains low thereafter, as monomers are rapidly consumed by conversion into oligomers and fibrillar species (Fig. 2a). When monomer release is oscillatory, the monomer concentration faithfully mirrors the square-wave input, reflecting the short timescale on which monomers are turned over. A higher monomer secretion rate results in a proportionally higher monomer concentration (Fig. 2a,b).

The oligomer concentration rapidly reaches a steady state that is maintained thereafter, in close agreement with the analytical solution given by Eq. (28). The steady-state value scales linearly with the monomer secretion rate (Fig. 3a). Under oscillatory release, oligomers respond gradually rather than abruptly, rising slowly when the secretion rate increases and declining slowly when it falls (Fig. 3b), reflecting the finite timescale of oligomer turnover.

The fibril particle concentration *P* (*t* ) , defined as the molar concentration of fibrils irrespective of their length, increases as *t*^1/2^ , consistent with Eq. (29) and Fig. 4a. In contrast, *M* (*t* ) , the concentration of IAPP monomer equivalents incorporated into all fibrils and therefore a measure of total fibril mass per unit volume, increases approximately linearly with time, consistent with the leading-order analytical result in Eq. (16) and Fig. 5a. Both quantities exhibit only small-amplitude oscillations under pulsatile monomer release (Figs. 4b, 5b), indicating that the fibrillar pool is relatively insensitive to short-term fluctuations in input. The accumulated cytotoxic burden associated with IAPP oligomers likewise increases linearly with time (Fig. 6a), a direct consequence of the near-constant oligomer concentration whose integral is linear. Higher monomer secretion rates amplify this burden proportionally, while oscillatory release produces negligible oscillations in cytotoxicity (Fig. 6b).

The fraction of the islet occupied by amyloid deposits increases linearly with time (Fig. 7a), and a higher monomer secretion rate results in a proportionally greater fraction of the islet area replaced by amyloid. The model predicts approximately 48% replacement of the islet area after 30 years at the highest secretion rate, which is lower than the more than 85% replacement reported in end-stage disease (Jaikaran & Clark, 2001). This gap reflects the fact that end-stage disease is likely to involve additional pathological processes including beta-cell death and release of intracellular IAPP, not captured in the baseline model. Oscillatory release produces only minor fluctuations in amyloid burden (Fig. 7b). For the limiting case of infinitely long monomer and oligomer half-lives, the numerical results agree well with the analytical solutions of Eqs. (16) and (27)–(31), providing a check on the numerical solution (Figs. S1–S3). The agreement between the numerical and analytical results constitutes an internal verification of the numerical implementation; it should not be interpreted as an independent validation of the biological model.

### 3.3. The effect of capillary clearance

The capillary clearance rate constant exerts a selective and physically interpretable influence on the system. A higher clearance rate reduces the steady-state monomer concentration by accelerating removal of monomers from the interstitial space (Fig. S4), and consequently attenuates the accumulation of fibrillar species of all lengths (Fig. S6), total fibril mass (Fig. S7), and the fraction of the islet occupied by amyloid deposits (Fig. S9). In contrast, the clearance rate constant has no significant effect on oligomer concentration (Fig. S5) or accumulated cytotoxicity (Fig. S8). This selective insensitivity arises because IAPP oligomers exceed the size threshold of the capillary fenestrations and are therefore excluded from this clearance pathway altogether, a structural feature of the islet microvasculature with important implications for therapeutic strategies aimed at enhancing capillary drainage.

### 3.4. The effect of enzymatic degradation

The enzymatic degradation half-life of IAPP monomers influences the CV through a common mechanism: a longer half-life slows proteolytic removal, allowing monomers to persist in the interstitial space for longer and increasing the proportion that undergo conversion into oligomers and fibrillar species. Accordingly, both monomer concentration (Fig. S10) and, to a modest extent, oligomer concentration (Fig. S11) increase with the degradation half-life. The concentrations of fibrillar species of all lengths (Fig. S12), total fibril mass (Fig. S13), and the fraction of the islet occupied by amyloid deposits (Fig. S15) all increase correspondingly. The effect on accumulated cytotoxicity is very small (Fig. S14), reflecting the modest nature of the oligomer concentration increase, an effect sufficiently small to become negligible on longer timescales. Taken together, these results identify enzymatic degradation as a secondary process whose influence is real but quantitatively subordinate to monomer secretion and capillary clearance.

### 3.5. The effect of oligomer dissociation

The rate constant for spontaneous oligomer dissociation into monomers produces effects that are, in several respects, the reverse of those described above. A higher dissociation rate constant increases monomer concentration, since dissociating oligomers return monomers to the interstitial pool (Fig. S16), while simultaneously reducing oligomer concentration (Fig. S17). The downstream consequences of reduced oligomer availability propagate through the aggregation cascade: fibrillar species of all lengths (Fig. S18), total fibril mass (Fig. S19), accumulated cytotoxicity (Fig. S20), and the fraction of the islet occupied by amyloid deposits (Fig. S21) all decrease with increasing dissociation rate. This parameter therefore plays an important role in the model, as it directly controls the size of the oligomeric pool and, through it, both the cytotoxic burden and the rate of amyloid deposition.

### 3.6. The effect of oligomer-surface-catalyzed conversion and fibril elongation

The rate constant for oligomer-surface-catalyzed monomer-to-oligomer conversion increases the oligomer concentration (Fig. S23) while producing only a negligible decrease in monomer concentration (Fig. S22), since monomers are continuously replenished by beta-cell secretion. The enlarged oligomeric pool in turn drives greater fibrillar accumulation (Figs. S24, S25) and accumulated cytotoxicity (Fig. S26), as well as a larger amyloid deposit fraction at short timescales (Fig. S27). These effects progressively diminish at longer timescales, however, suggesting that the monomer-to-oligomer conversion rate constant plays a diminishing role at long times, where the dynamics are governed primarily by monomer production and oligomer dissociation.

The fibril elongation rate constant acts through a different mechanism. Higher elongation rates consume monomers more rapidly (Fig. S28), reducing monomer concentration while leaving the oligomer concentration unaffected (Fig. S29), since oligomers are not directly involved in elongation. The concentration of fibrillar species decreases with increasing elongation rate constant (Fig. S30) because, in the model, it is directly proportional to the monomer concentration; however, the total fibril mass increases (Fig. S31), since each elongation event adds mass to existing fibrils. Accumulated cytotoxicity is insensitive to the elongation rate constant (Fig. S32), consistent with the insensitivity of the oligomer concentration, while the fraction of amyloid deposits increases with increasing elongation rate (Fig. S33).

An increase in the rate constant for oligomer-to-fibril conversion simultaneously depletes both the monomer pool (Fig. S34) and the oligomer pool (Fig. S35), resulting in increased concentrations of fibrillar species (Fig. S36) and total fibril mass (Fig. S37). Accumulated cytotoxicity decreases with this rate constant at short timescales (Fig. S38), as a consequence of oligomer depletion, though this effect vanishes at longer times. The amyloid deposit fraction increases correspondingly (Fig. S39).

### 3.7. Sensitivity analysis of accumulated cytotoxicity

The dimensionless relative sensitivity coefficients reveal a clear hierarchy among the model parameters in their influence on accumulated cytotoxicity. The basal monomer secretion rate is the dominant driver: its dimensionless sensitivity approaches unity in the long-time limit (Fig. 8a), implying that a 1% increase in secretion rate produces a 1% increase in cumulative cytotoxicity. At the opposite extreme, the rate constant for spontaneous oligomer dissociation approaches a limiting sensitivity of −1 (Fig. 9b), meaning that a 1% increase in dissociation rate produces a 1% decrease in cumulative cytotoxicity, confirming that this parameter is an equally potent but opposing modulator. All other parameters, including the capillary clearance rate constant (Fig. 8b), enzymatic degradation half-life (Fig. 9a), oligomer-surface-catalyzed conversion rate constant (Fig. 10a), fibril elongation rate constant (Fig. 10b), and oligomer-to-fibril conversion rate constant (Fig. 11), have dimensionless sensitivities that remain well below unity throughout and approach zero asymptotically. This hierarchy has a clear therapeutic implication: interventions targeting monomer secretion or oligomer dissociation are predicted to have the greatest impact on the long-term cytotoxic burden, while those targeting downstream aggregation kinetics are expected to have comparatively modest effects.

## 4. Discussion, limitations, and future directions

The central aim of this study was to develop a mathematical model of IAPP aggregation in the islets of Langerhans that is based on the physiology of beta cells and the microanatomy of the islet, and to use it to identify the parameters that most strongly influence the long-term cytotoxic burden and amyloid deposition in type 2 diabetes. The results reveal a clear and therapeutically significant hierarchy among the model parameters, with potential implications for the design of intervention strategies.

Following the hypothesis that oligomeric intermediates are the most cytotoxic IAPP species (Haataja *et al*., 2008), (Raleigh *et al*., 2017), (Gurlo *et al*., 2010), the concept of accumulated cytotoxicity is introduced, defined as the time integral of the IAPP oligomer concentration, which quantifies the cumulative damage inflicted by IAPP oligomers on pancreatic beta cells. A sensitivity analysis of accumulated cytotoxicity with respect to the model parameters is then performed.

The most significant finding of the sensitivity analysis is the dominant influence of two parameters in determining the long-term accumulated cytotoxicity: the basal rate of IAPP monomer secretion by beta cells, and the rate constant for spontaneous oligomer dissociation. Both approach a dimensionless sensitivity of magnitude unity in the long-time limit, the former positive and the latter negative, implying that a 1% increase in either parameter produces an approximately 1% change in cumulative cytotoxicity, with the sign determined by the parameter. All other parameters, including the capillary clearance rate constant, the enzymatic degradation half-life, the fibril elongation rate constant, and the oligomer-to-fibril conversion rate constant, have sensitivities that remain well below unity and diminish toward zero at long times. This hierarchy reflects a fundamental feature of the model: in the long-time limit, the oligomer concentration, and therefore cytotoxicity, is governed primarily by the balance between monomer production and the competing pathways of oligomer formation and dissociation, while downstream kinetic processes play a secondary role.

The dominant influence of the monomer secretion rate is consistent with the well-established observation that beta-cell secretory demand is a primary risk factor for islet amyloidosis. Conditions that chronically increase insulin and IAPP secretion, including obesity, insulin resistance, and glucotoxicity, are strongly associated with islet amyloid formation and beta-cell loss in both human and animal studies (Hull *et al*., 2004), (Hayden *et al*., 2007). The model provides a quantitative framework for explaining why increased secretion leads to a higher steady-state monomer concentration, which in turn drives oligomer formation and, through it, accumulated cytotoxicity in direct proportion.

The equally important but opposing role of oligomer dissociation is a novel finding of the model with direct therapeutic implications. The model predicts that enhancing the rate at which oligomers spontaneously disassemble into monomers, returning IAPP to its soluble, non-toxic form, reduces cumulative cytotoxicity with a sensitivity comparable to that of reducing monomer secretion. This suggests that small molecules or peptides that promote oligomer dissociation without promoting fibril formation could represent a potentially effective therapeutic strategy, complementary to approaches that target monomer production.

An interesting mechanistic finding is the selective influence of capillary clearance on the system. Higher clearance rates reduce monomer concentration and, consequently, fibril accumulation and amyloid deposition, but leave the oligomer concentration and accumulated cytotoxicity essentially unaffected. This selectivity arises from the physical exclusion of IAPP oligomers from the fenestrated capillaries. To the author’s knowledge, this structural feature of the islet microvasculature has not previously been incorporated into mathematical models of islet amyloidosis. The implication is that the progressive microvascular remodeling that occurs in type 2 diabetes, including endothelial thickening, reduced fenestration, and increased extracellular matrix deposition (Aplin *et al*., 2024), would be expected, according to the model, to increase fibril accumulation and amyloid deposition, but would have no protective effect on the oligomer-mediated cytotoxic burden that presumably drives beta-cell apoptosis. This decoupling between amyloid deposition and accumulated cytotoxicity is an important conceptual result that has implications for the interpretation of histological studies, which typically measure amyloid area fraction as a surrogate for disease severity (Clark & Nilsson, 2004). The present model suggests that amyloid area fraction and accumulated cytotoxicity are not equivalent measures and may respond differently to therapeutic interventions.

Sensitivity analysis of accumulated cytotoxicity with respect to the oligomer-to-fibril conversion rate constant reveals a therapeutically important trade-off. According to the model, accelerating this conversion rate constant reduces the oligomeric pool and therefore transiently reduces cytotoxicity at short timescales, but simultaneously accelerates fibril accumulation and amyloid deposition. This trade-off reflects the double-edged nature of therapeutic strategies that seek to divert IAPP away from toxic oligomeric conformations toward relatively inert fibrillar deposits. Such an approach is supported by the autophagic sequestration mechanism described in (Rivera *et al*., 2014), in which the adaptor protein p62 channels aggregation-prone IAPP into relatively inert fibrils, thereby bypassing the formation of toxic oligomeric intermediates. Another potentially promising therapeutic strategy is the rational design of small molecules or peptide inhibitors that selectively promote the formation of non-toxic oligomeric conformations at the expense of toxic ones, thereby potentially diverting the aggregation pathway away from cytotoxicity (Saravanan *et al*., 2019). The present model provides a quantitative framework for evaluating the net benefit of such therapeutic strategies.

The model predicts that approximately 48% of the islet is occupied by amyloid deposits after 30 years at the highest monomer secretion rate, compared with more than 85% reported in some histological observations of end-stage type 2 diabetes (Jaikaran & Clark, 2001). Thus, the present prediction should not be regarded as quantitative agreement with the end-stage histological value. Rather, the model captures substantial progressive amyloid deposition while underpredicting the extent of replacement of the islet area observed in advanced disease. This gap is plausible because the present model omits several late-stage processes that can amplify amyloid accumulation, particularly beta-cell death and the associated release of intracellular IAPP, which can seed further extracellular fibril growth and generate a positive feedback between amyloid deposition and beta-cell loss (Westermark *et al*., 2000). The comparison should therefore be interpreted as qualitative or semiquantitative context rather than as independent validation of the model.

Several important limitations of the present model should be acknowledged. First, the model is formulated at the level of a single, idealized islet of Langerhans represented as a well-mixed compartment. This formulation neglects spatial heterogeneity of amyloid deposition within and between islets, which is well documented in histological studies (Ullsten *et al*., 2017), (Wang *et al*., 2023), as well as the non-uniform distribution of beta cells, capillaries, and extracellular matrix within the islet. A spatially resolved model, for example, a reaction–diffusion formulation, would be needed to capture these features.

Second, the model treats the beta-cell population as fixed and does not account for the progressive loss of beta cells that both results from and further amplifies IAPP aggregation. In reality, beta-cell apoptosis reduces the source of IAPP monomers over time while simultaneously releasing intracellular IAPP into the extracellular space (Westermark *et al*., 2011), thereby creating a complex positive feedback loop that is not captured in the present framework.

Third, several model parameters, most notably the oligomer-to-fibril conversion rate constant and the rate of preassembled oligomer release from dysfunctional secretory granules, are not directly measurable and must either be treated as free parameters or estimated indirectly. The sensitivity analysis demonstrates that the oligomer-to-fibril conversion rate constant has a diminishing influence on long-term outcomes, which partially mitigates this uncertainty, but the basal oligomer release rate from dysfunctional granules remains poorly constrained.

Fourth, the model adopts the Finke-Watzky framework (Morris *et al*., 2008), (Iashchishyn *et al*., 2017), in which an oligomer is treated as a single activated monomer. While this is a deliberate simplification that reduces the dimensionality of the model, it does not capture the stepwise assembly of oligomeric intermediates of varying size, a process known to give rise to a heterogeneous mixture of species with differing toxicities and aggregation propensities (Warren *et al*., 2026).

Several natural extensions of the present model would substantially enhance its predictive power and clinical relevance. Most importantly, incorporating beta-cell population dynamics, including oligomer-induced apoptosis and the consequent release of intracellular IAPP, would close the positive feedback loop between amyloid deposition and beta-cell loss that is central to the pathophysiology of type 2 diabetes. Such an extension would allow the model to predict the time course of beta-cell mass decline, which is a clinically measurable endpoint, and to identify parameter regimes in which the feedback loop becomes self-sustaining.

A second important extension would be the incorporation of spatial heterogeneity, by formulating the model as a system of reaction-diffusion equations within a realistic islet geometry. This would allow the model to capture the perivascular initiation of amyloid deposits and their centripetal growth, as well as the diffusion of IAPP monomers and oligomers through the interstitial space, processes that are inherently spatial and cannot be captured in a well-mixed framework.

Third, the model could be extended to incorporate the inhibitory effect of insulin on IAPP fibrillogenesis and the progressive loss of this protection due to impaired proinsulin processing in dysfunctional beta cells. This would allow the model to capture the synergistic roles of IAPP overproduction and insulin deficiency in promoting islet amyloidosis.

Fourth, the present model could serve as the basis for a pharmacokinetic–pharmacodynamic (PK–PD) framework for evaluating therapeutic interventions. The sensitivity analysis has already identified monomer secretion and oligomer dissociation as the two potentially most effective levers for reducing cytotoxicity; a PK-PD extension would allow the model to predict the dose-response relationship for candidate therapeutic agents targeting these processes.

## Author contributions

Andrey V. Kuznetsov is the sole author of this paper.

## Data availability

No additional data are associated with this article.

## Ethics Statement

Not applicable. This theoretical study did not involve human participants or animals.

## Abbreviations

Aβ: amyloid beta
AD: Alzheimer’s disease
CV: control volume (representing an interstitial space of an islet of Langerhans)
IADs: islet amyloid deposits
IAPP: islet amyloid polypeptide, also known as amylin
IDE: insulin-degrading enzyme

## Supplementary Materials

### S1. Numerical Solution

The coupled system of ordinary differential equations comprising Eqs. (1) and (4)–(6), subject to the initial conditions specified in Eq. (8), was integrated numerically in MATLAB (R2024a, MathWorks, Natick, MA, USA) using the built-in ODE45 solver. The ODE45 solver is based on the explicit Dormand–Prince Runge–Kutta scheme with adaptive step-size control and is well suited to nonstiff initial-value problems of the type considered here. Both the relative and absolute error tolerances (RelTol and AbsTol) were set to 10⁻¹⁰.

### S2. Supplementary figures

**Fig. S1.**
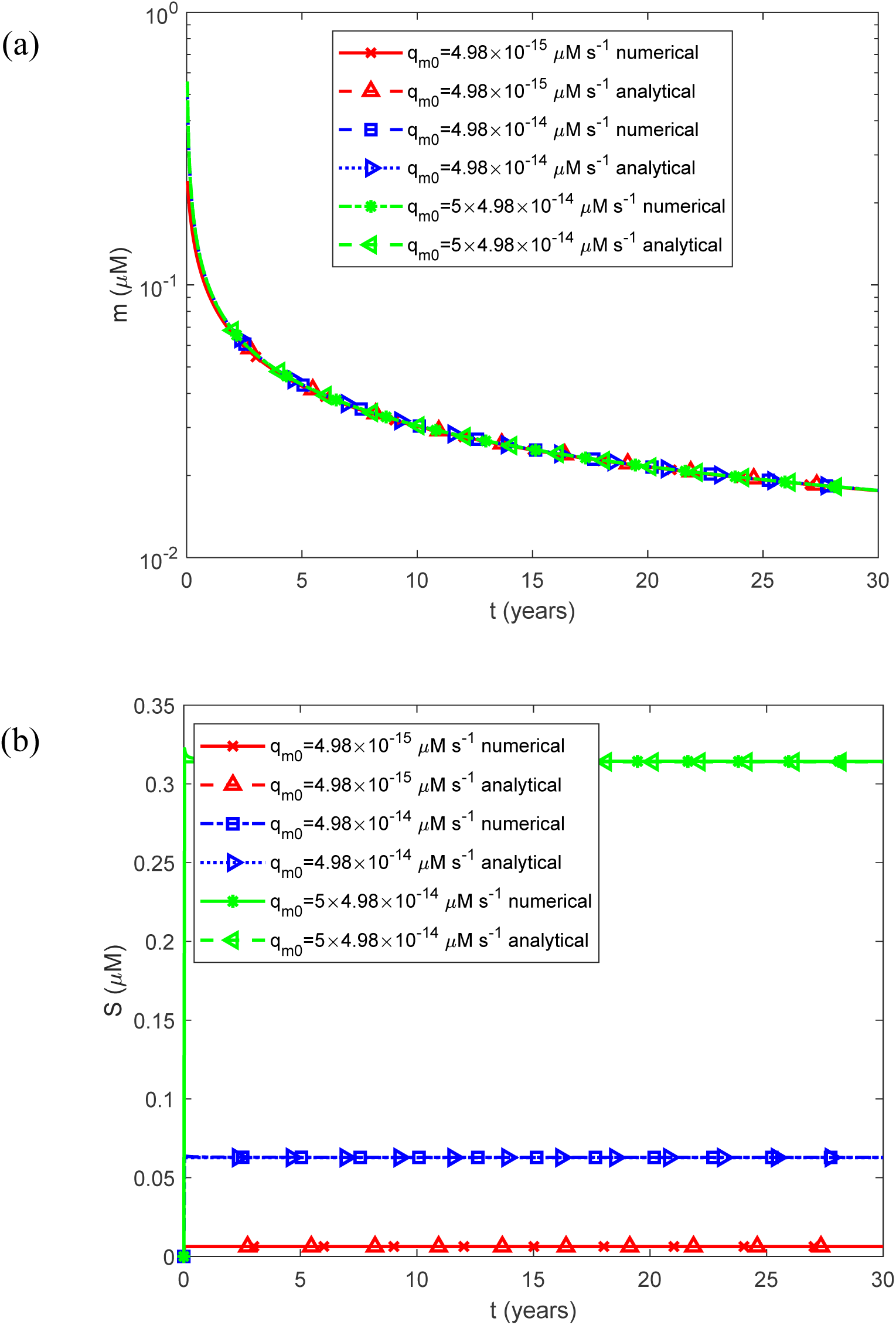
Comparison of the numerical and analytical solutions for the limiting case of infinitely long half-lives of IAPP monomers and oligomers, *T*_1/2,*m*_ →∞ and *T*_1/2,*S*_ →∞. (a) Molar concentration of free IAPP monomers as a function of time, *m*(*t*) . (b) Molar concentration of free IAPP oligomers as a function of time, *S*(*t*) . Results are shown for a constant (time-averaged) monomer release rate 1.25*q_m_*_0_ for three representative values of the basal rate at which IAPP monomers are secreted by beta cells, *q_m_*_0_ .

**Fig. S2.**
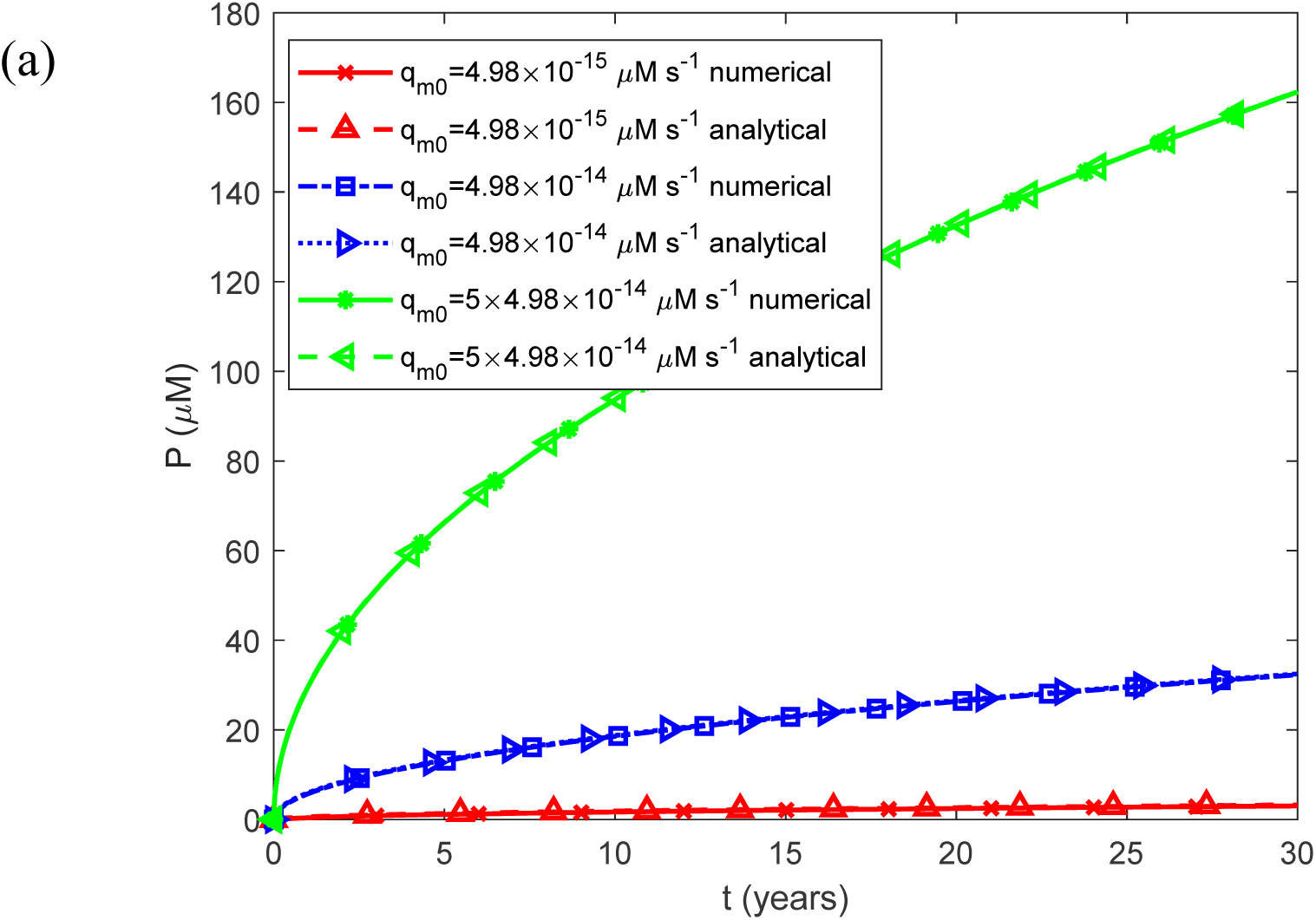
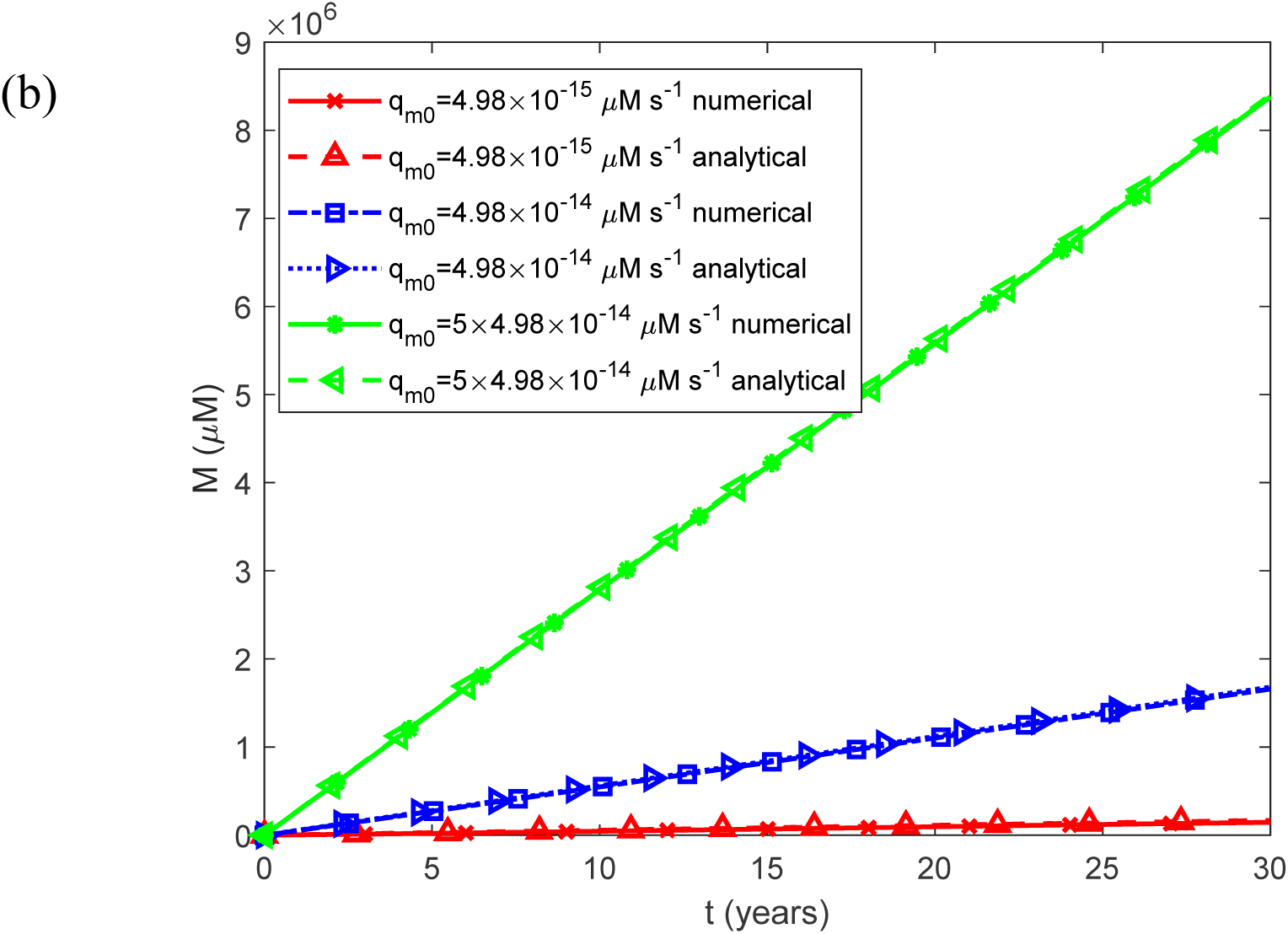
Comparison of the numerical and analytical solutions for the limiting case of infinitely long half-lives of IAPP monomers and oligomers, *T*_1/2,*m*_ →∞ and *T*_1/2,*S*_ →∞. (a) Fibril particle concentration *P*(*t*) , i.e., the molar concentration of fibrils irrespective of length, as a function of time. (b) Fibril mass concentration *M* (*t*) , expressed as the molar concentration of IAPP monomer equivalents incorporated into all fibrils, as a function of time. Results are shown for a constant (time-averaged) monomer release rate 1.25*q_m_*_0_ for three representative values of the basal rate at which IAPP monomers are secreted by beta cells, *q_m_*_0_ .

**Fig. S3.**
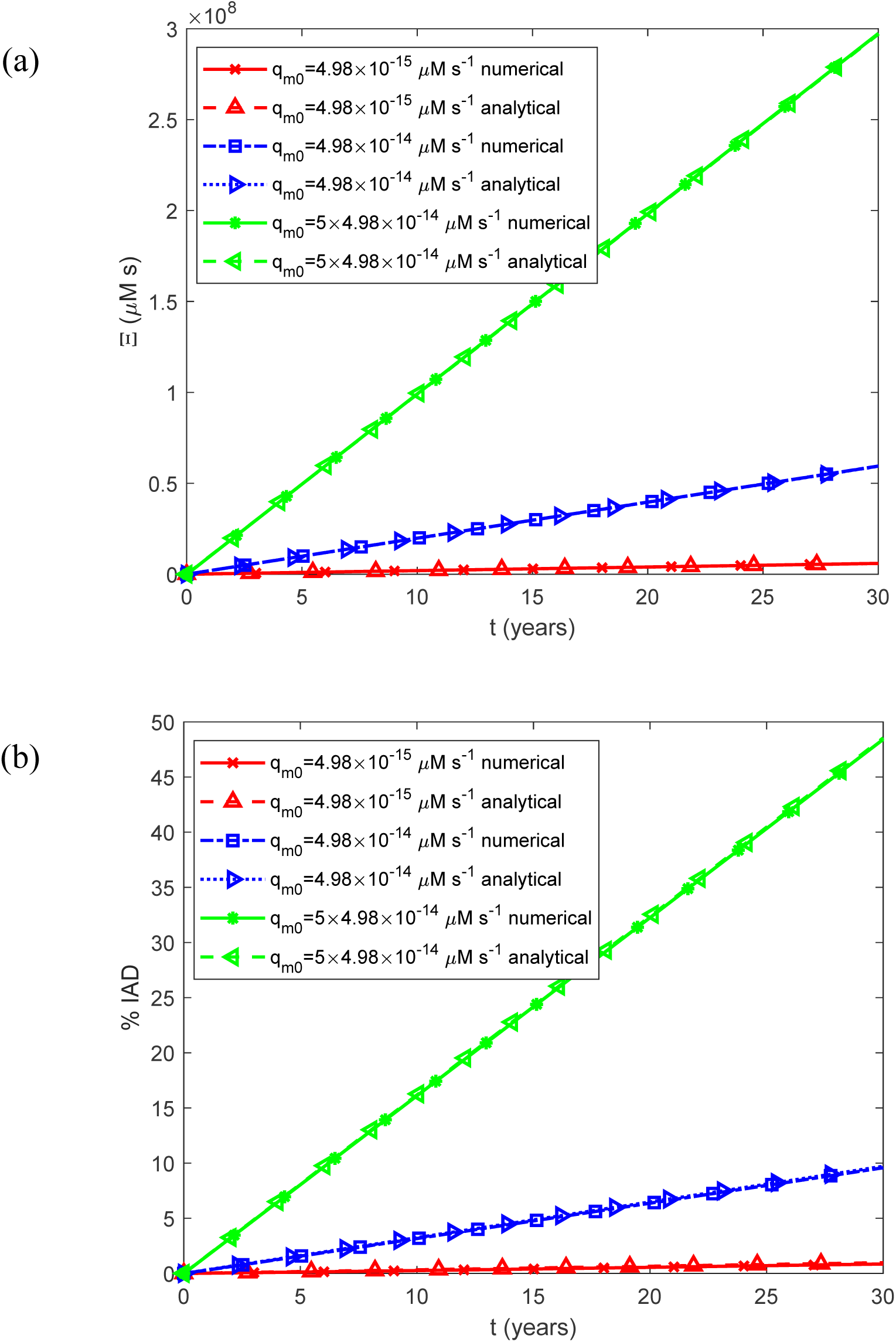
Comparison of the numerical and analytical solutions for the limiting case of infinitely long half-lives of IAPP monomers and oligomers, *T*_1/2,*m*_ →∞ and *T*_1/2,*S*_ →∞. (a) Accumulated cytotoxicity (integrated IAPP oligomer exposure) as a function of time, Ξ(*t*) . (b) Predicted percentage of the islet occupied by IADs, %IADs, interpreted as the expected histological area fraction under the stereological assumption described in Section 2.2, as a function of time. Results are shown for a constant (time-averaged) monomer release rate of 1.25*q_m_*_0_ monomers are secreted by beta cells, *q_m_*_0_ . for three representative values of the basal rate at which IAPP

**Fig. S4.**
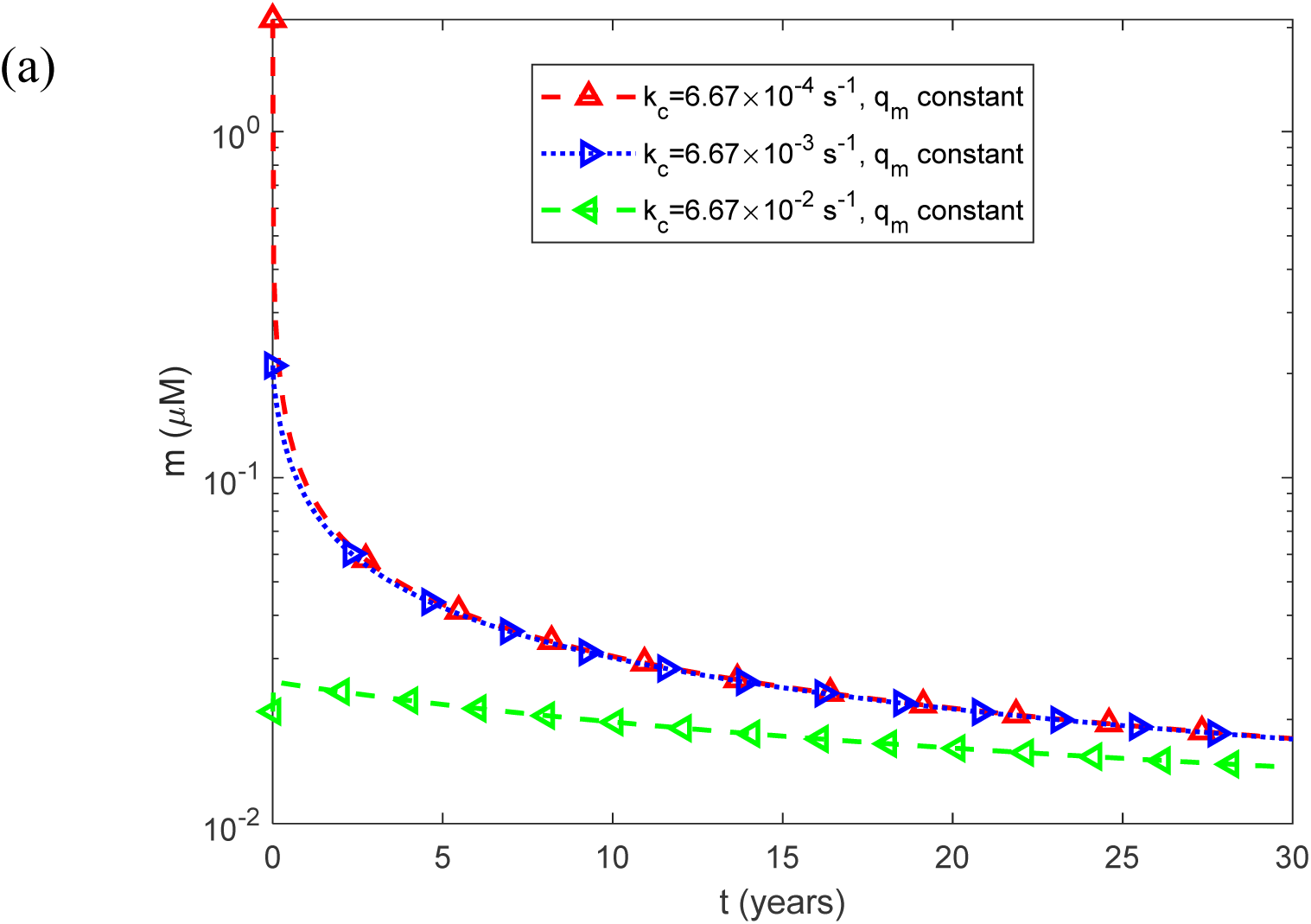
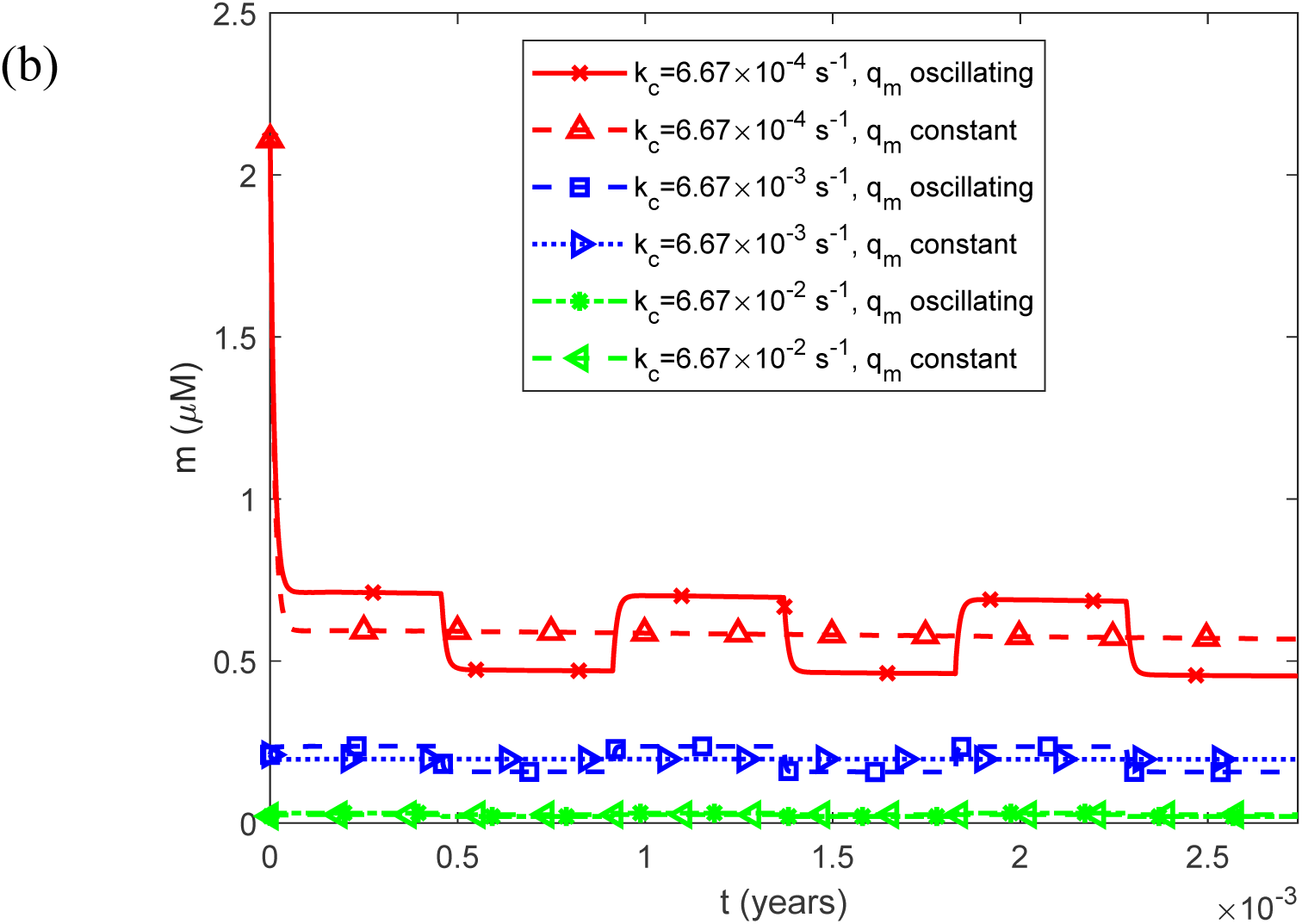
(a) Molar concentration of free IAPP monomers as a function of time, *m*(*t*) , for a constant (time-averaged) monomer release rate of 1.25*q_m_*_0_ . (b) As in (a), but restricted to the time interval [0, 0.0027 years], equivalent to 1 day, illustrating the effect of oscillatory monomer release on the short-term dynamics of the free monomer concentration. Results are shown for three representative values of the rate constant governing the clearance of IAPP monomers through fenestrated capillaries, *k_c_* .

**Fig. S5.**
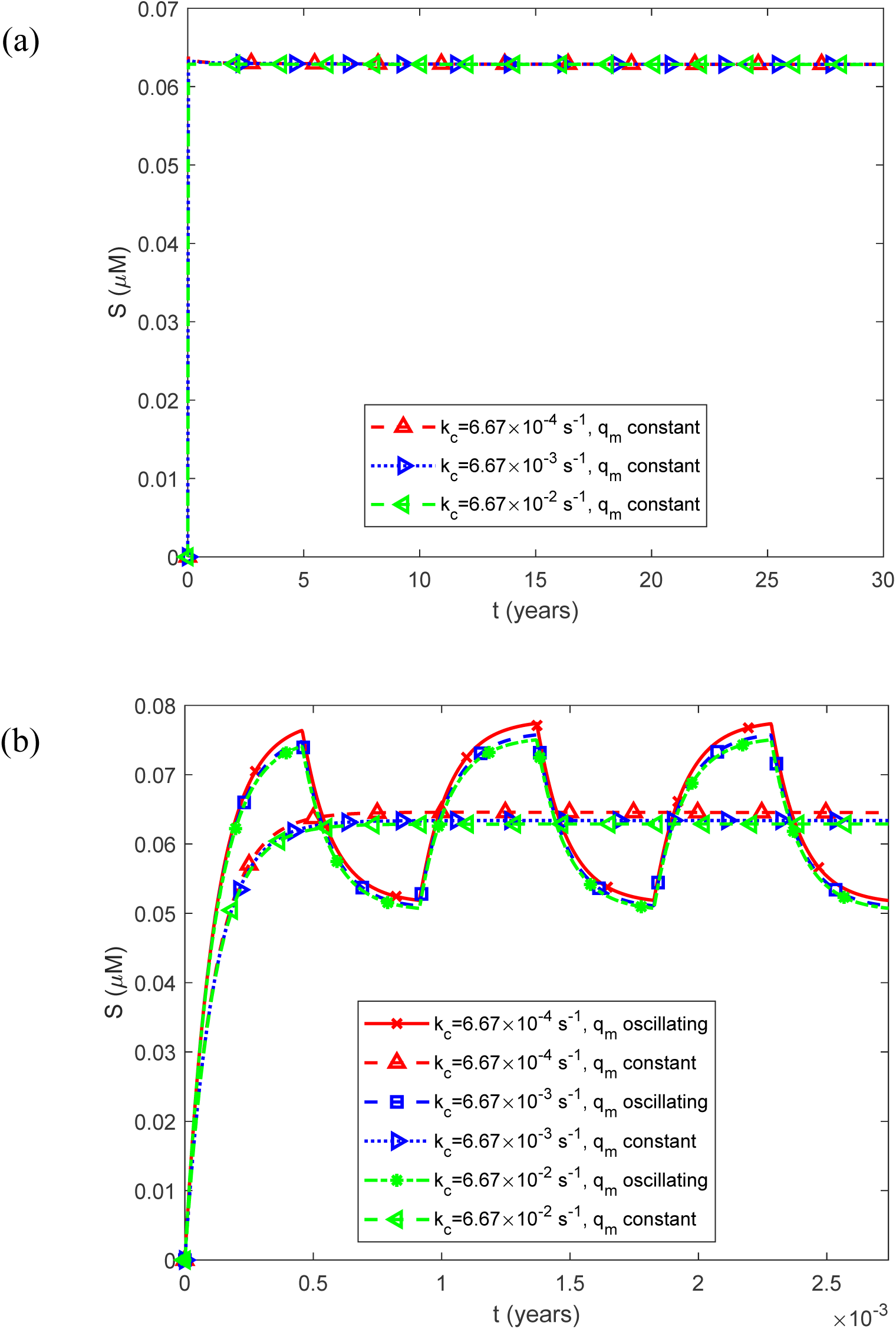
(a) Molar concentration of free IAPP oligomers as a function of time, *S*(*t*) , for a constant (time-averaged) monomer release rate of 1.25*q_m_*_0_ . (b) As in (a), but restricted to the time interval [0, 0.0027 years], equivalent to 1 day, illustrating the effect of oscillatory monomer release on the short-term dynamics of the free oligomer concentration. Results are shown for three representative values of the rate constant governing the clearance of IAPP monomers through fenestrated capillaries, *k_c_* .

**Fig. S6.**
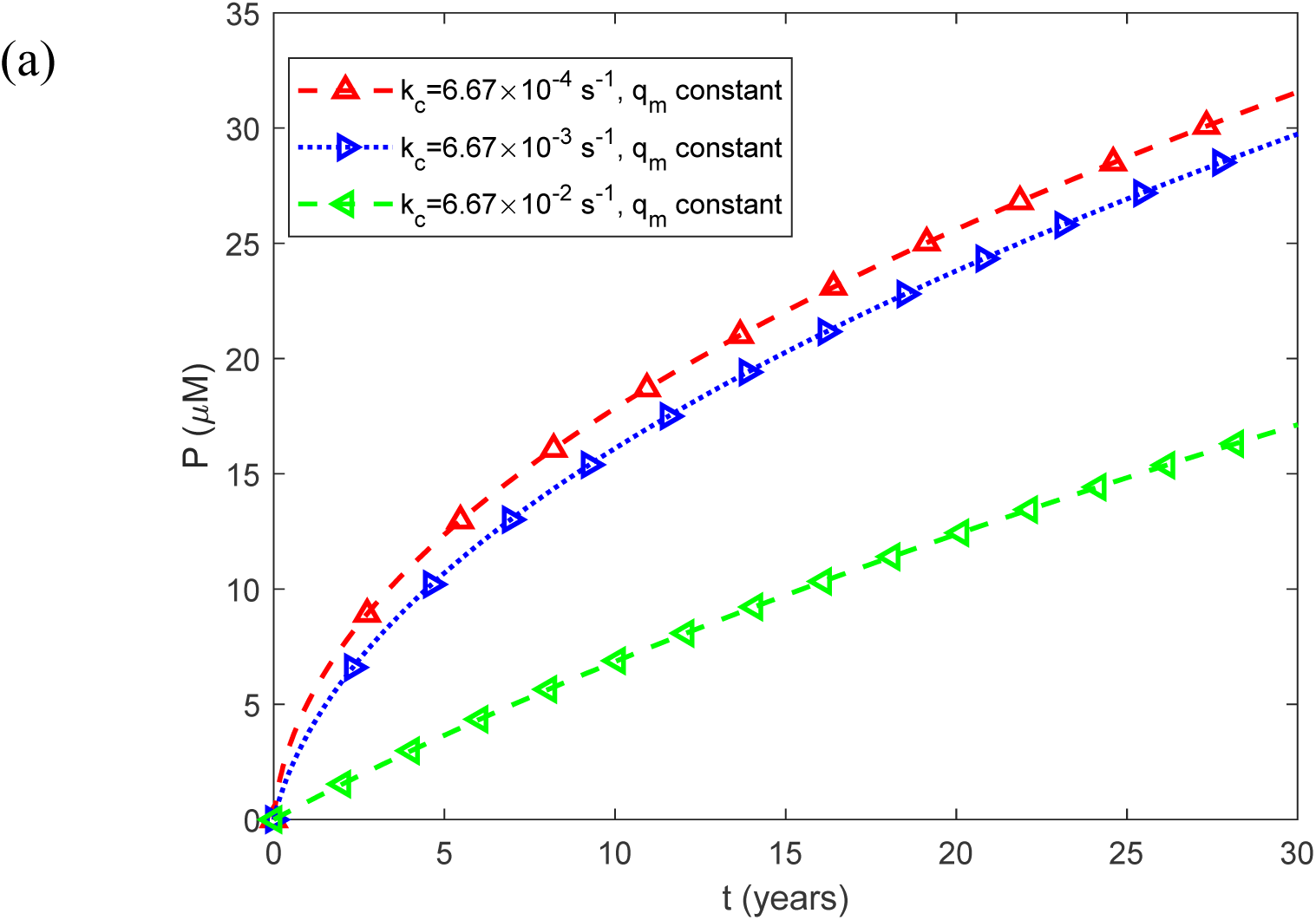
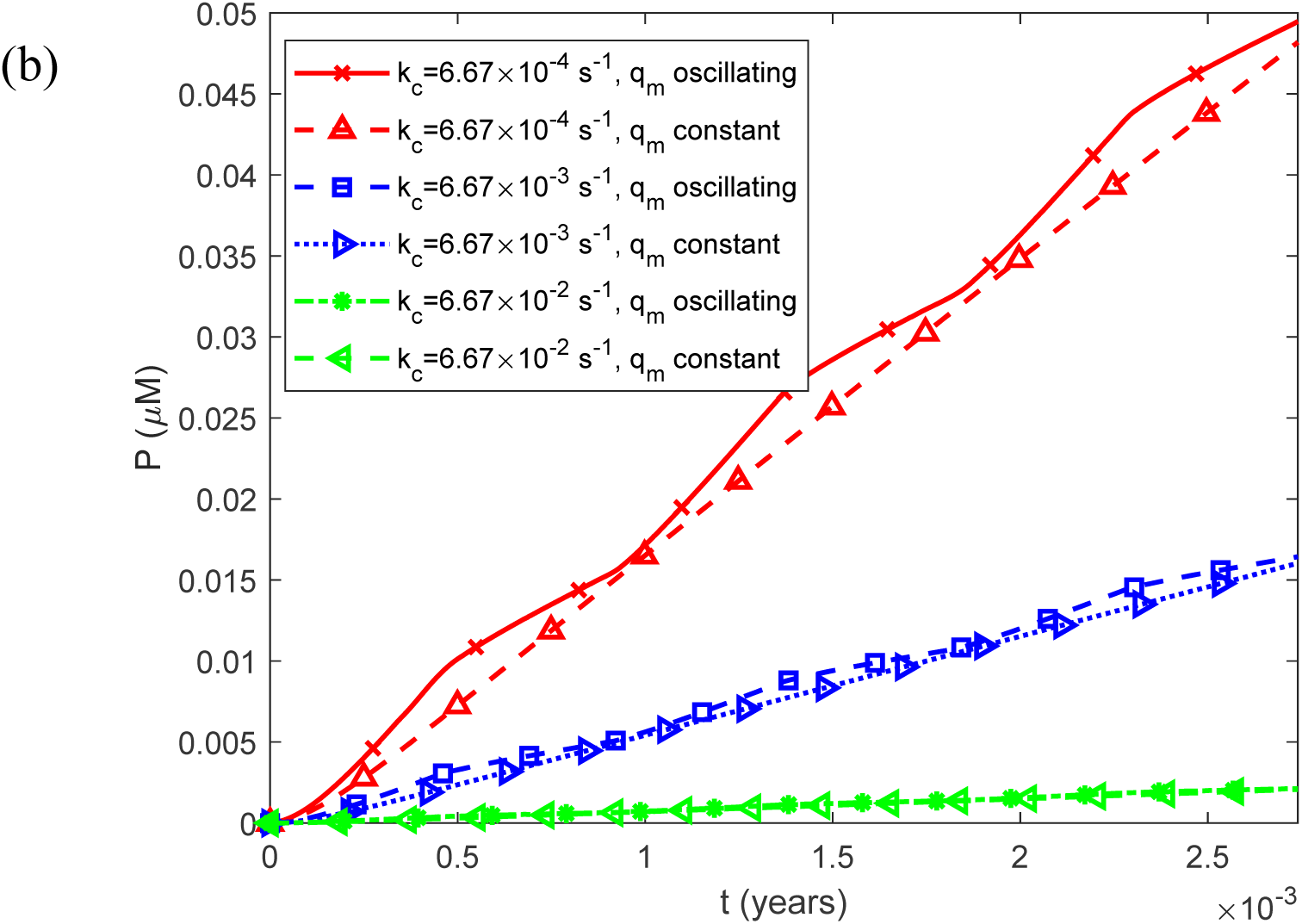
(a) Fibril particle concentration *P*(*t*) , i.e., the molar concentration of fibrils irrespective of length, as a function of time for a constant (time-averaged) monomer release rate of 1.25*q_m_*_0_ . (b) As in (a), but restricted to the time interval [0, 0.0027 years], equivalent to 1 day, illustrating the effect of oscillatory monomer release on the short-term dynamics of the fibril particle concentration. Results are shown for three representative values of the rate constant governing the clearance of IAPP monomers through fenestrated capillaries, *k_c_* .

**Fig. S7.**
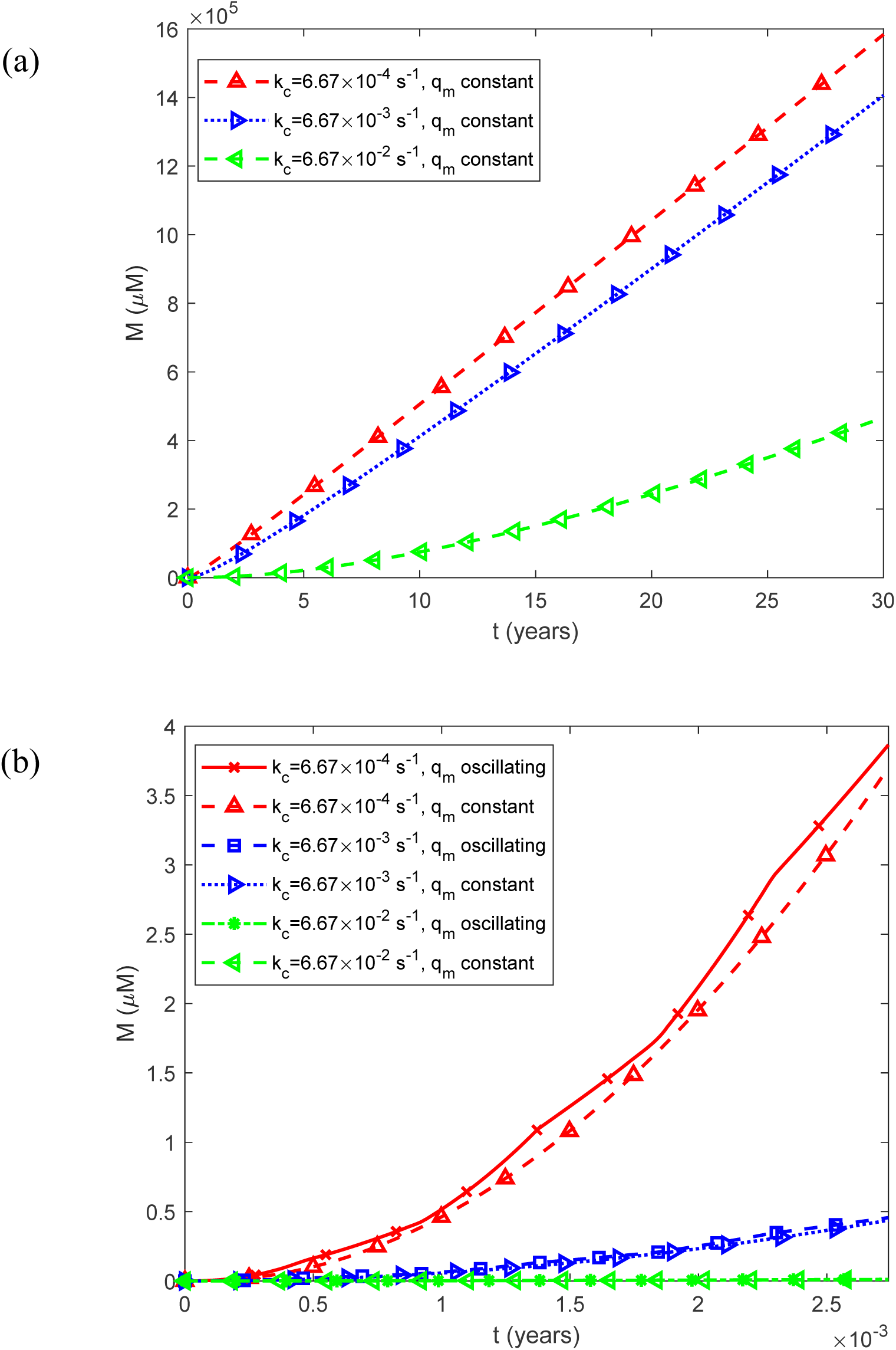
(a) Fibril mass concentration *M* (*t*) , expressed as the molar concentration of IAPP monomer equivalents incorporated into all fibrils, as a function of time for a constant (time-averaged) monomer release rate of 1.25*q_m_*_0_ . (b) As in (a), but restricted to the time interval [0, 0.0027 years], equivalent to 1 day, illustrating the effect of oscillatory monomer release on the short-term dynamics of total fibril mass. Results are shown for three representative values of the rate constant governing the clearance of IAPP monomers through fenestrated capillaries, *k_c_* .

**Fig. S8.**
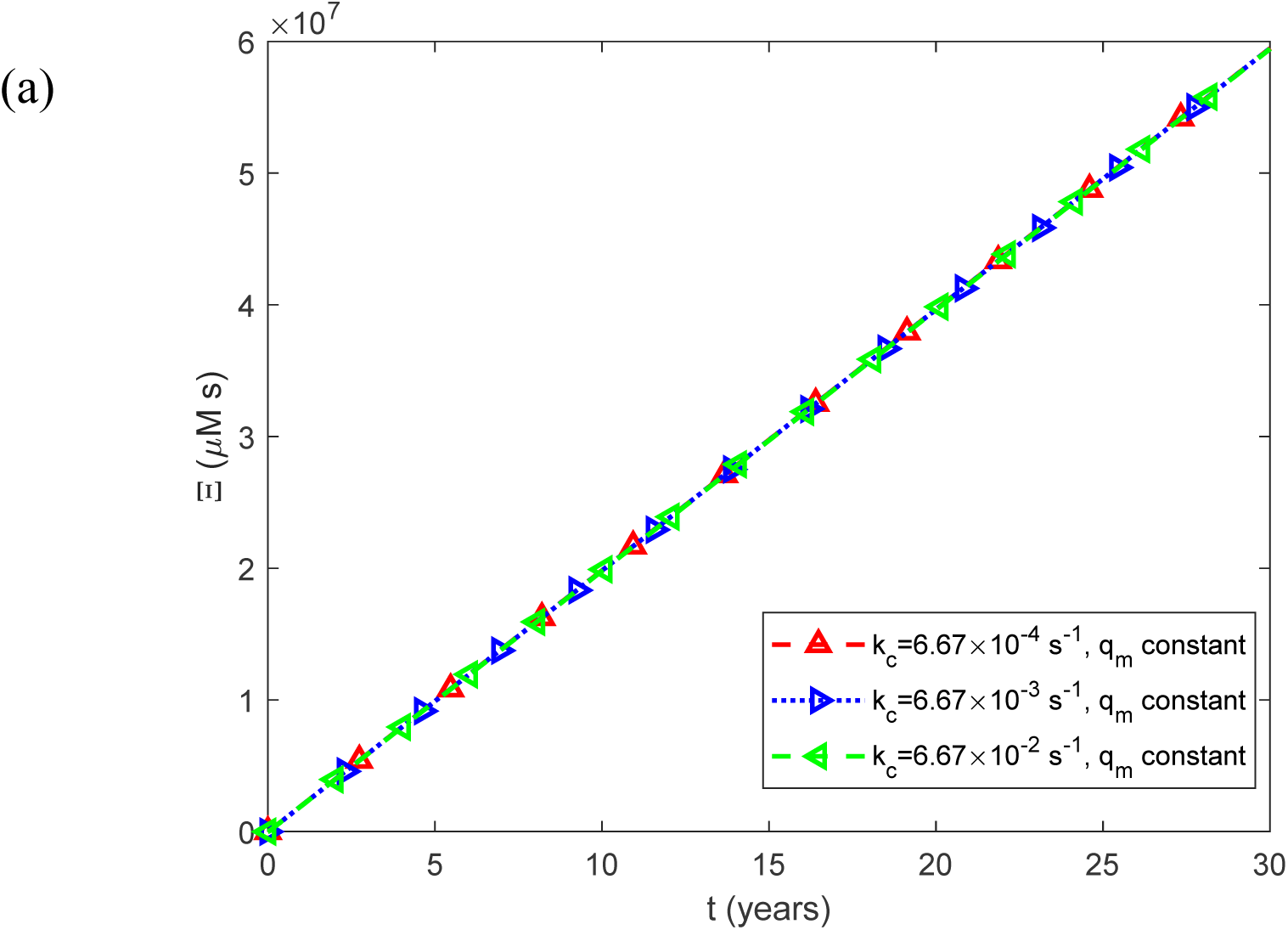
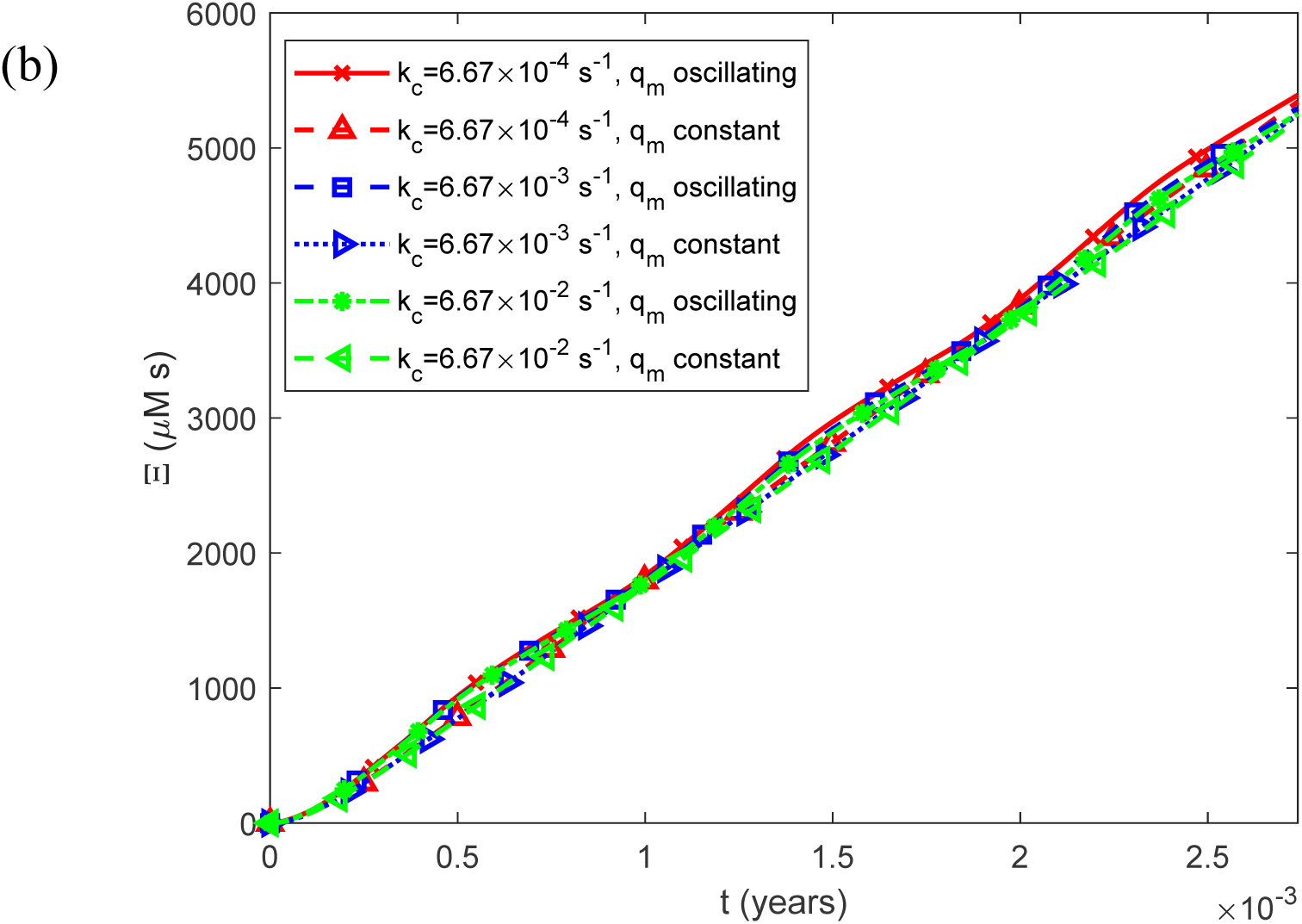
(a) Accumulated cytotoxicity (integrated IAPP oligomer exposure) as a function of time, Ξ(*t*) , for a constant (time-averaged) monomer release rate of 1.25*q_m_*_0_ . (b) As in (a), but restricted to the time interval [0, 0.0027 years], equivalent to 1 day, illustrating the effect of oscillatory monomer release on the short-term dynamics of accumulated cytotoxicity. Results are shown for three representative values of the rate constant governing the clearance of IAPP monomers through fenestrated capillaries, *k_c_* .

**Fig. S9.**
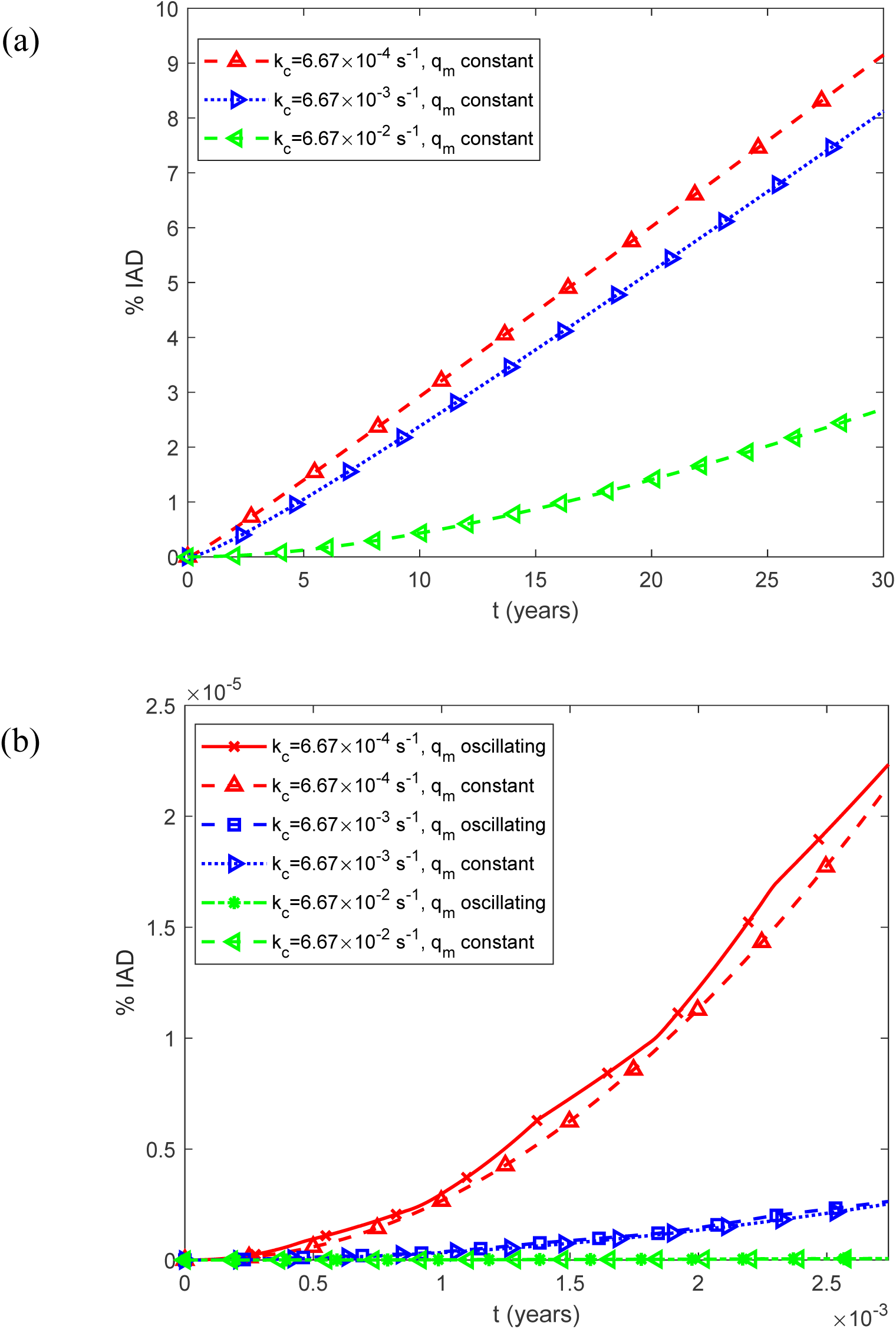
(a) Predicted percentage of the islet occupied by IADs, interpreted as the expected histological area fraction under the stereological assumption described in Section 2.2, as a function of time, %IADs, for a constant (time-averaged) monomer release rate of 1.25*q_m_*_0_ . (b) As in (a), but restricted to the time interval [0, 0.0027 years], equivalent to 1 day, illustrating the effect of oscillatory monomer release on the short-term dynamics of amyloid deposition. Results are shown for three representative values of the rate constant governing the clearance of IAPP monomers through fenestrated capillaries, *k_c_* .

**Fig. S10.**
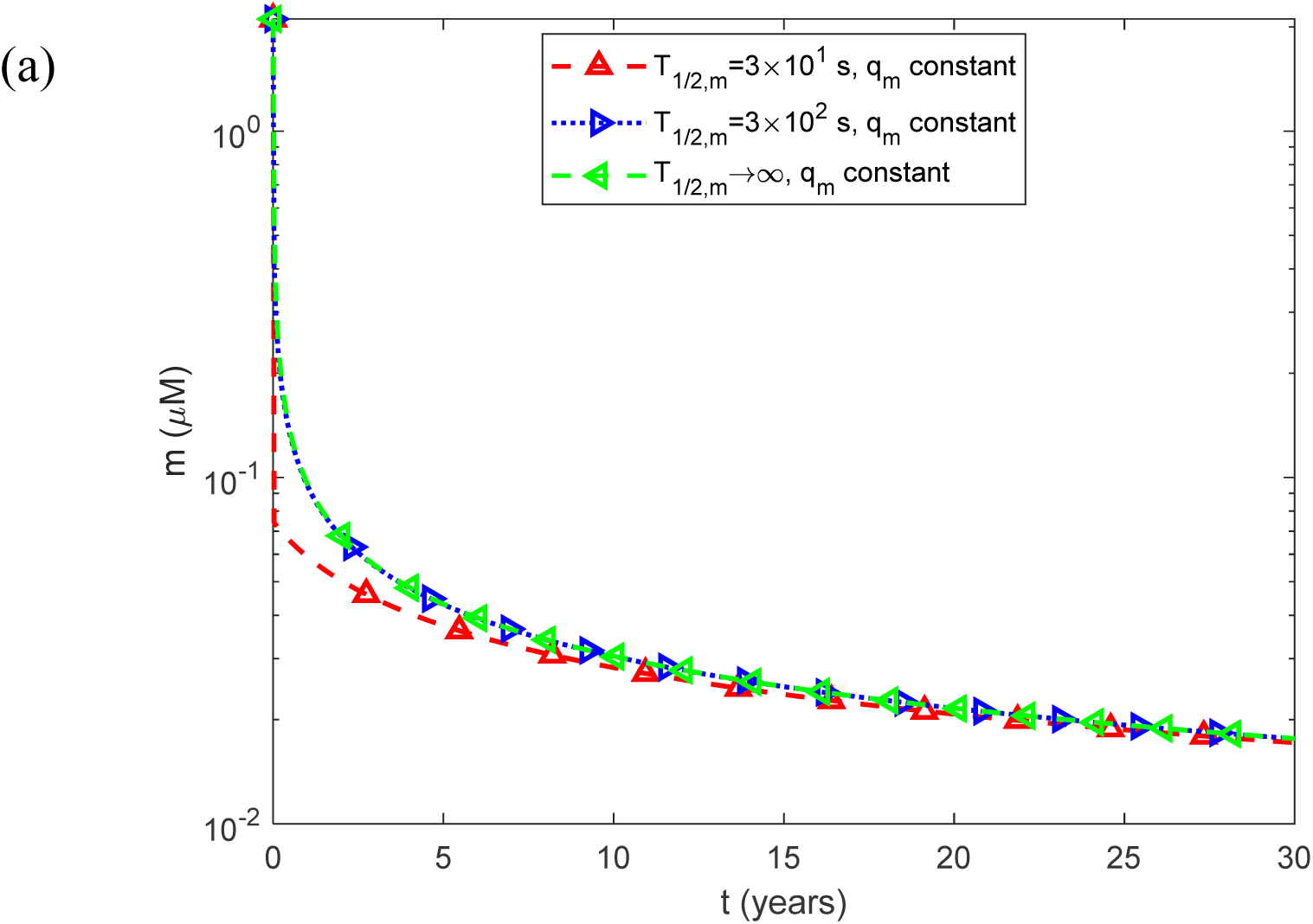
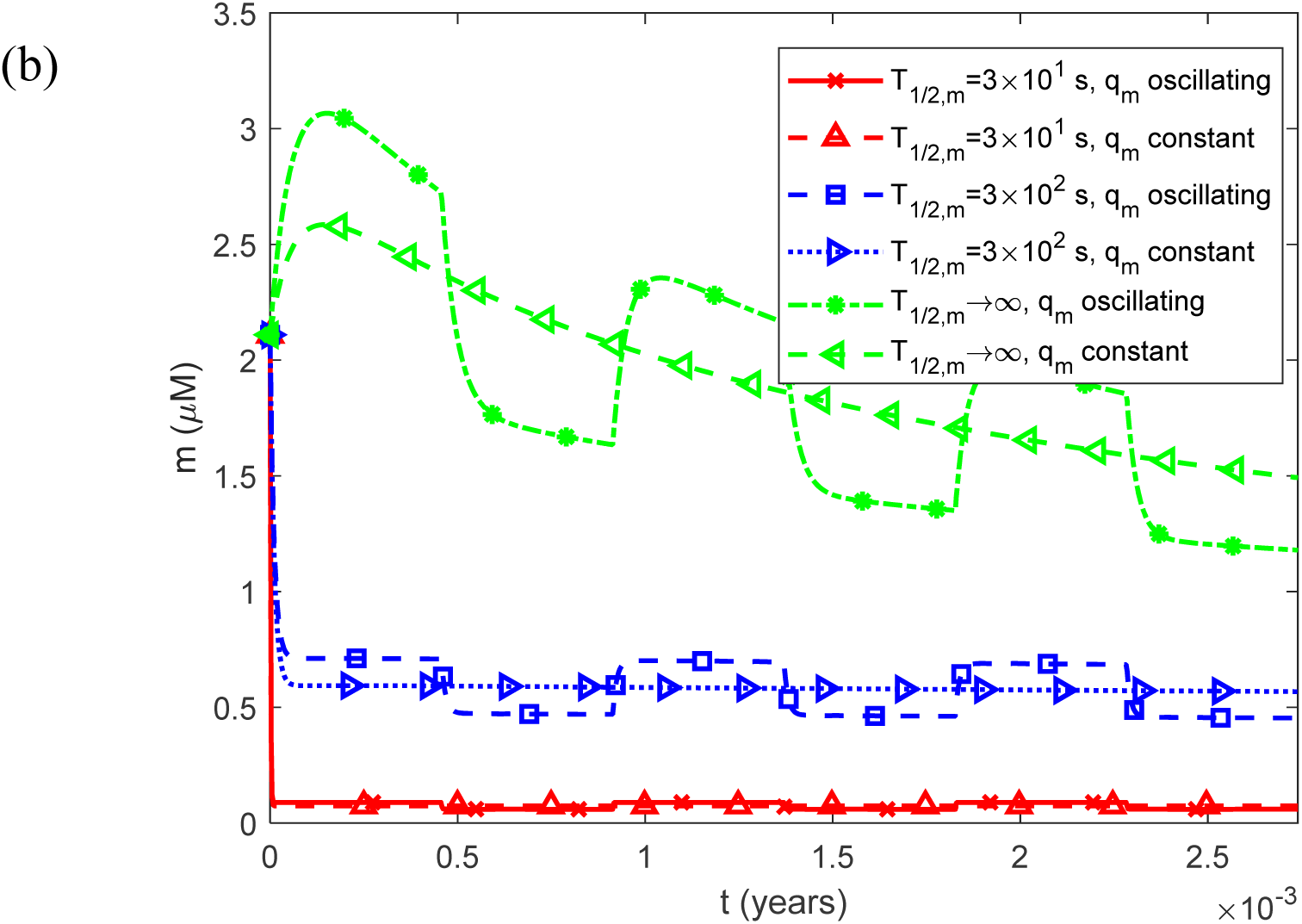
(a) Molar concentration of free IAPP monomers as a function of time, *m*(*t*) , for a constant (time-averaged) monomer release rate of 1.25*q_m_*_0_ . (b) As in (a), but restricted to the time interval [0, 0.0027 years], equivalent to 1 day, illustrating the effect of oscillatory monomer release on the short-term dynamics of the free monomer concentration. Results are shown for three representative values of the half-life characterizing the enzymatic degradation of IAPP monomers, *T*_1/2,*m*_ .

**Fig. S11.**
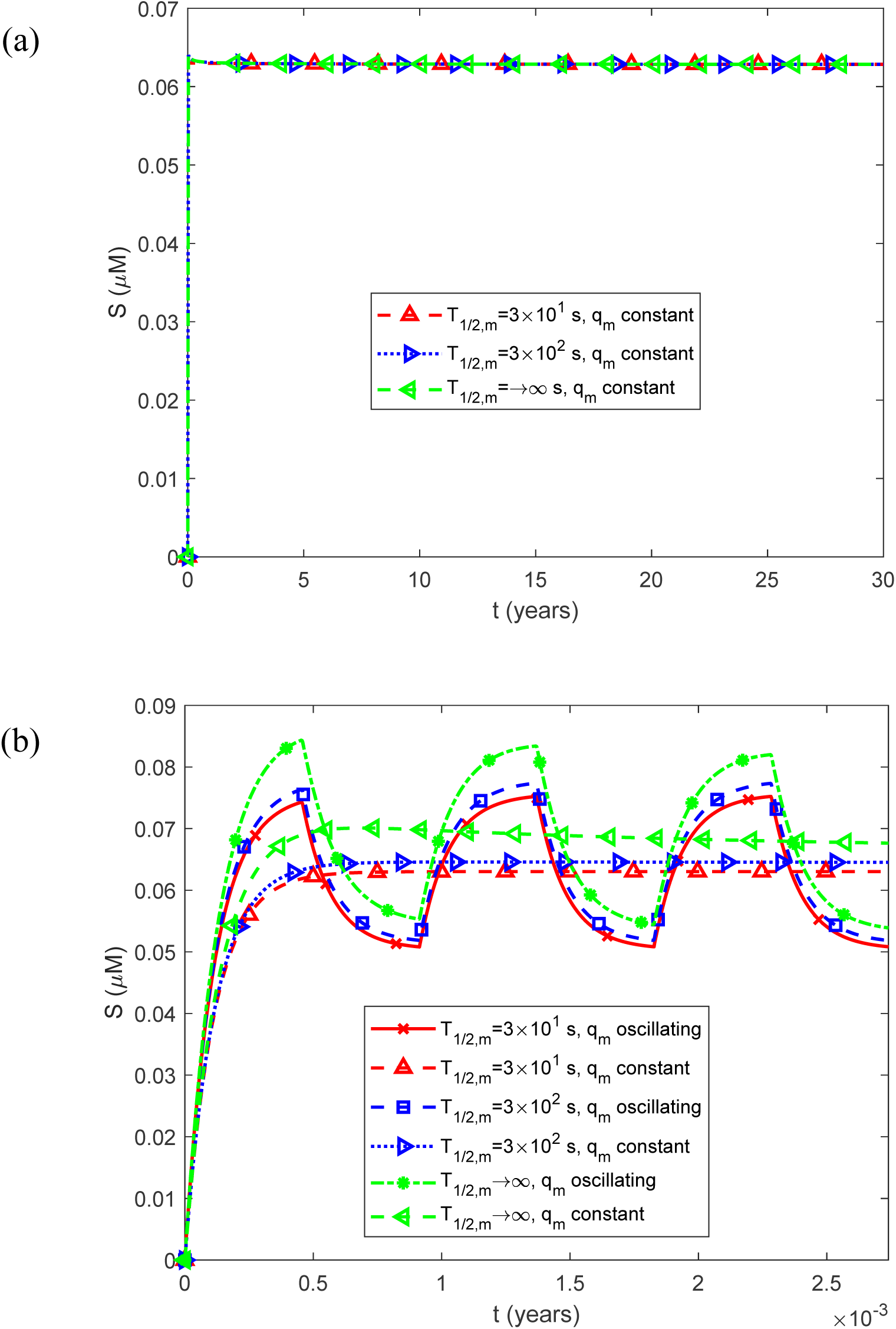
(a) Molar concentration of free IAPP oligomers as a function of time, *S*(*t*) , for a constant (time-averaged) monomer release rate of 1.25*q_m_*_0_ . (b) As in (a), but restricted to the time interval [0, 0.0027 years], equivalent to 1 day, illustrating the effect of oscillatory monomer release on the short-term dynamics of the free oligomer concentration. Results are shown for three representative values of the half-life characterizing the enzymatic degradation of IAPP monomers, *T*_1/2,*m*_ .

**Fig. S12.**
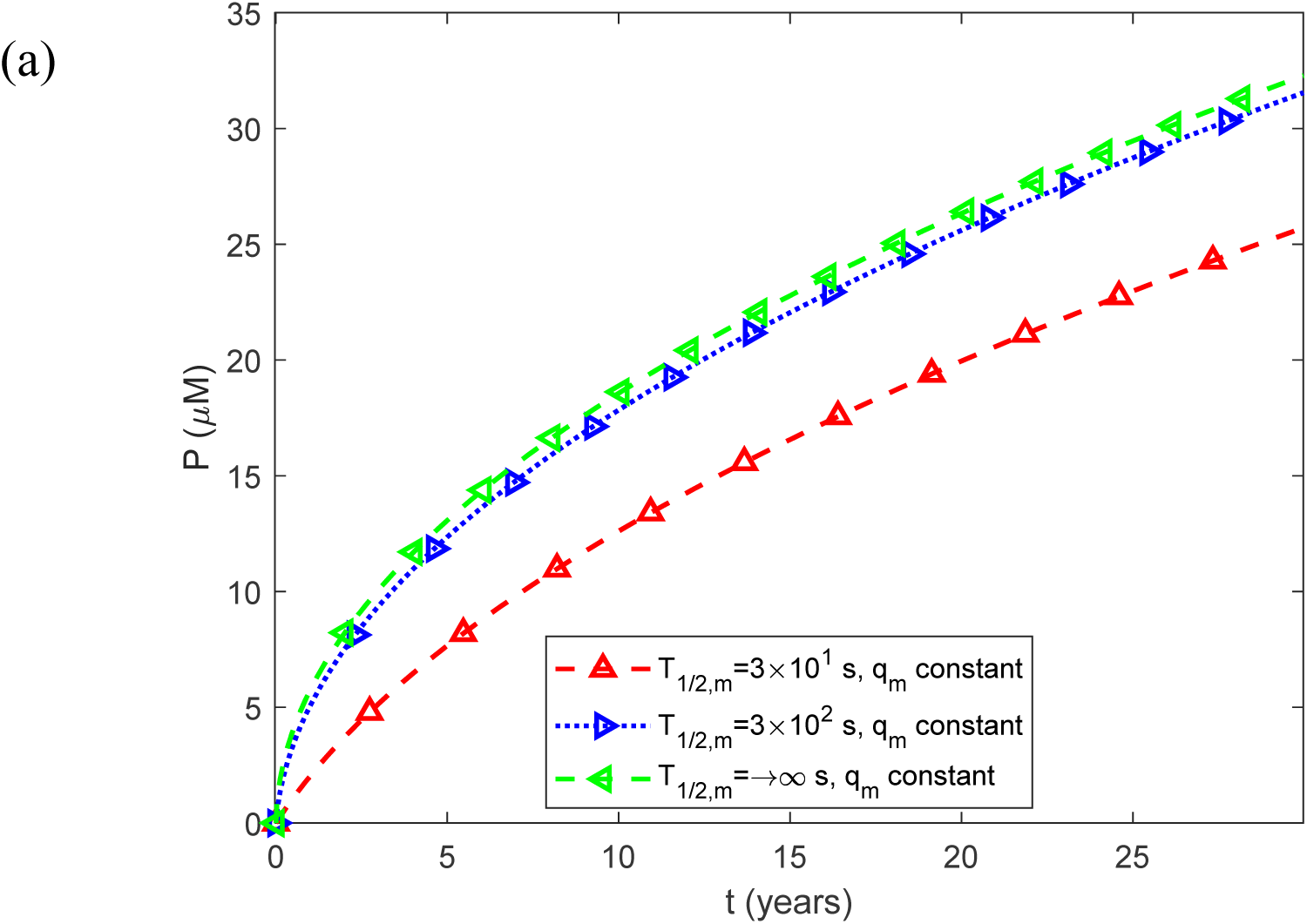
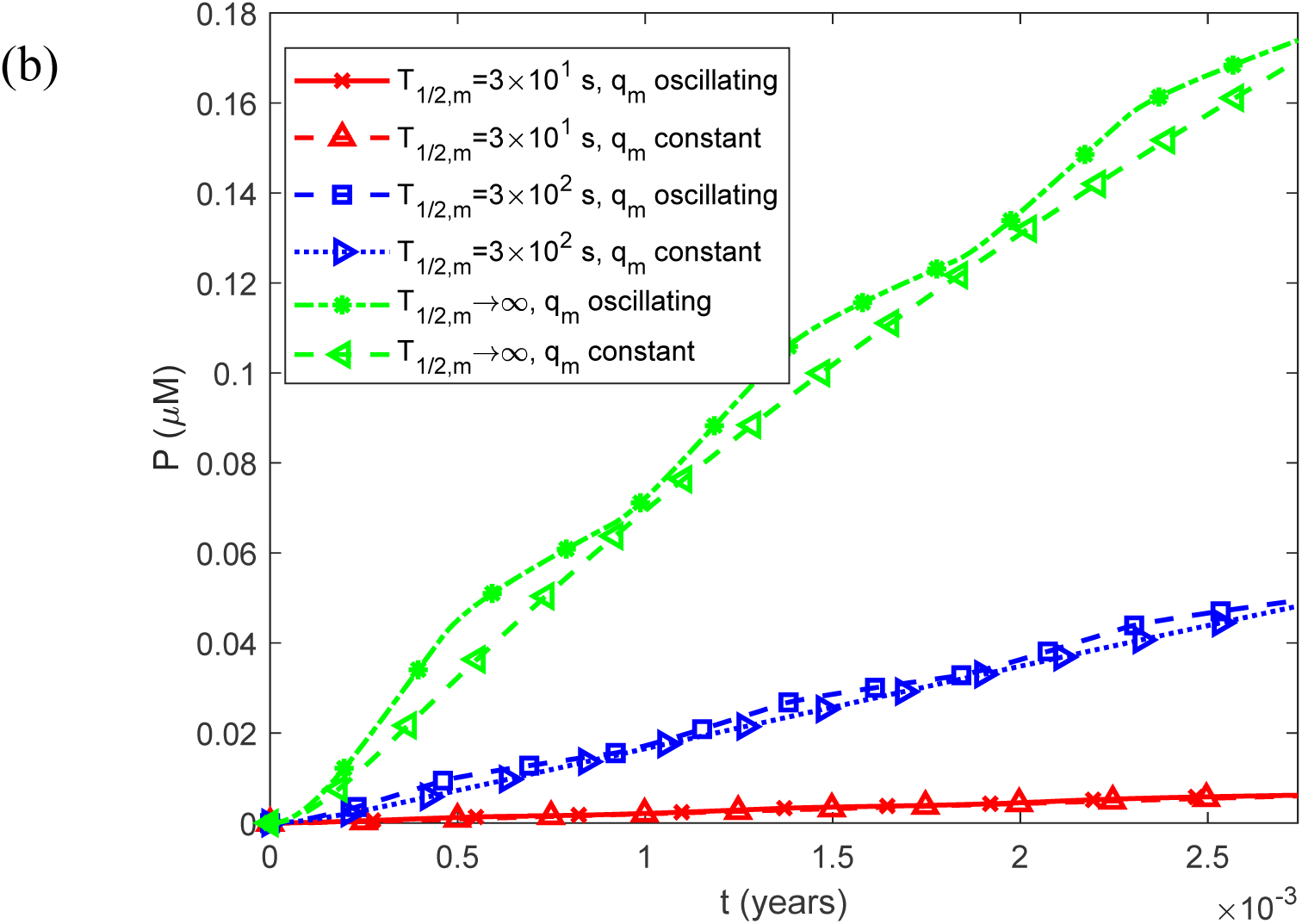
(a) Fibril particle concentration *P*(*t*) , i.e., the molar concentration of fibrils irrespective of length, as a function of time for a constant (time-averaged) monomer release rate of 1.25*q_m_*_0_ . (b) As in (a), but restricted to the time interval [0, 0.0027 years], equivalent to 1 day, illustrating the effect of oscillatory monomer release on the short-term dynamics of the fibril particle concentration. Results are shown for three representative values of the half-life characterizing the enzymatic degradation of IAPP monomers, *T*_1/2,*m*_ .

**Fig. S13.**
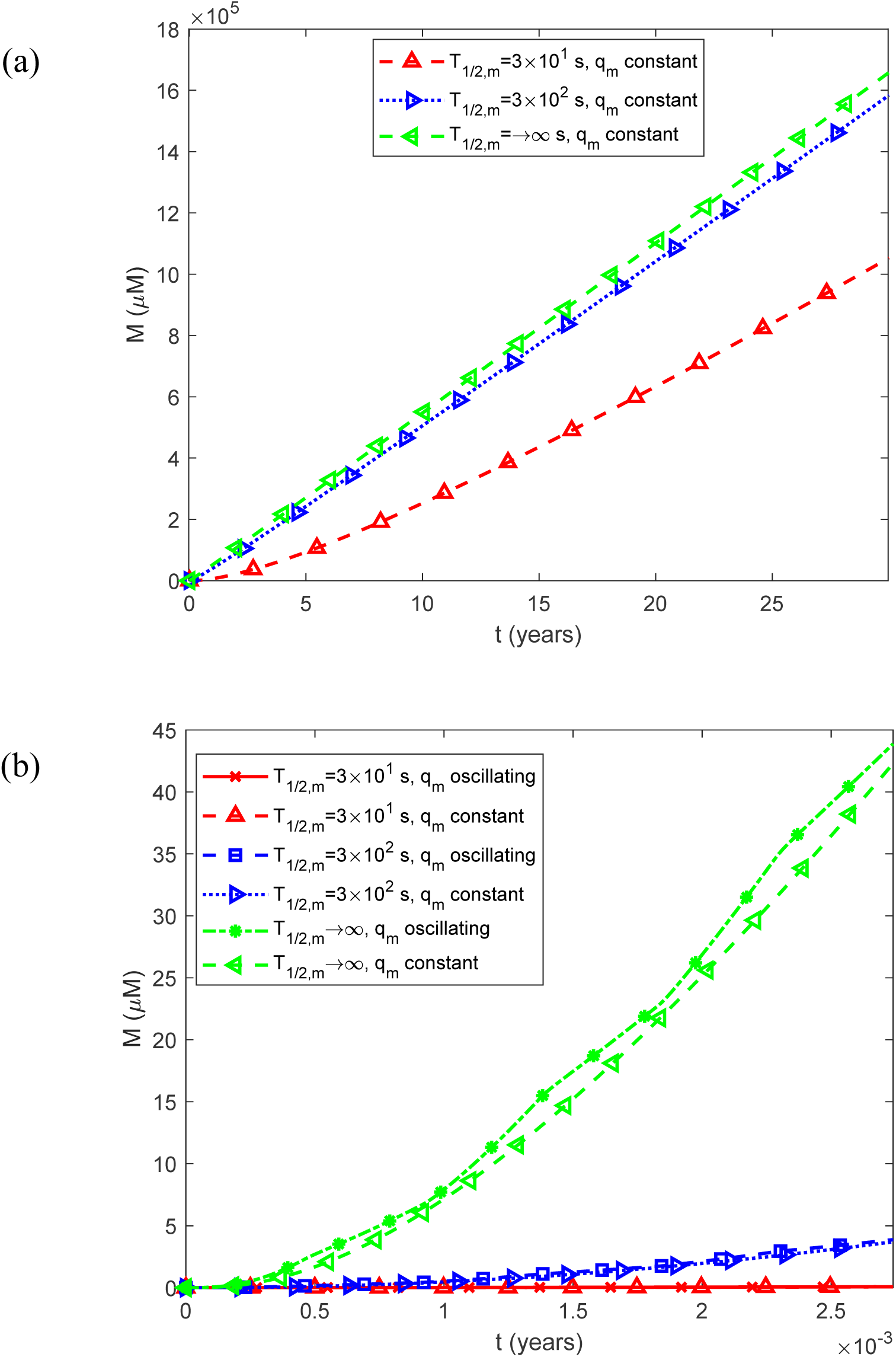
(a) Fibril mass concentration *M* (*t*) , expressed as the molar concentration of IAPP monomer equivalents incorporated into all fibrils, as a function of time for a constant (time-averaged) monomer release rate of 1.25*q_m_*_0_ . (b) As in (a), but restricted to the time interval [0, 0.0027 years], equivalent to 1 day, illustrating the effect of oscillatory monomer release on the short-term dynamics of total fibril mass. Results are shown for three representative values of the half-life characterizing the enzymatic degradation of IAPP monomers, *T*_1/2,*m*_ .

**Fig. S14.**
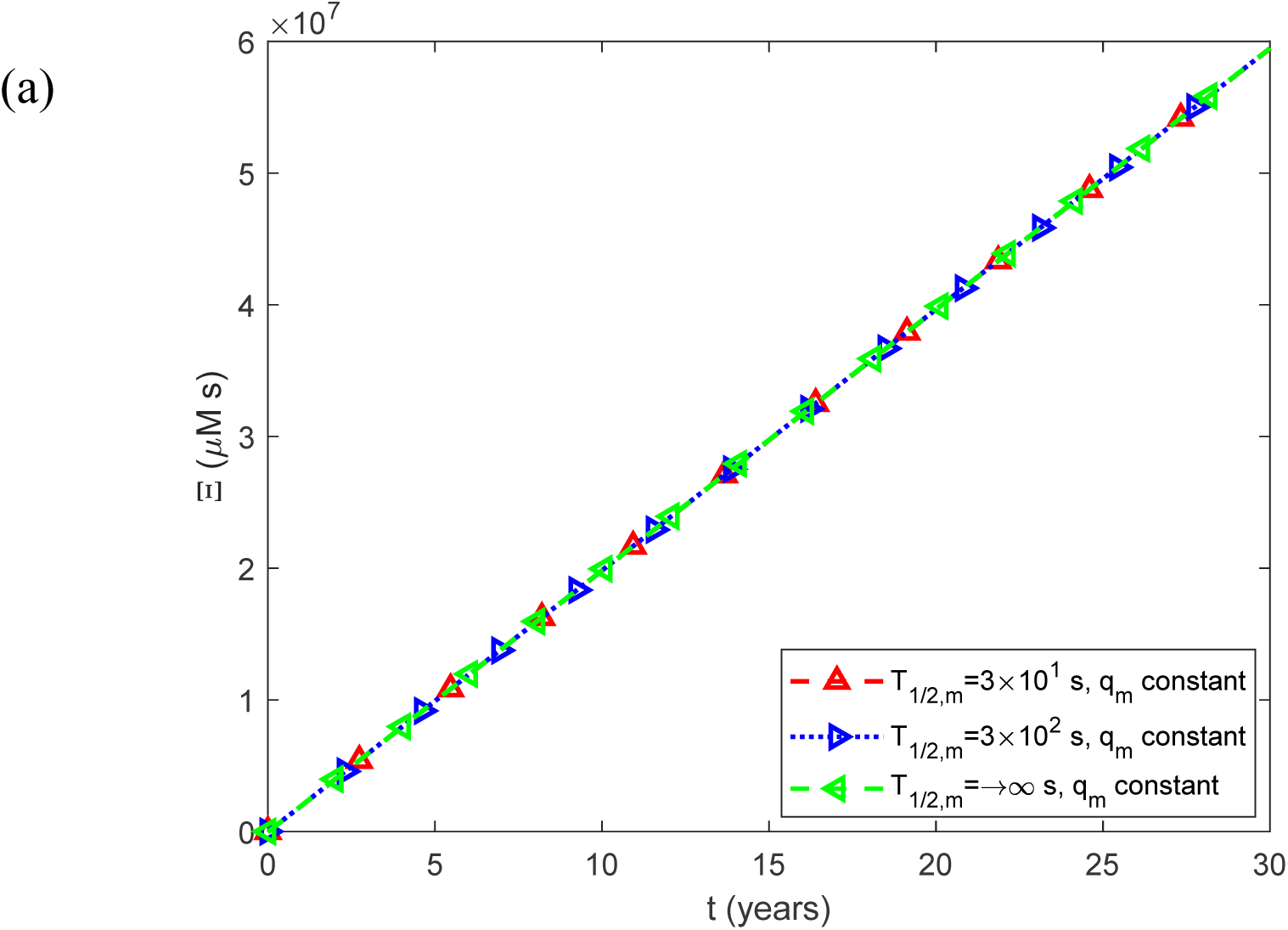
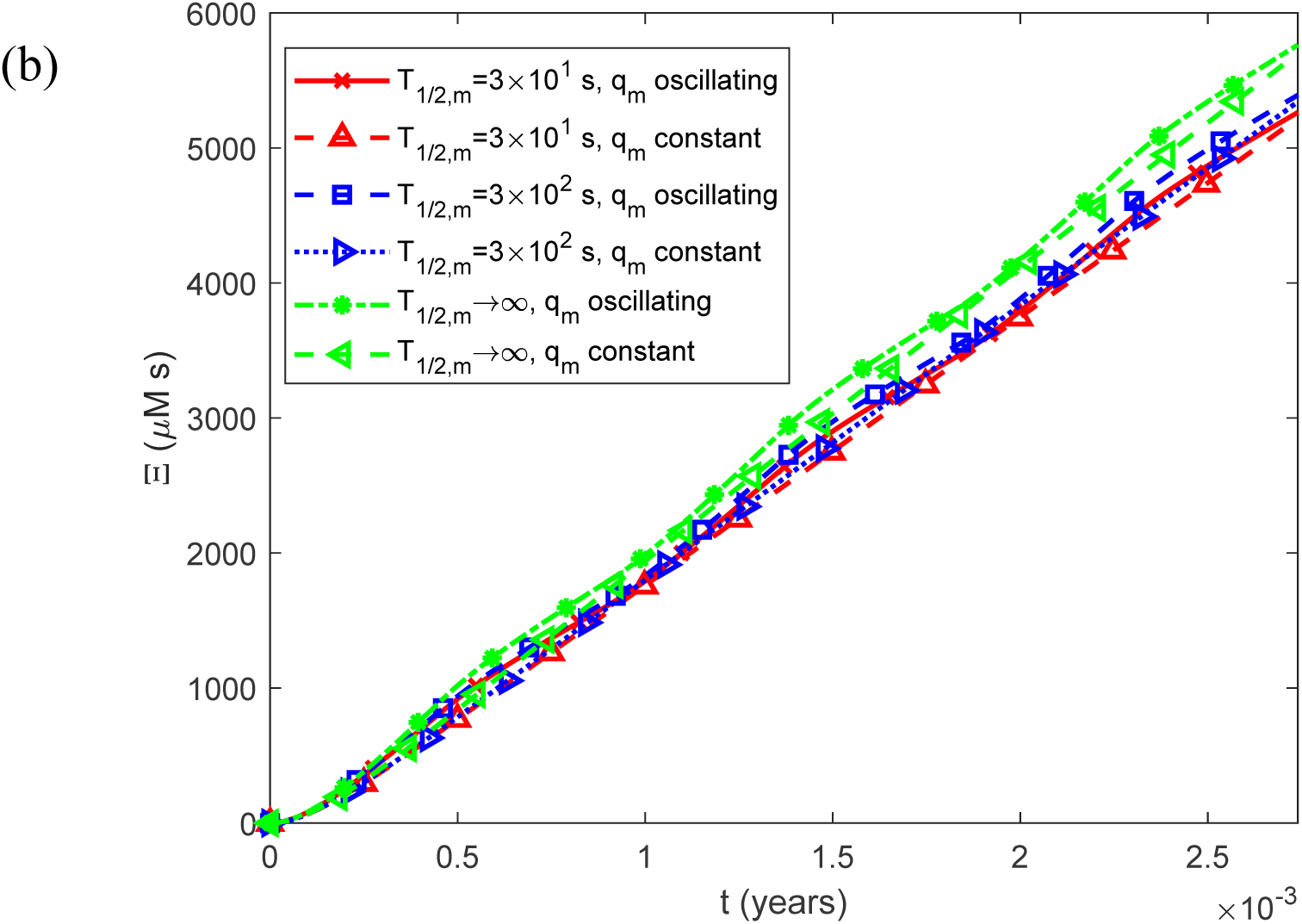
(a) Accumulated cytotoxicity (integrated IAPP oligomer exposure) as a function of time, Ξ(*t*) , for a constant (time-averaged) monomer release rate of 1.25*q_m_*_0_ . (b) As in (a), but restricted to the time interval [0, 0.0027 years], equivalent to 1 day, illustrating the effect of oscillatory monomer release on the short-term dynamics of accumulated cytotoxicity. Results are shown for three representative values of the half-life characterizing the enzymatic degradation of IAPP monomers, *T*_1/2,*m*_ .

**Fig. S15.**
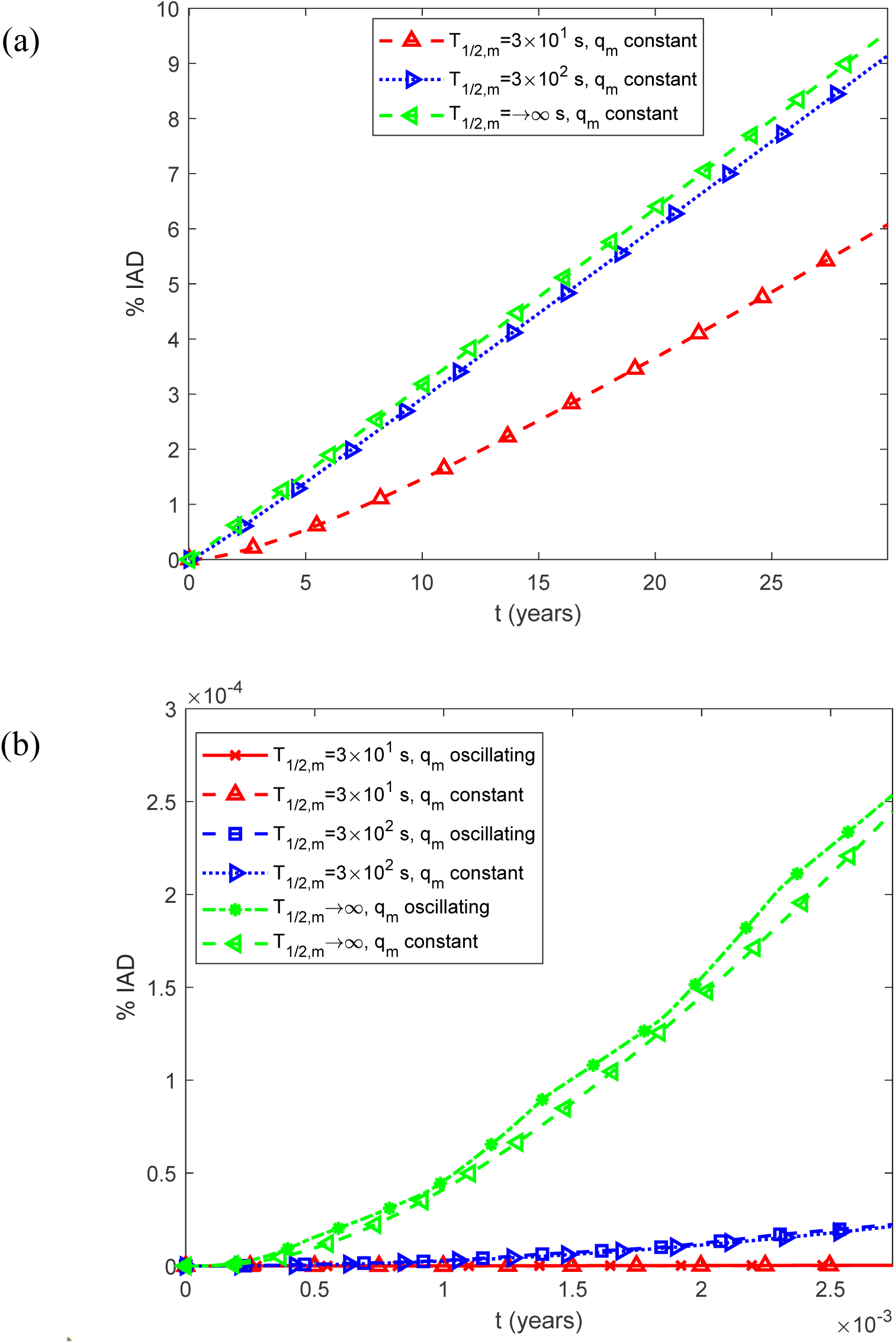
(a) Predicted percentage of the islet occupied by IADs, interpreted as the expected histological area fraction under the stereological assumption described in Section 2.2, as a function of time, %IADs, for a constant (time-averaged) monomer release rate of 1.25*q_m_*_0_ . (b) As in (a), but restricted to the time interval [0, 0.0027 years], equivalent to 1 day, illustrating the effect of oscillatory monomer release on the short-term dynamics of amyloid deposition. Results are shown for three representative values of the half-life characterizing the enzymatic degradation of IAPP monomers, *T*_1/2,*m*_ .

**Fig. S16.**
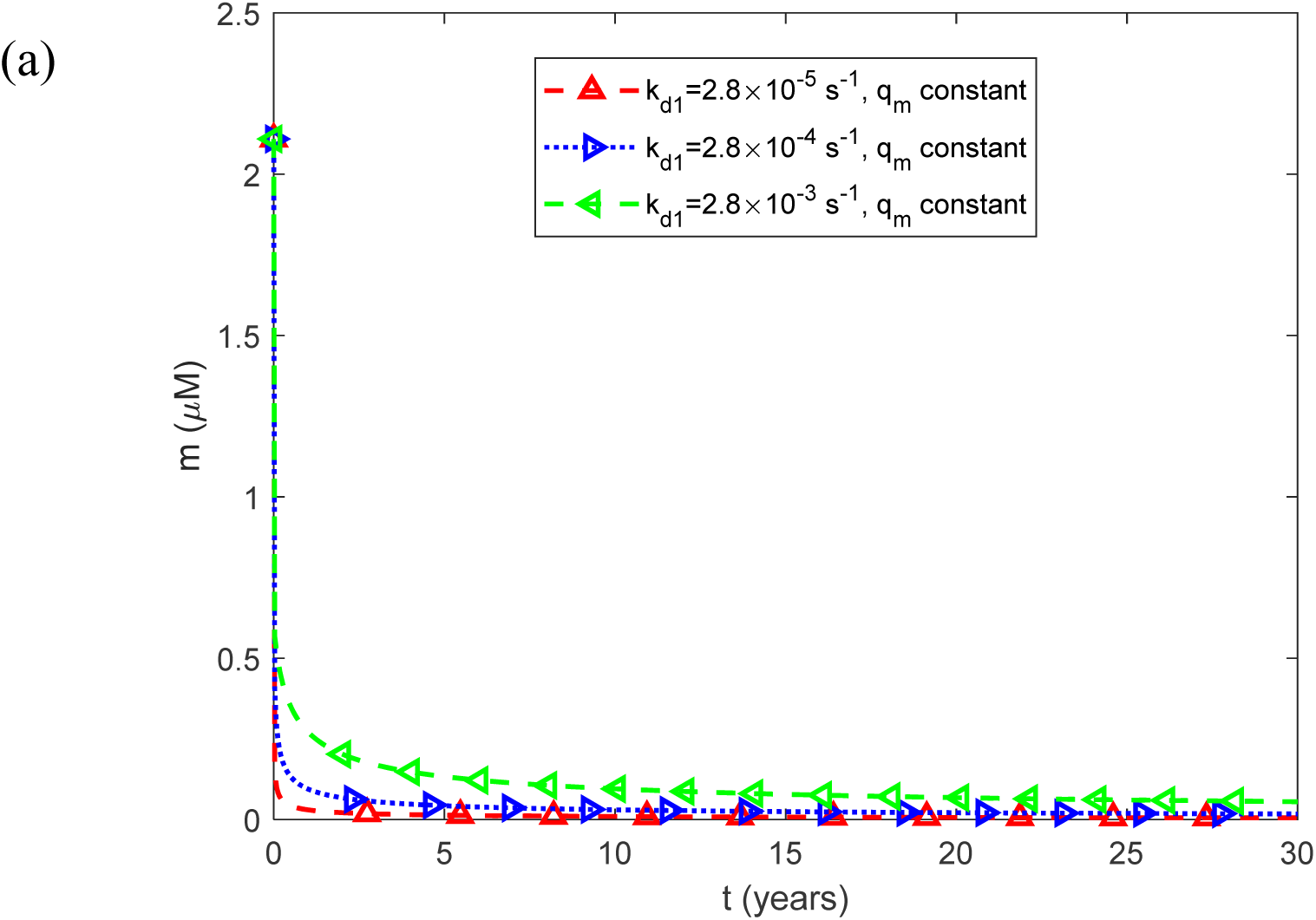
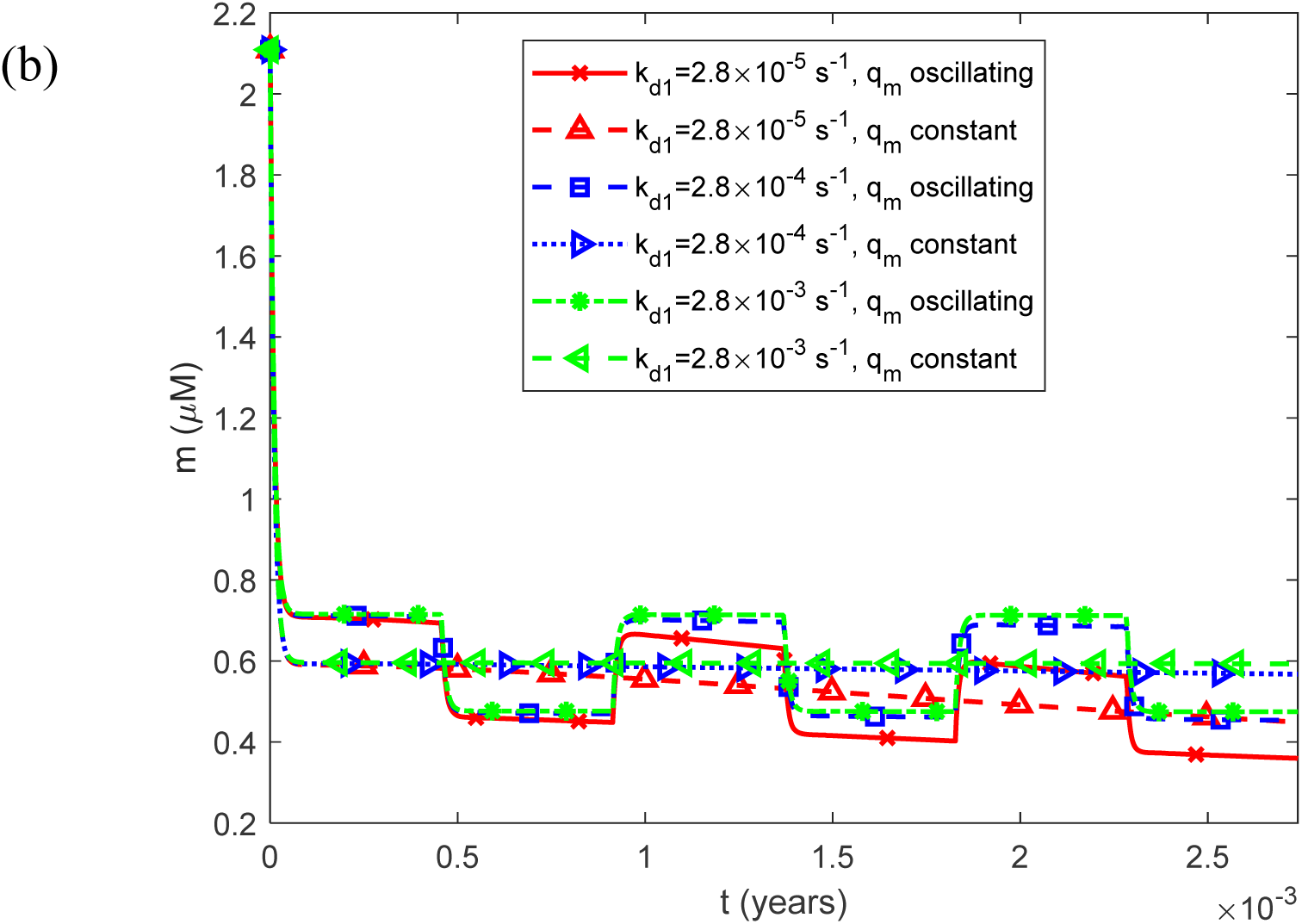
(a) Molar concentration of free IAPP monomers as a function of time, *m*(*t*) , for a constant (time-averaged) monomer release rate of 1.25*q_m_*_0_ . (b) As in (a), but restricted to the time interval [0, 0.0027 years], equivalent to 1 day, illustrating the effect of oscillatory monomer release on the short-term dynamics of the free monomer concentration. Results are shown for three representative values of the rate constant for the spontaneous dissociation of IAPP oligomers into constituent monomers, *k_d_*_1_.

**Fig. S17.**
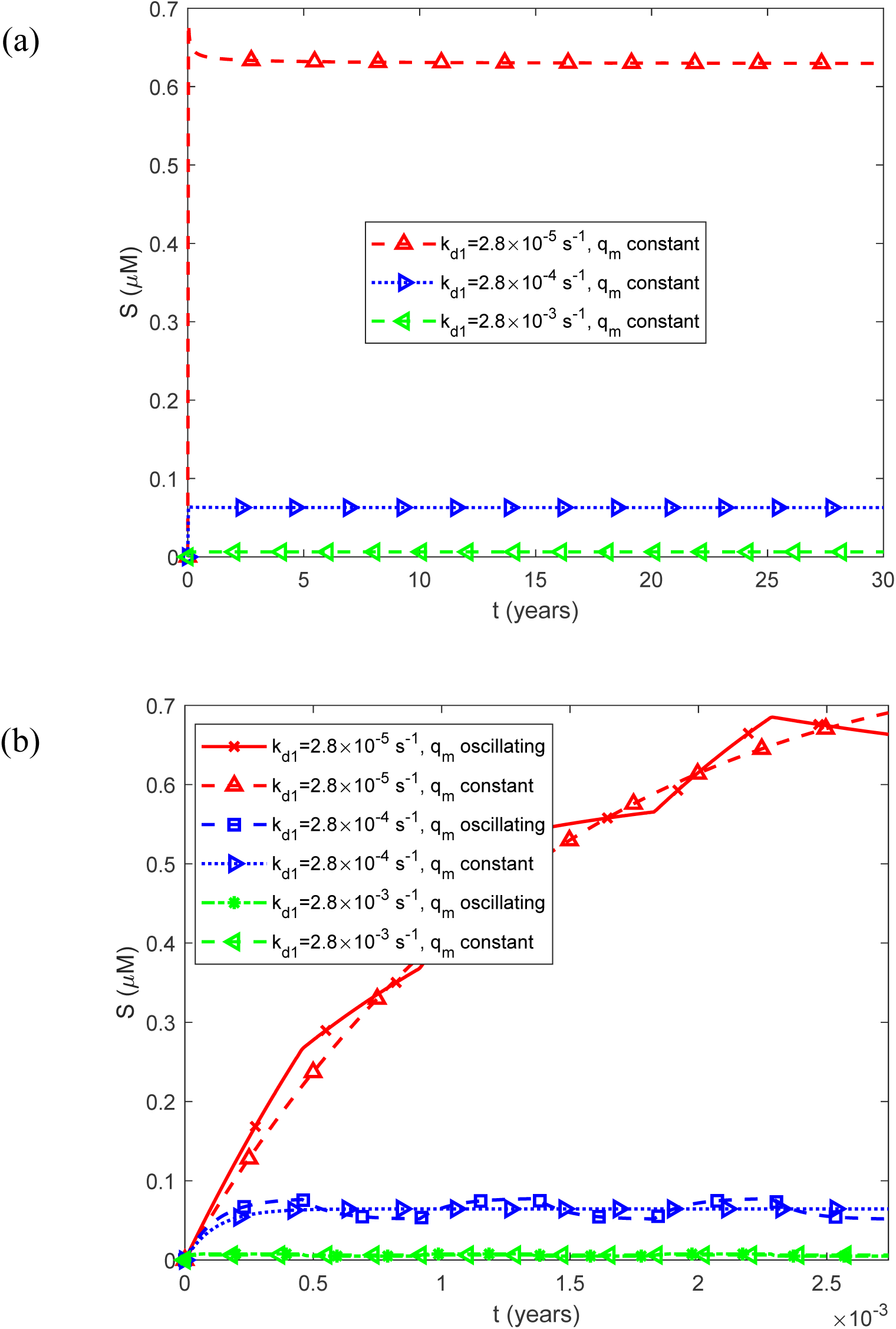
(a) Molar concentration of free IAPP oligomers as a function of time, *S*(*t*) , for a constant (time-averaged) monomer release rate of 1.25*q_m_*_0_ . (b) As in (a), but restricted to the time interval [0, 0.0027 years], equivalent to 1 day, illustrating the effect of oscillatory monomer release on the short-term dynamics of the free oligomer concentration. Results are shown for three representative values of the rate constant for the spontaneous dissociation of IAPP oligomers into constituent monomers, *k_d_*_1_.

**Fig. S18.**
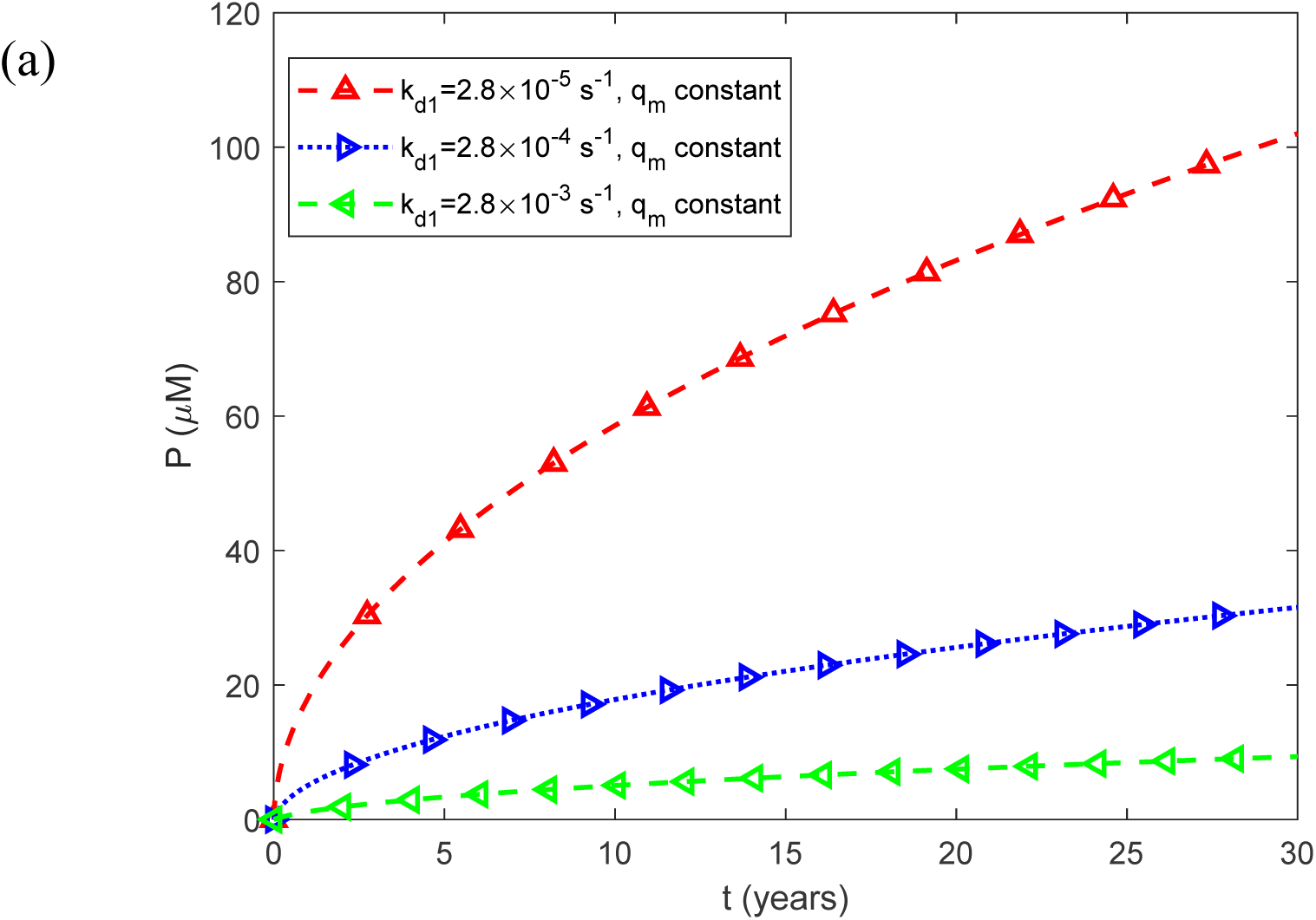
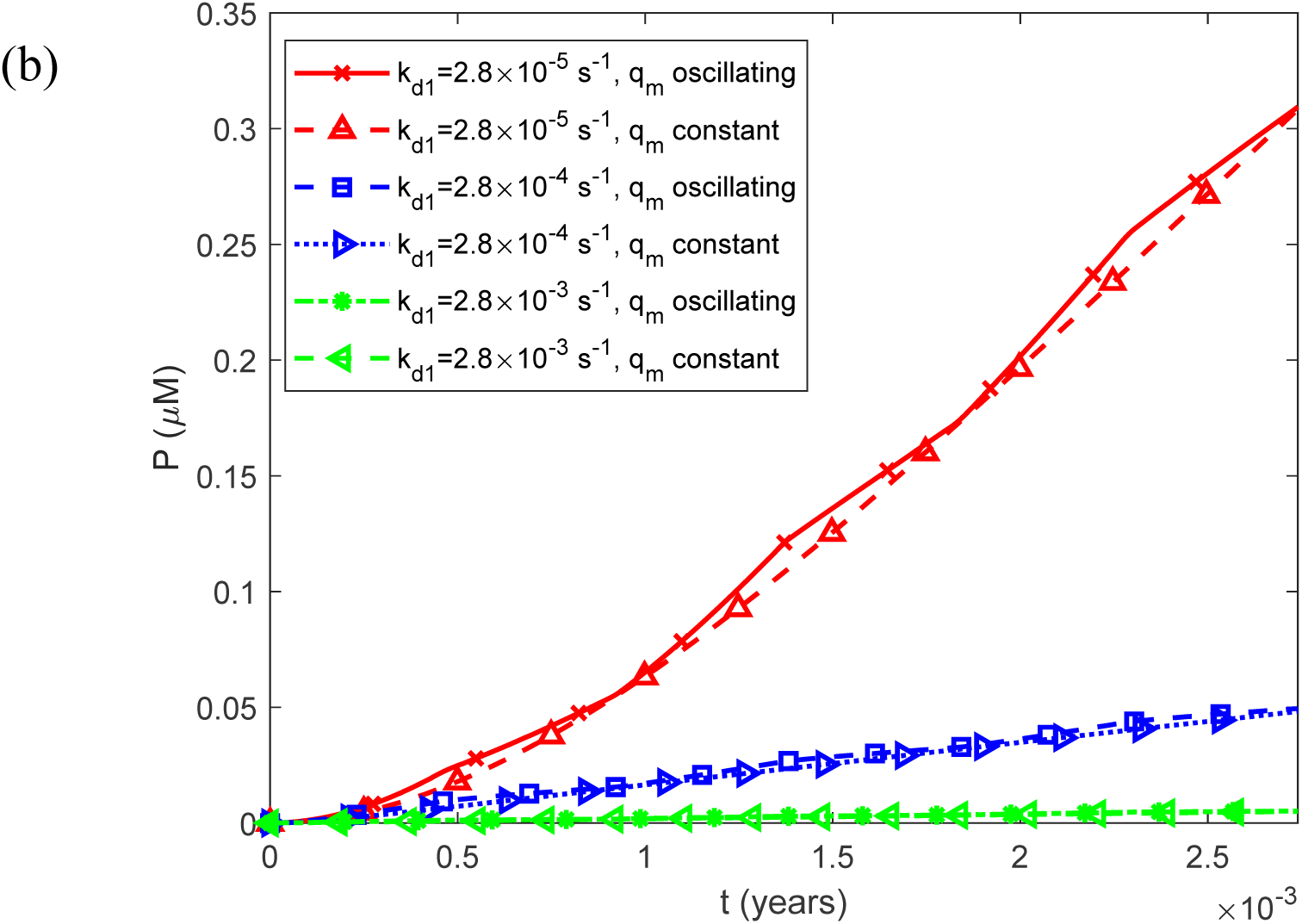
(a) Fibril particle concentration *P*(*t*) , i.e., the molar concentration of fibrils irrespective of length, as a function of time for a constant (time-averaged) monomer release rate of 1.25*q_m_*_0_ . (b) As in (a), but restricted to the time interval [0, 0.0027 years], equivalent to 1 day, illustrating the effect of oscillatory monomer release on the short-term dynamics of the fibril particle concentration. Results are shown for three representative values of the rate constant for the spontaneous dissociation of IAPP oligomers into constituent monomers, *k_d_*_1_.

**Fig. S19.**
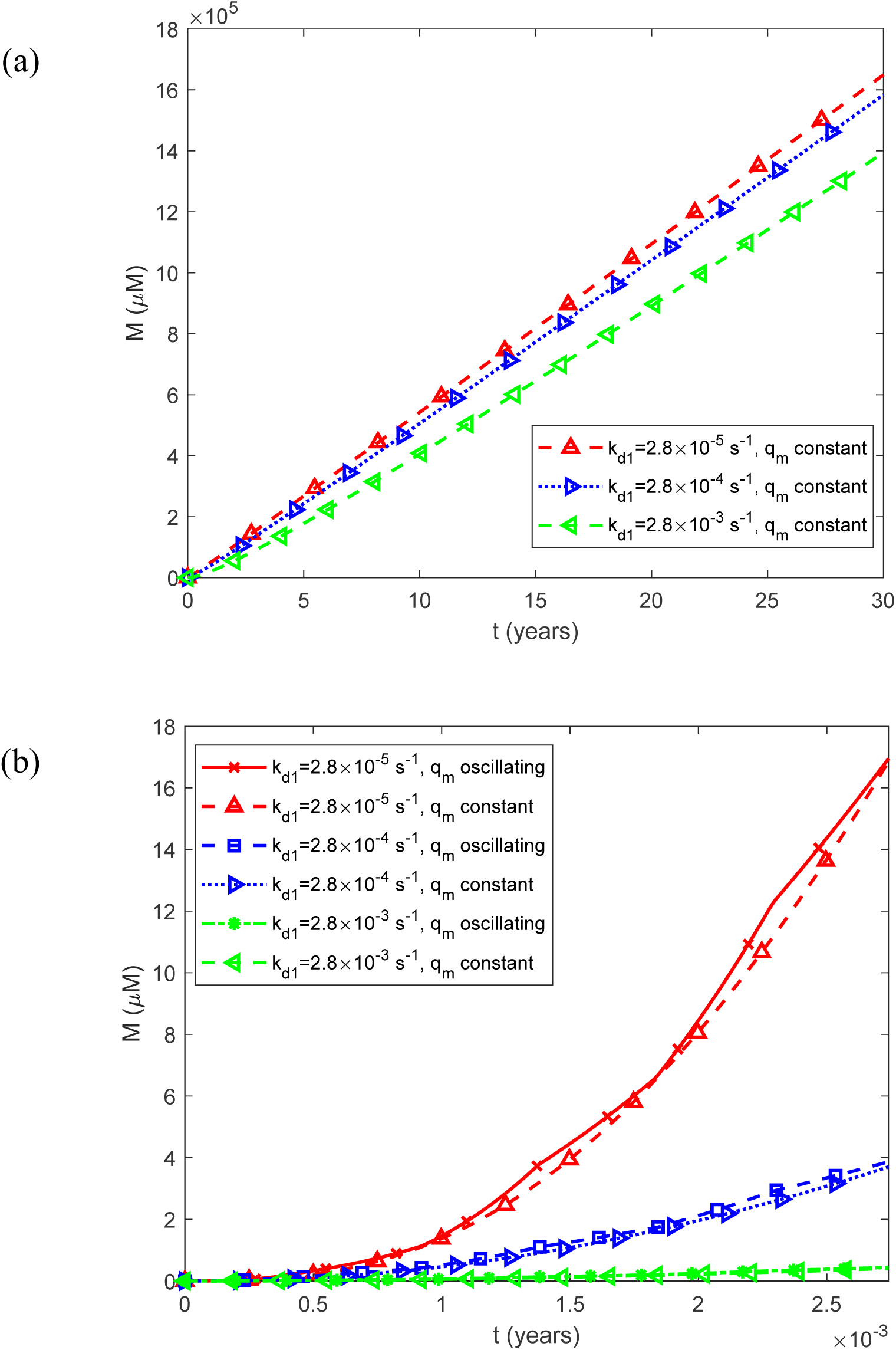
(a) Fibril mass concentration *M* (*t*) , expressed as the molar concentration of IAPP monomer equivalents incorporated into all fibrils, as a function of time for a constant (time-averaged) monomer release rate of 1.25*q_m_*_0_ . (b) As in (a), but restricted to the time interval [0, 0.0027 years], equivalent to 1 day, illustrating the effect of oscillatory monomer release on the short-term dynamics of total fibril mass. Results are shown for three representative values of the rate constant for the spontaneous dissociation of IAPP oligomers into constituent monomers, *k_d_*_1_.

**Fig. S20.**
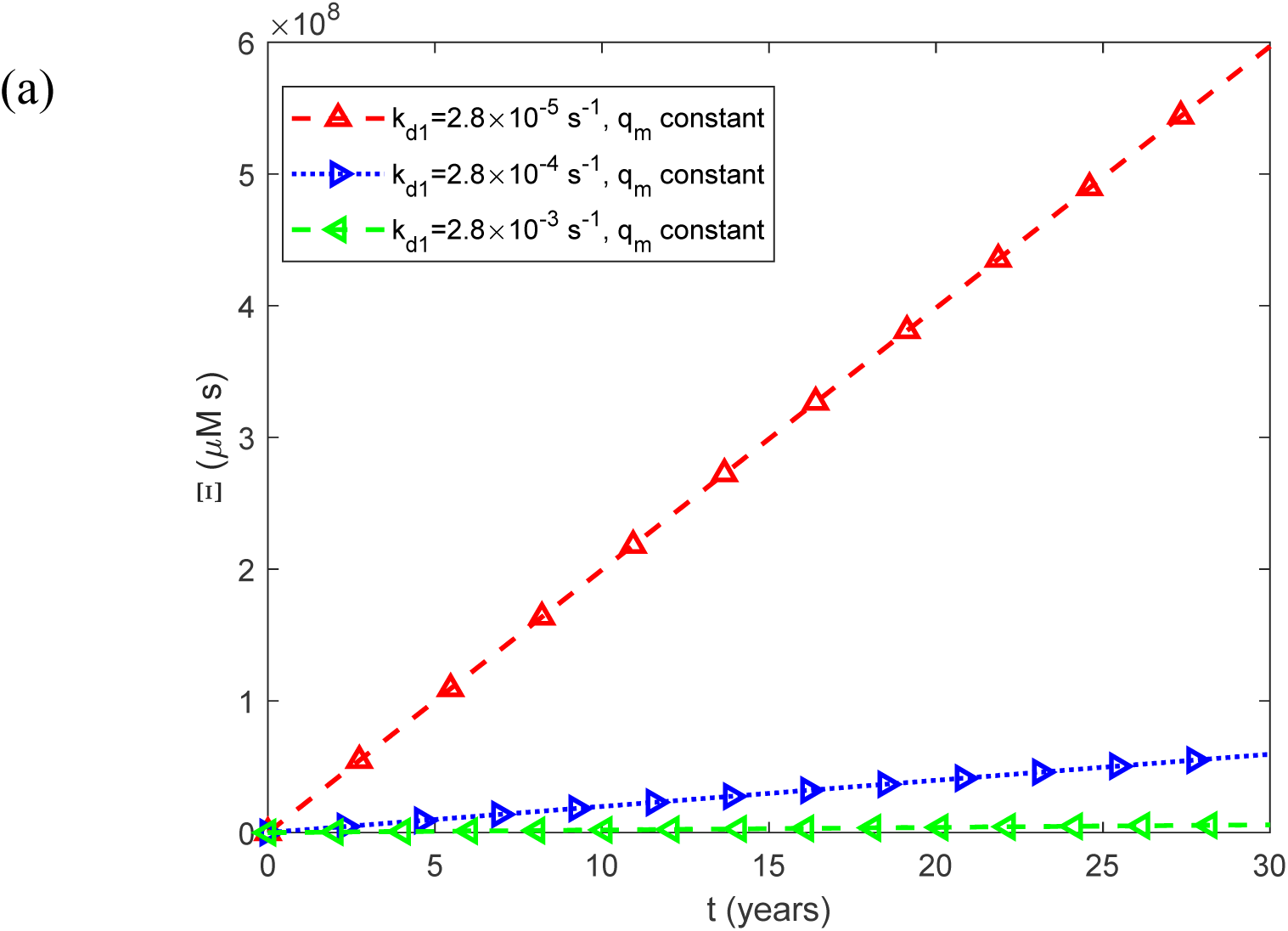
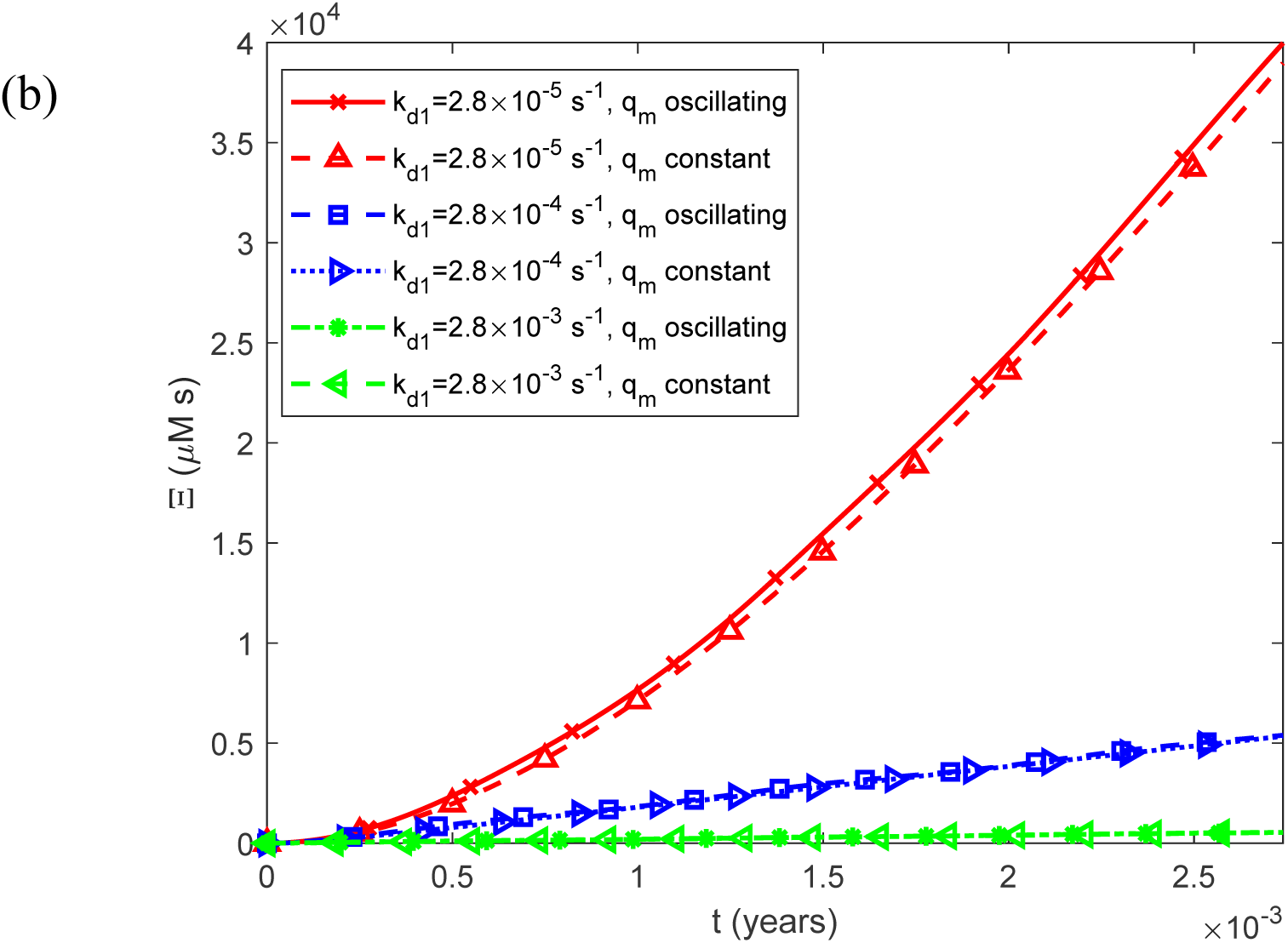
(a) Accumulated cytotoxicity (integrated IAPP oligomer exposure) as a function of time, Ξ(*t*) , for a constant (time-averaged) monomer release rate of 1.25*q_m_*_0_ . (b) As in (a), but restricted to the time interval [0, 0.0027 years], equivalent to 1 day, illustrating the effect of oscillatory monomer release on the short-term dynamics of accumulated cytotoxicity. Results are shown for three representative values of the rate constant for the spontaneous dissociation of IAPP oligomers into constituent monomers, *k_d_*_1_.

**Fig. S21.**
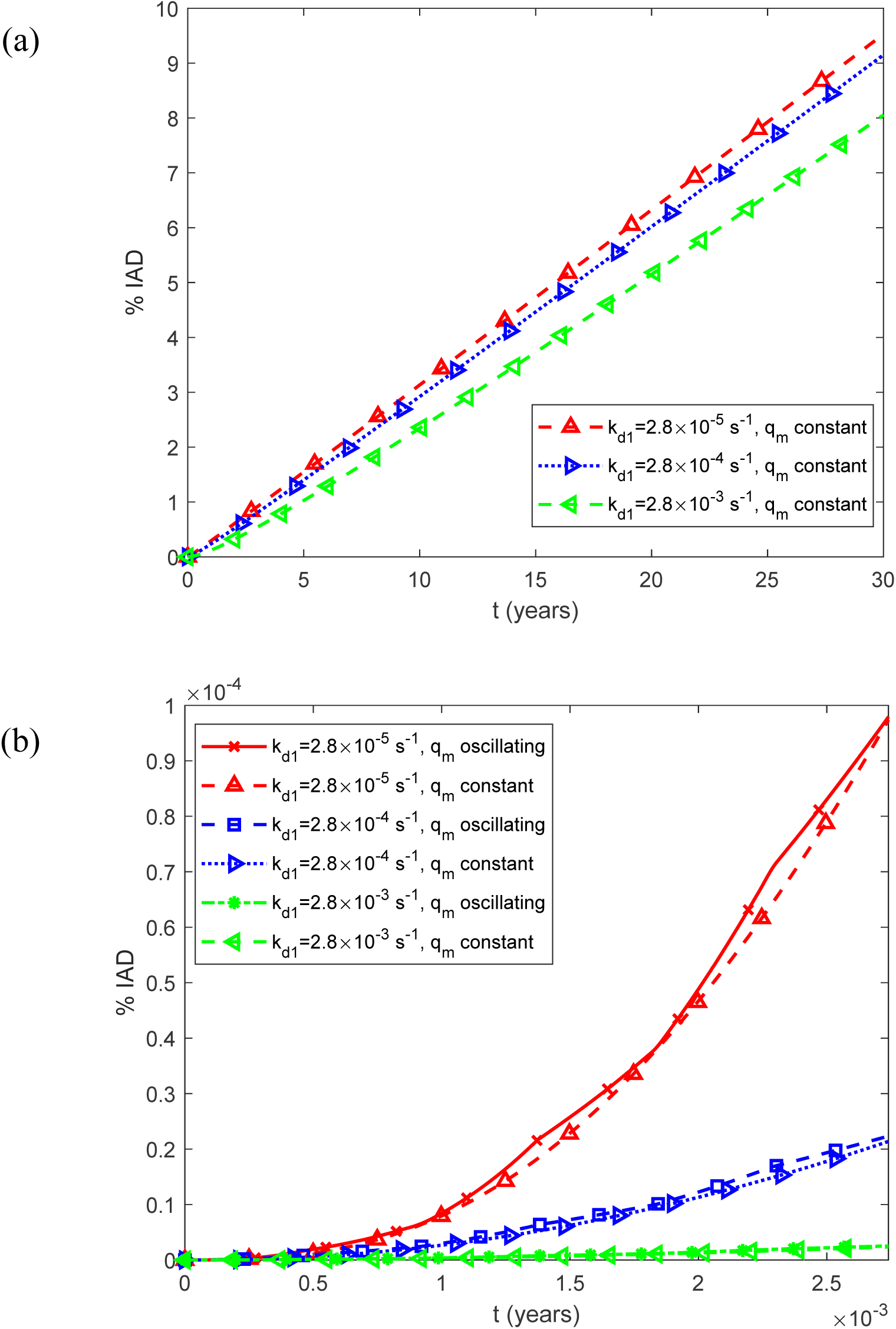
(a) Predicted percentage of the islet occupied by IADs, interpreted as the expected histological area fraction under the stereological assumption described in Section 2.2, as a function of time, %IADs, for a constant (time-averaged) monomer release rate of 1.25*q_m_*_0_ . (b) As in (a), but restricted to the time interval [0, 0.0027 years], equivalent to 1 day, illustrating the effect of oscillatory monomer release on the short-term dynamics of amyloid deposition. Results are shown for three representative values of the rate constant for the spontaneous dissociation of IAPP oligomers into constituent monomers, *k_d_*_1_.

**Fig. S22.**
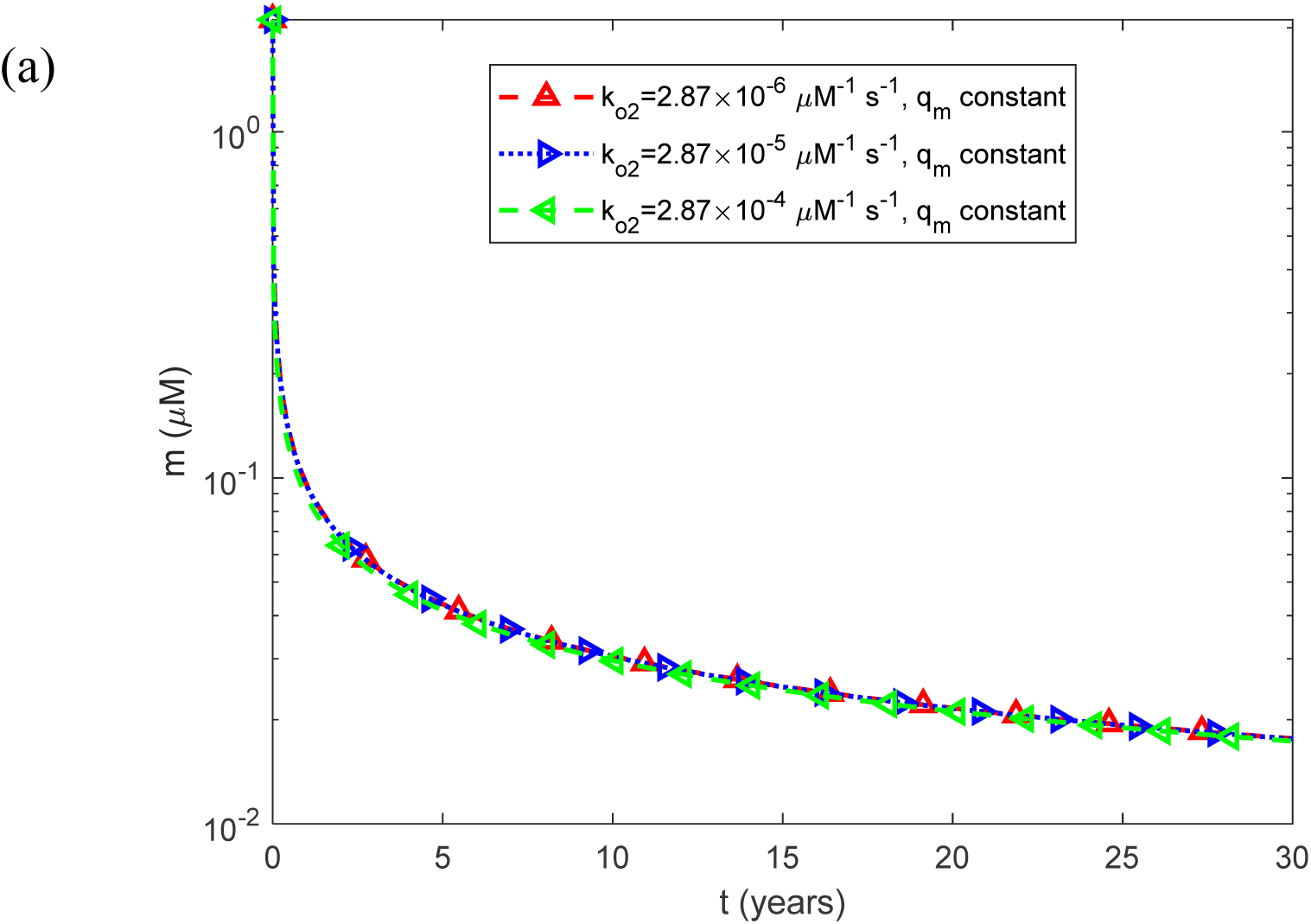
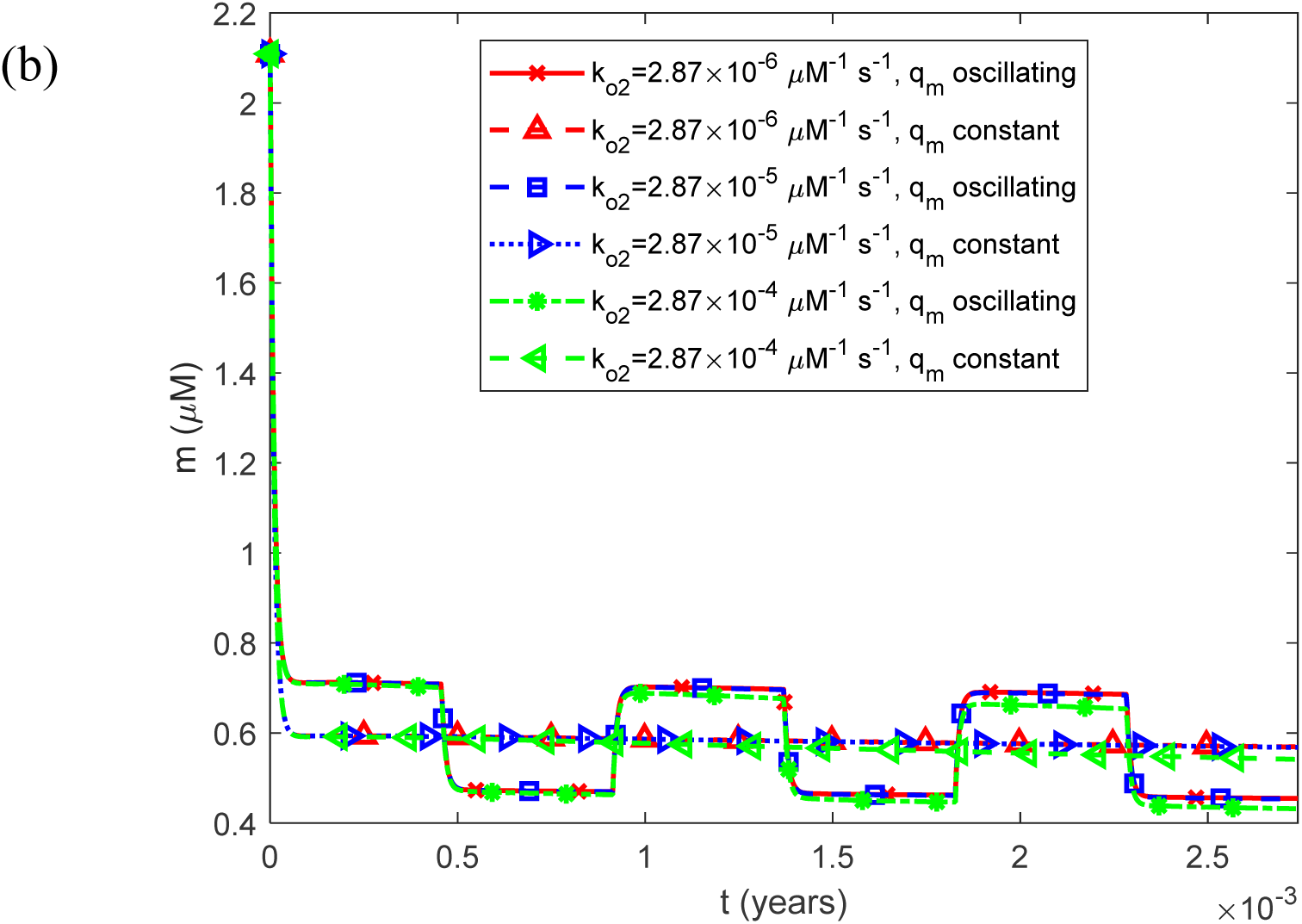
(a) Molar concentration of free IAPP monomers as a function of time, *m*(*t*) , for a constant (time-averaged) monomer release rate of 1.25*q_m_*_0_ . (b) As in (a), but restricted to the time interval [0, 0.0027 years], equivalent to 1 day, illustrating the effect of oscillatory monomer release on the short-term dynamics of the free monomer concentration. Results are shown for three representative values of the rate constant for oligomer-surface-catalyzed monomer-to-oligomer conversion, *k_o_*_2_ .

**Fig. S23.**
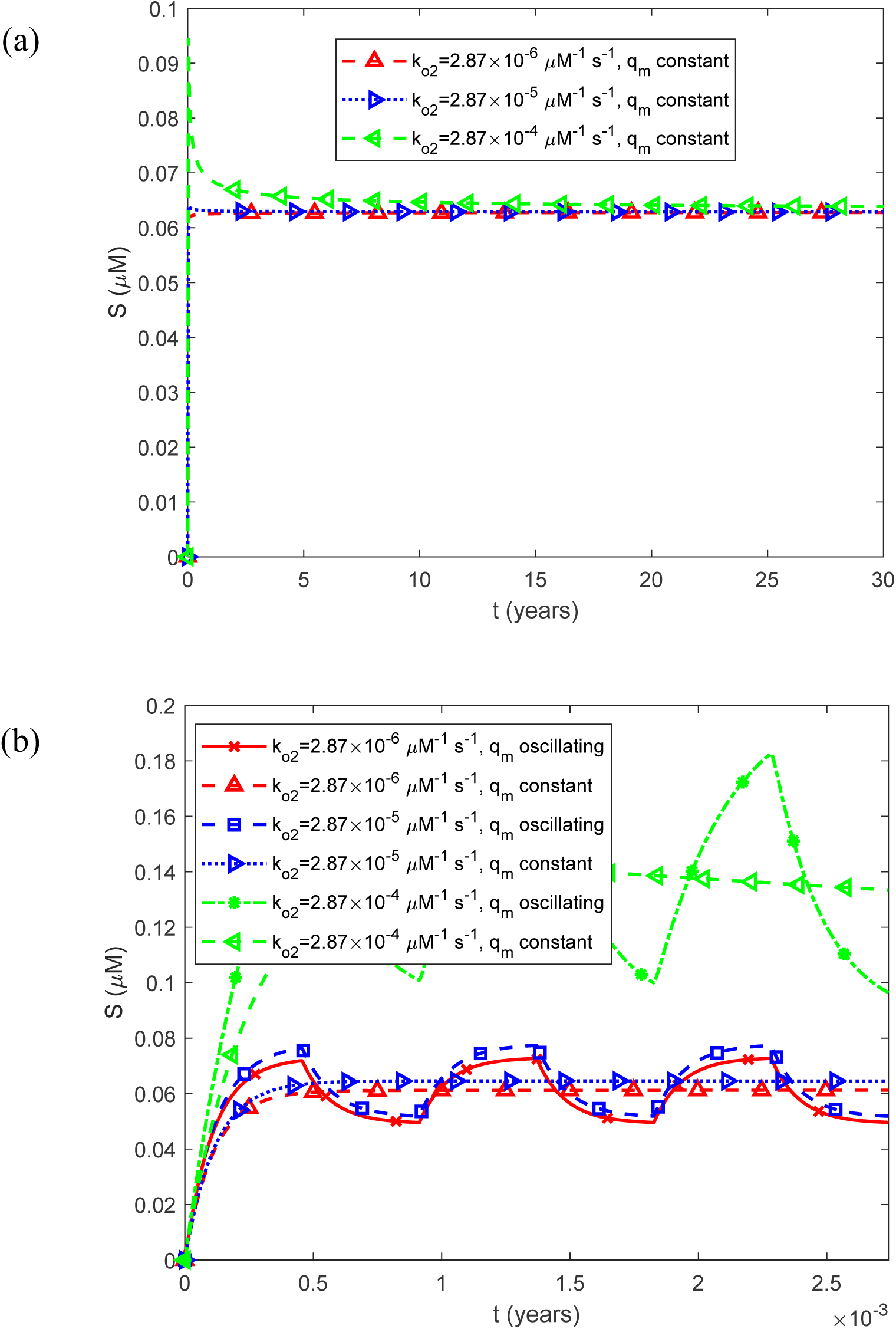
(a) Molar concentration of free IAPP oligomers as a function of time, *S*(*t*) , for a constant (time-averaged) monomer release rate of 1.25*q_m_*_0_ . (b) As in (a), but restricted to the time interval [0, 0.0027 years], equivalent to 1 day, illustrating the effect of oscillatory monomer release on the short-term dynamics of the free oligomer concentration. Results are shown for three representative values of the rate constant for oligomer-surface-catalyzed monomer-to-oligomer conversion, *k_o_*_2_ .

**Fig. S24.**
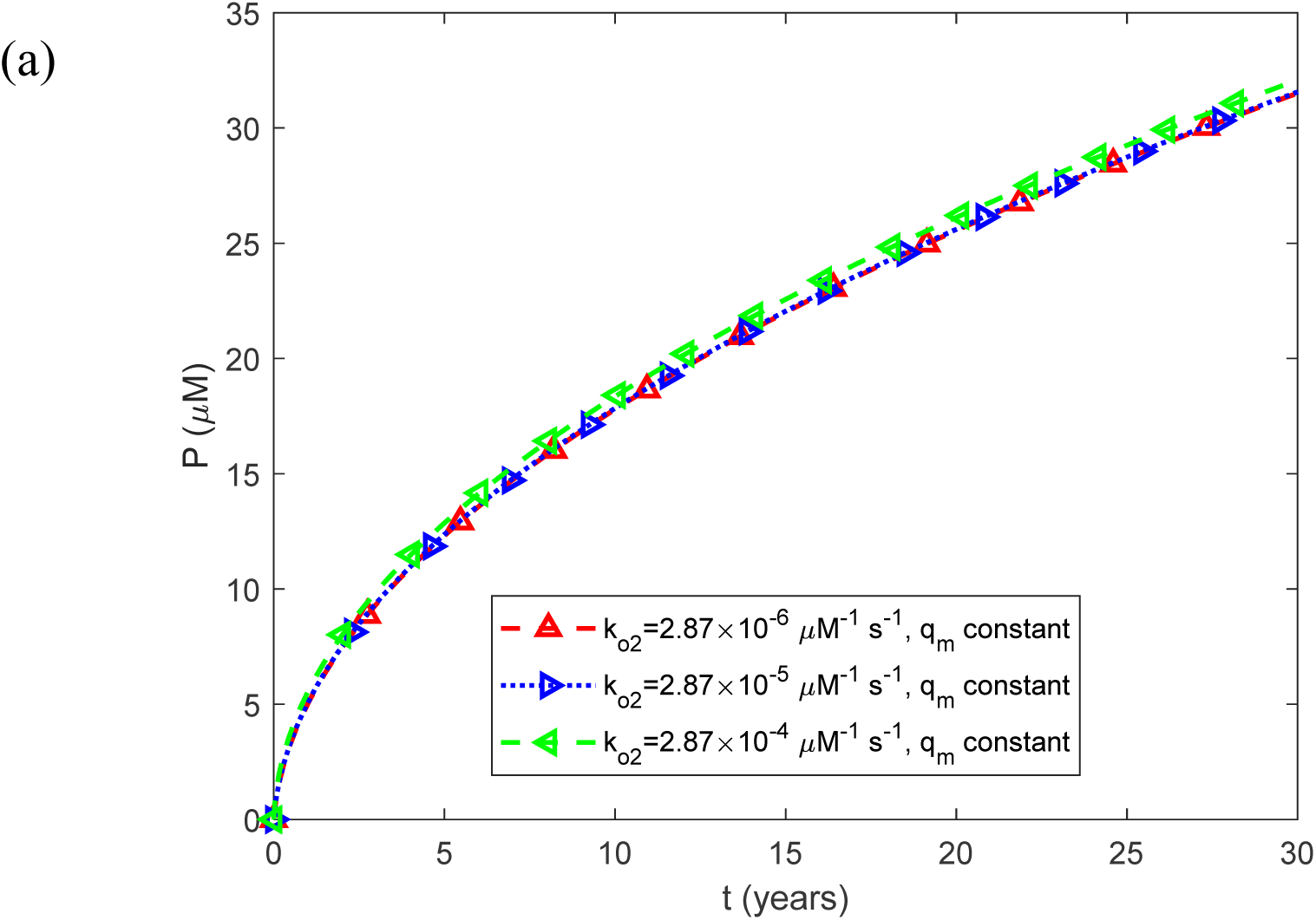
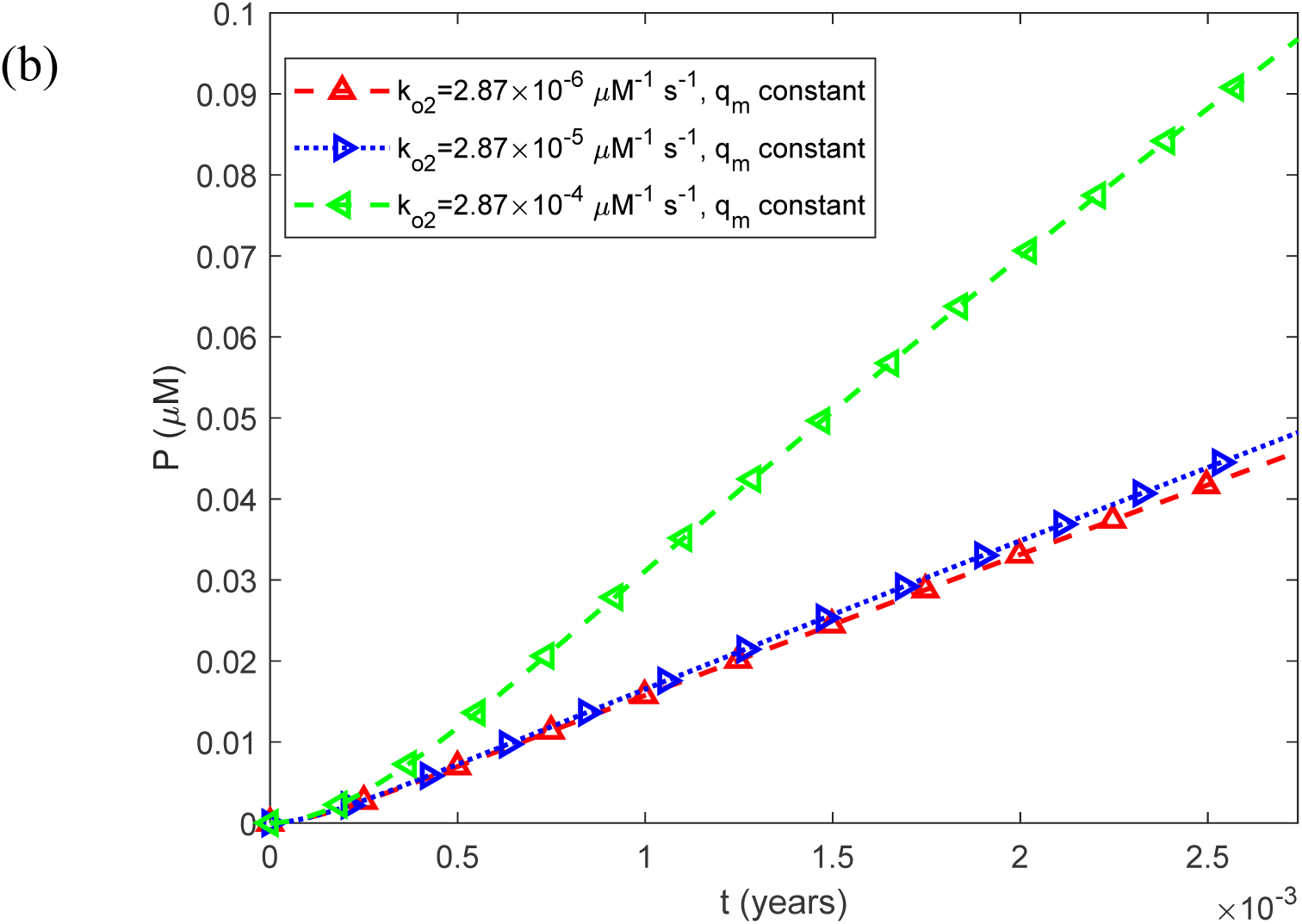
(a) Fibril particle concentration *P*(*t*) , i.e., the molar concentration of fibrils irrespective of length, as a function of time for a constant (time-averaged) monomer release rate of 1.25*q_m_*_0_ . (b) As in (a), but restricted to the time interval [0, 0.0027 years], equivalent to 1 day, illustrating the effect of oscillatory monomer release on the short-term dynamics of the fibril particle concentration. Results are shown for three representative values of the rate constant for oligomer-surface-catalyzed monomer-to-oligomer conversion, *k_o_*_2_ .

**Fig. S25.**
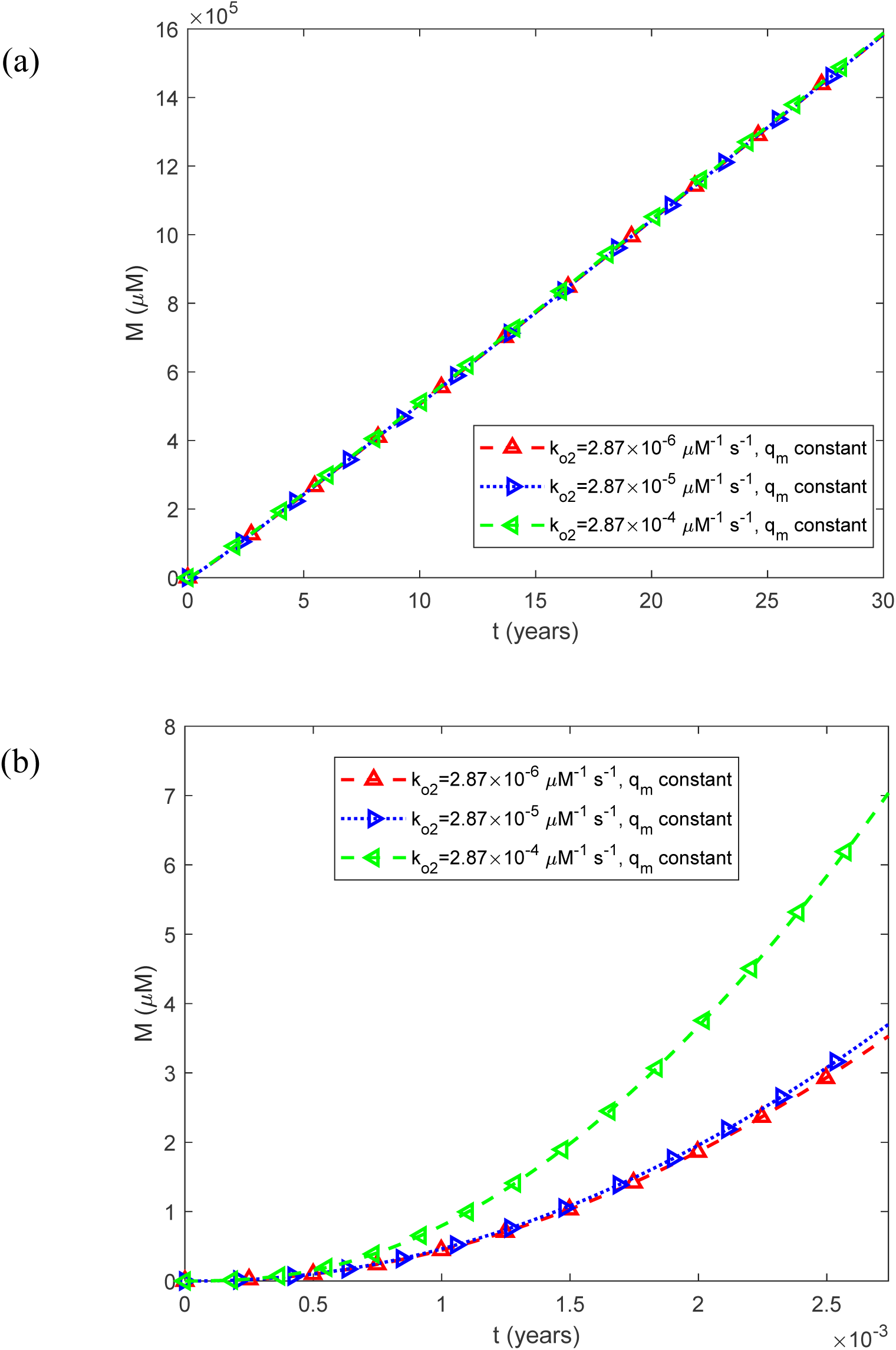
(a) Fibril mass concentration *M* (*t*) , expressed as the molar concentration of IAPP monomer equivalents incorporated into all fibrils, as a function of time for a constant (time-averaged) monomer release rate of 1.25*q_m_*_0_ . (b) As in (a), but restricted to the time interval [0, 0.0027 years], equivalent to 1 day, illustrating the effect of oscillatory monomer release on the short-term dynamics of total fibril mass. Results are shown for three representative values of the rate constant for oligomer-surface-catalyzed monomer-to-oligomer conversion, *k_o_*_2_ .

**Fig. S26.**
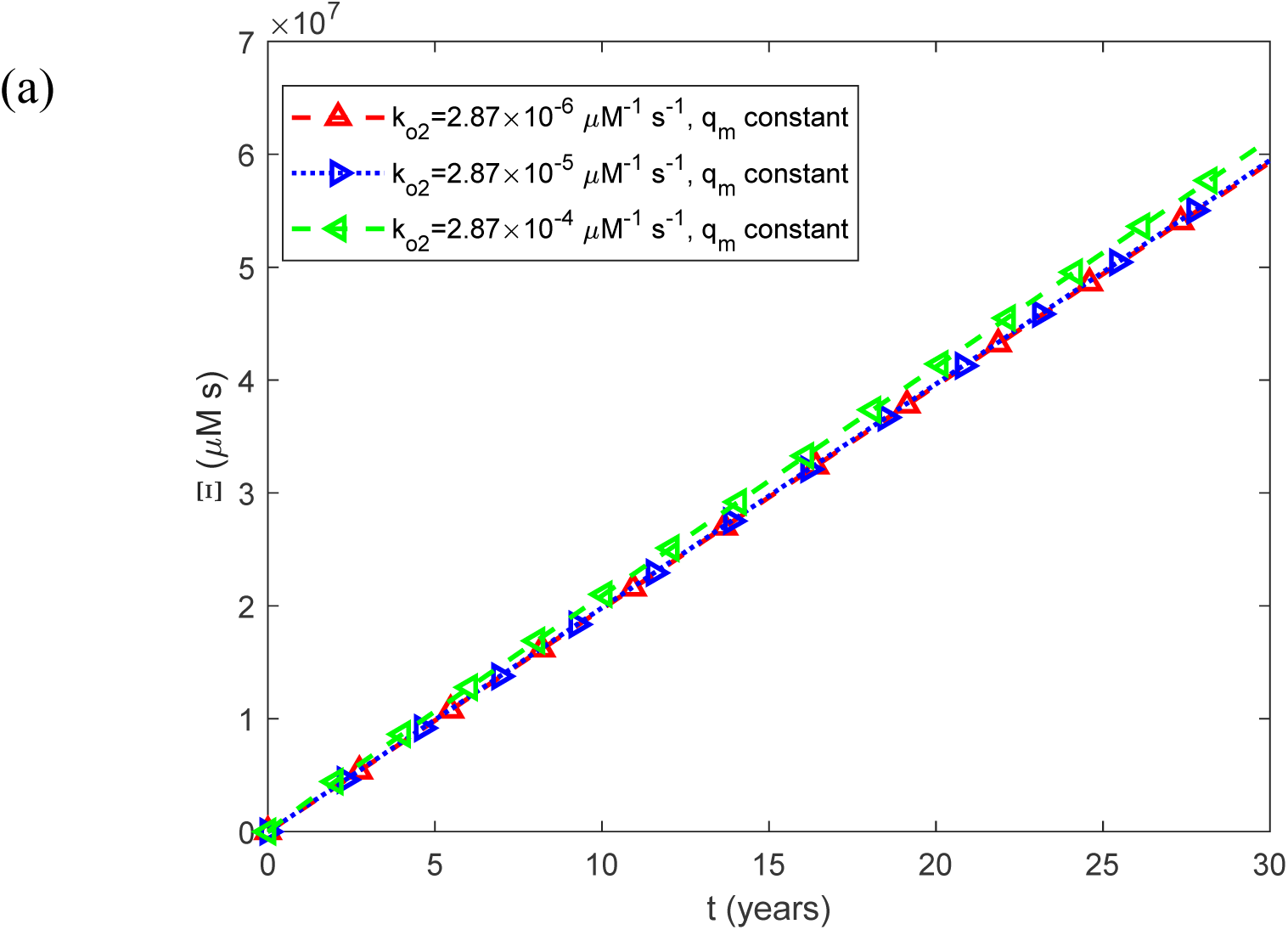
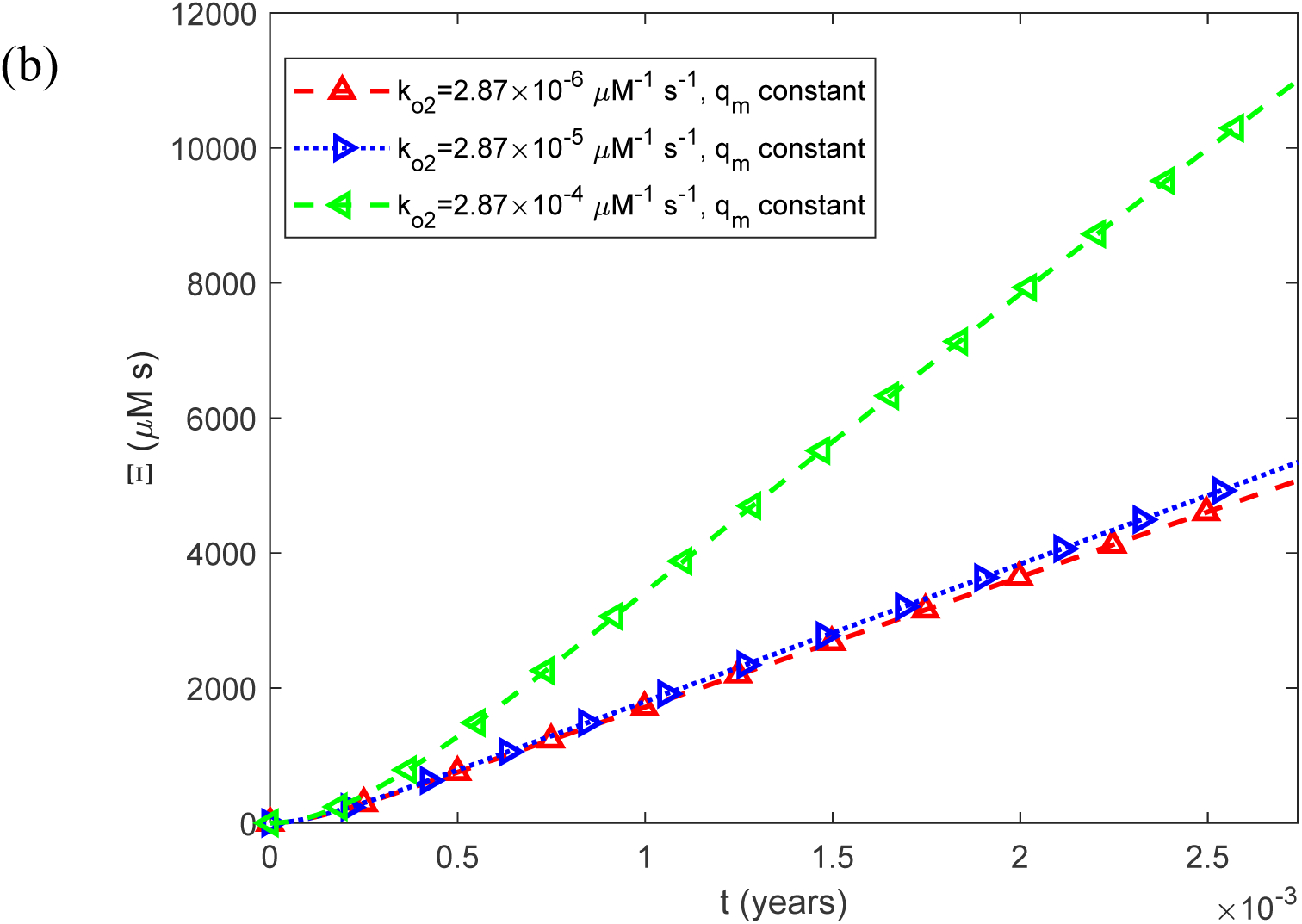
(a) Accumulated cytotoxicity (integrated IAPP oligomer exposure) as a function of time, Ξ(*t*) , for a constant (time-averaged) monomer release rate of 1.25*q_m_*_0_ . (b) As in (a), but restricted to the time interval [0, 0.0027 years], equivalent to 1 day, illustrating the effect of oscillatory monomer release on the short-term dynamics of accumulated cytotoxicity. Results are shown for three representative values of the rate constant for oligomer-surface-catalyzed monomer-to-oligomer conversion, *k_o_*_2_ .

**Fig. S27.**
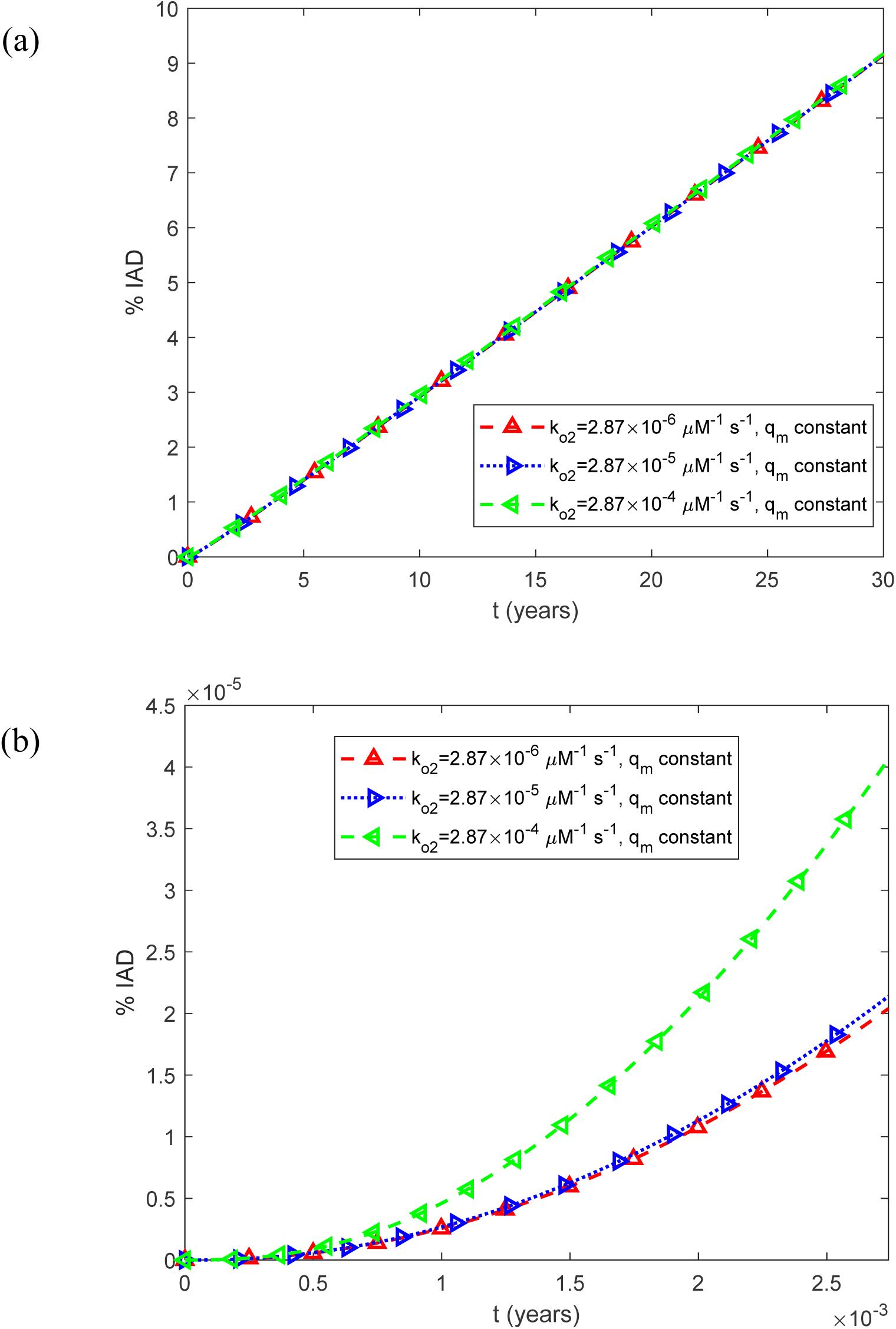
(a) Predicted percentage of the islet occupied by IADs, interpreted as the expected histological area fraction under the stereological assumption described in Section 2.2, as a function of time, %IADs, for a constant (time-averaged) monomer release rate of 1.25*q_m_*_0_ . (b) As in (a), but restricted to the time interval [0, 0.0027 years], equivalent to 1 day, illustrating the effect of oscillatory monomer release on the short-term dynamics of amyloid deposition. Results are shown for three representative values of the rate constant for oligomer-surface-catalyzed monomer-to-oligomer conversion, *k_o_*_2_ .

**Fig. S28.**
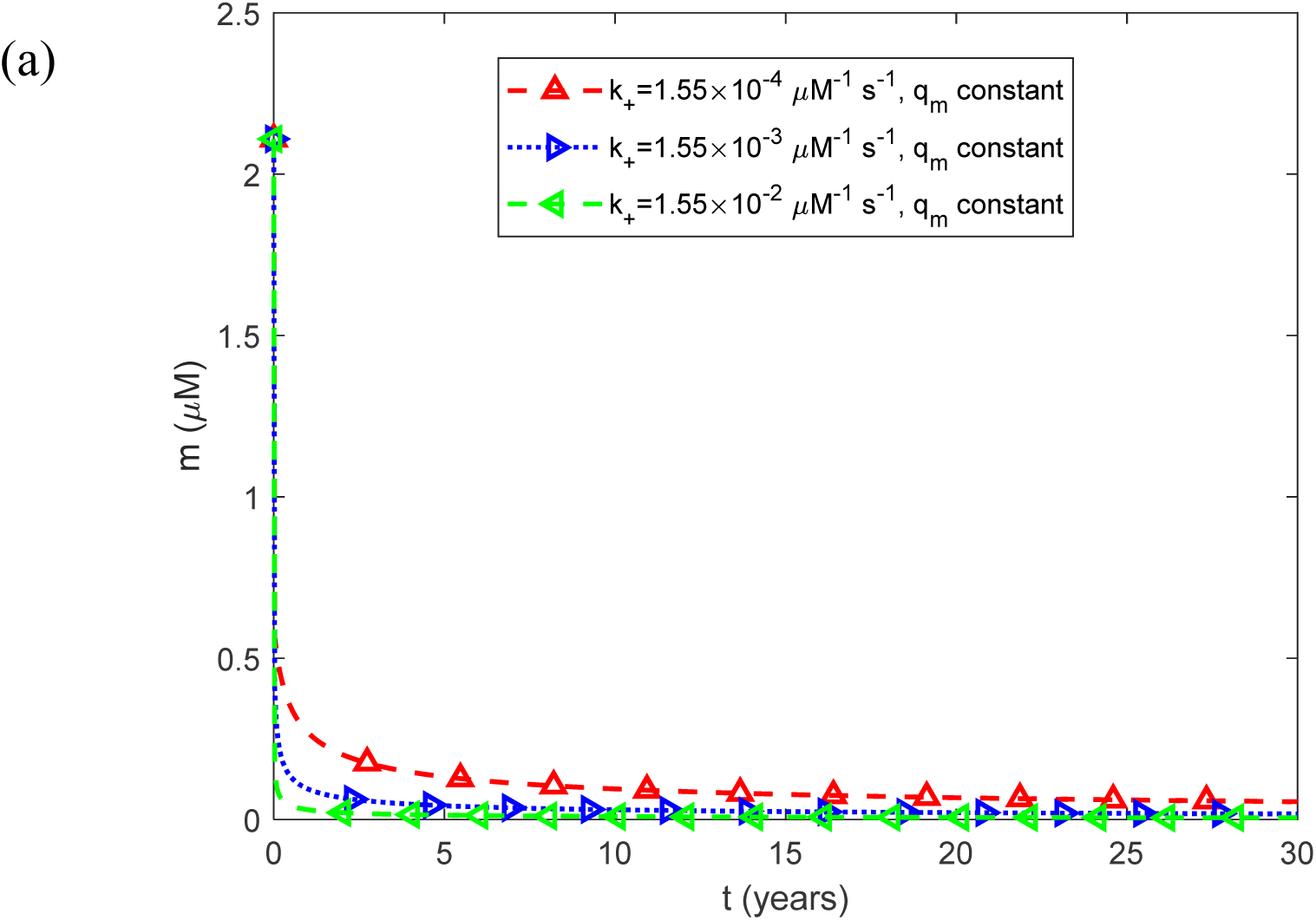
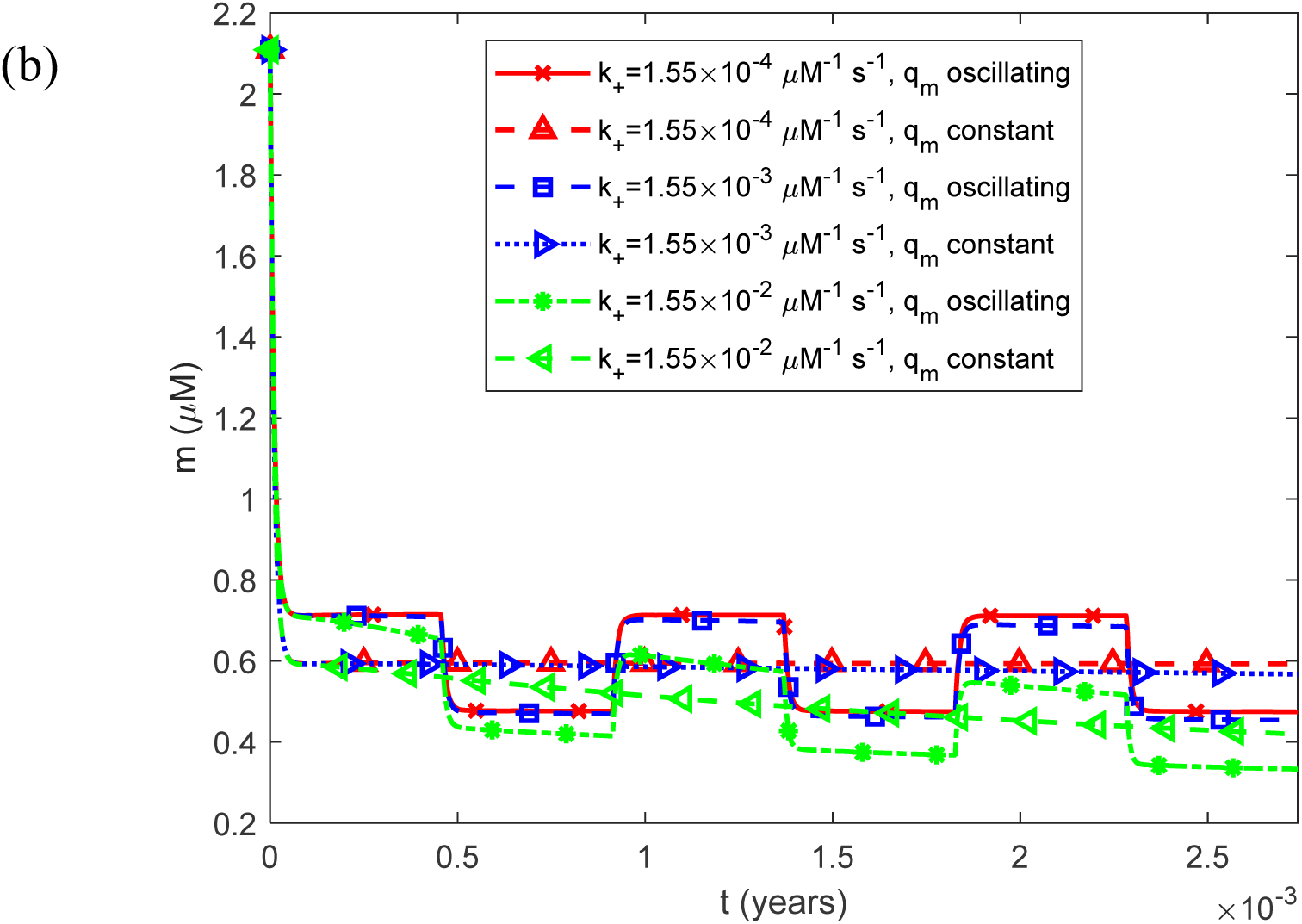
(a) Molar concentration of free IAPP monomers as a function of time, *m*(*t*) , for a constant (time-averaged) monomer release rate of 1.25*q_m_*_0_ . (b) As in (a), but restricted to the time interval [0, 0.0027 years], equivalent to 1 day, illustrating the effect of oscillatory monomer release on the short-term dynamics of the free monomer concentration. Results are shown for three representative values of the rate constant for IAPP fibril elongation by monomer addition, *k*_+_ .

**Fig. S29.**
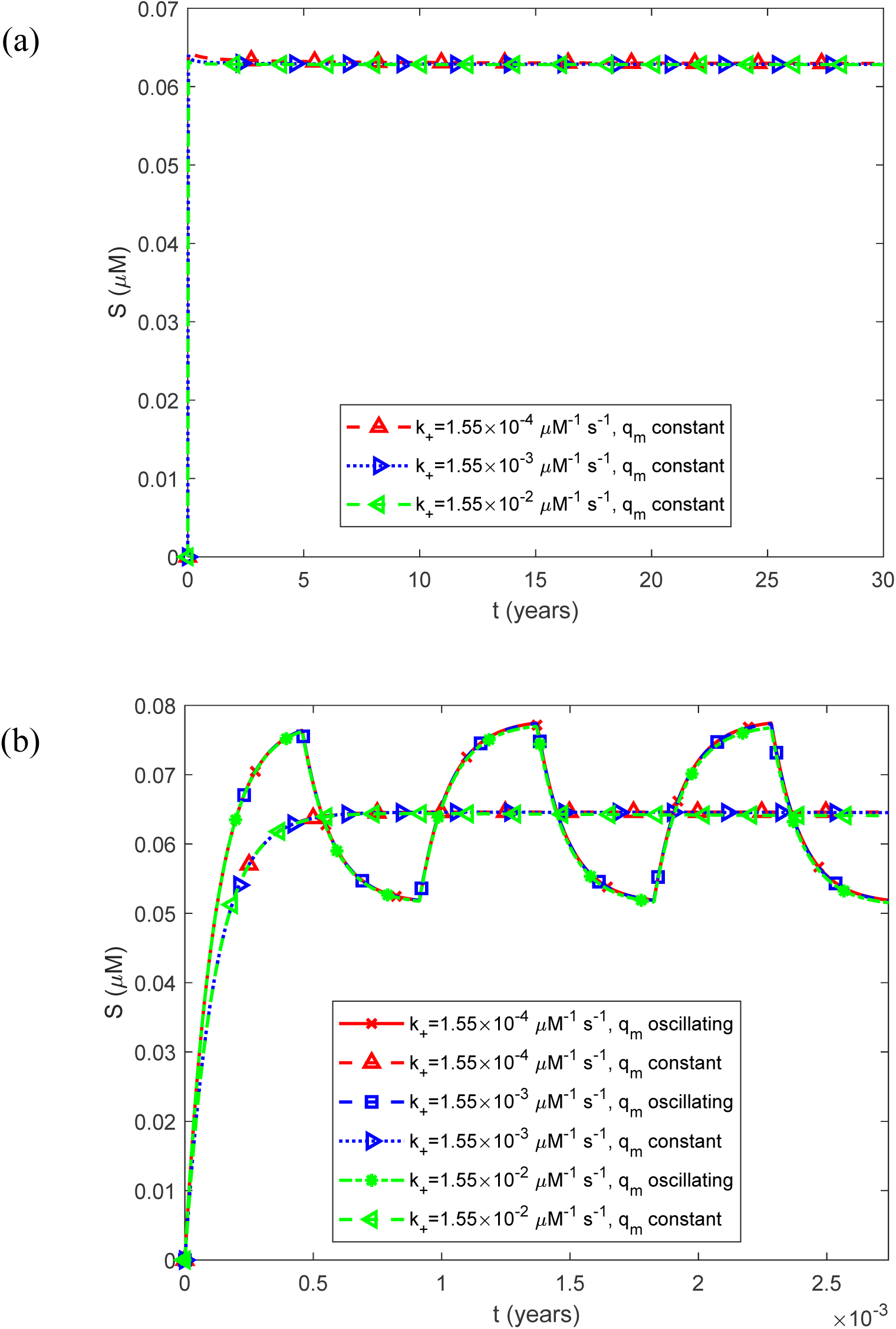
(a) Molar concentration of free IAPP oligomers as a function of time, *S*(*t*) , for a constant (time-averaged) monomer release rate of 1.25*q_m_*_0_ . (b) As in (a), but restricted to the time interval [0, 0.0027 years], equivalent to 1 day, illustrating the effect of oscillatory monomer release on the short-term dynamics of the free oligomer concentration. Results are shown for three representative values of the rate constant for IAPP fibril elongation by monomer addition, *k*_+_ .

**Fig. S30.**
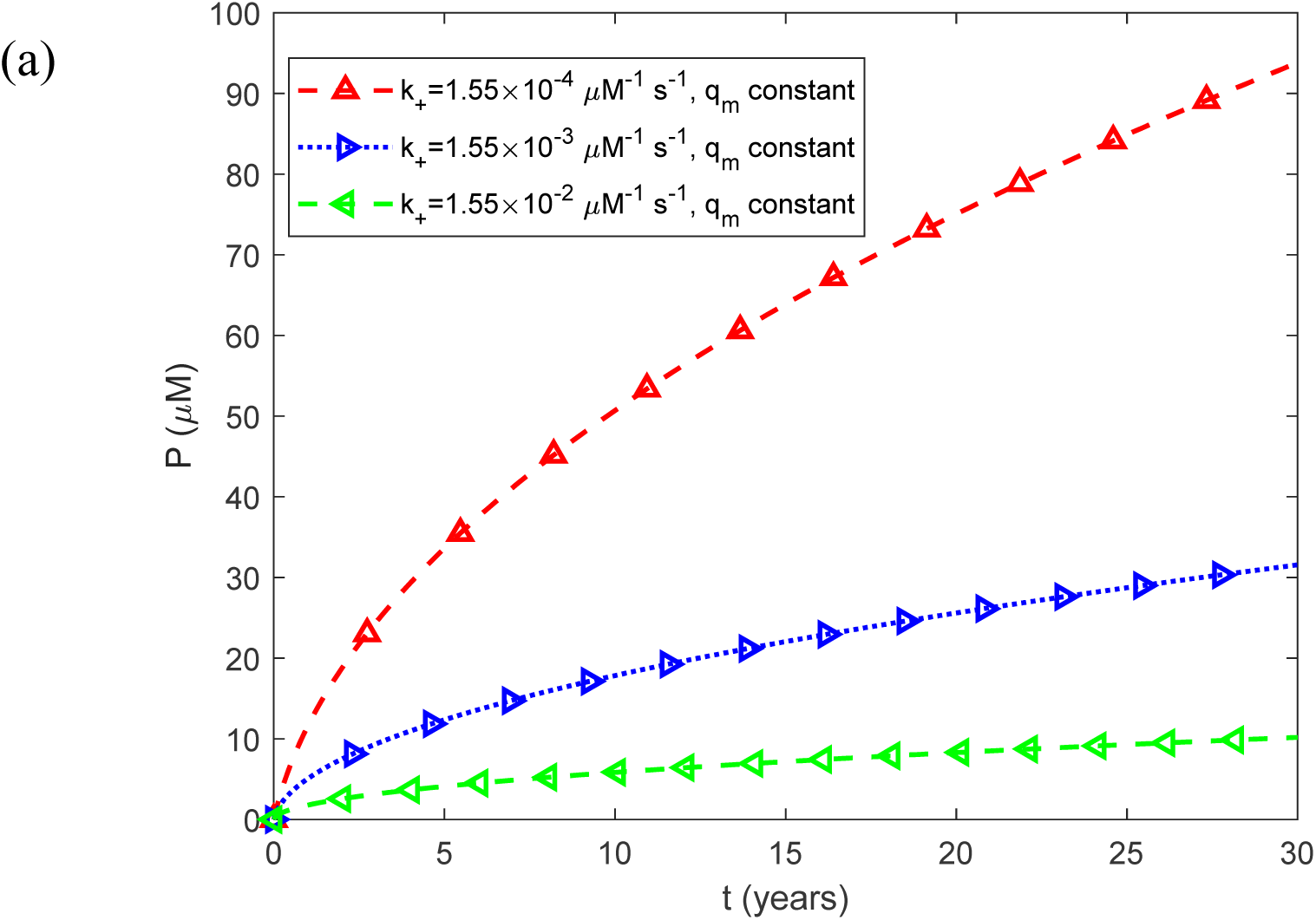
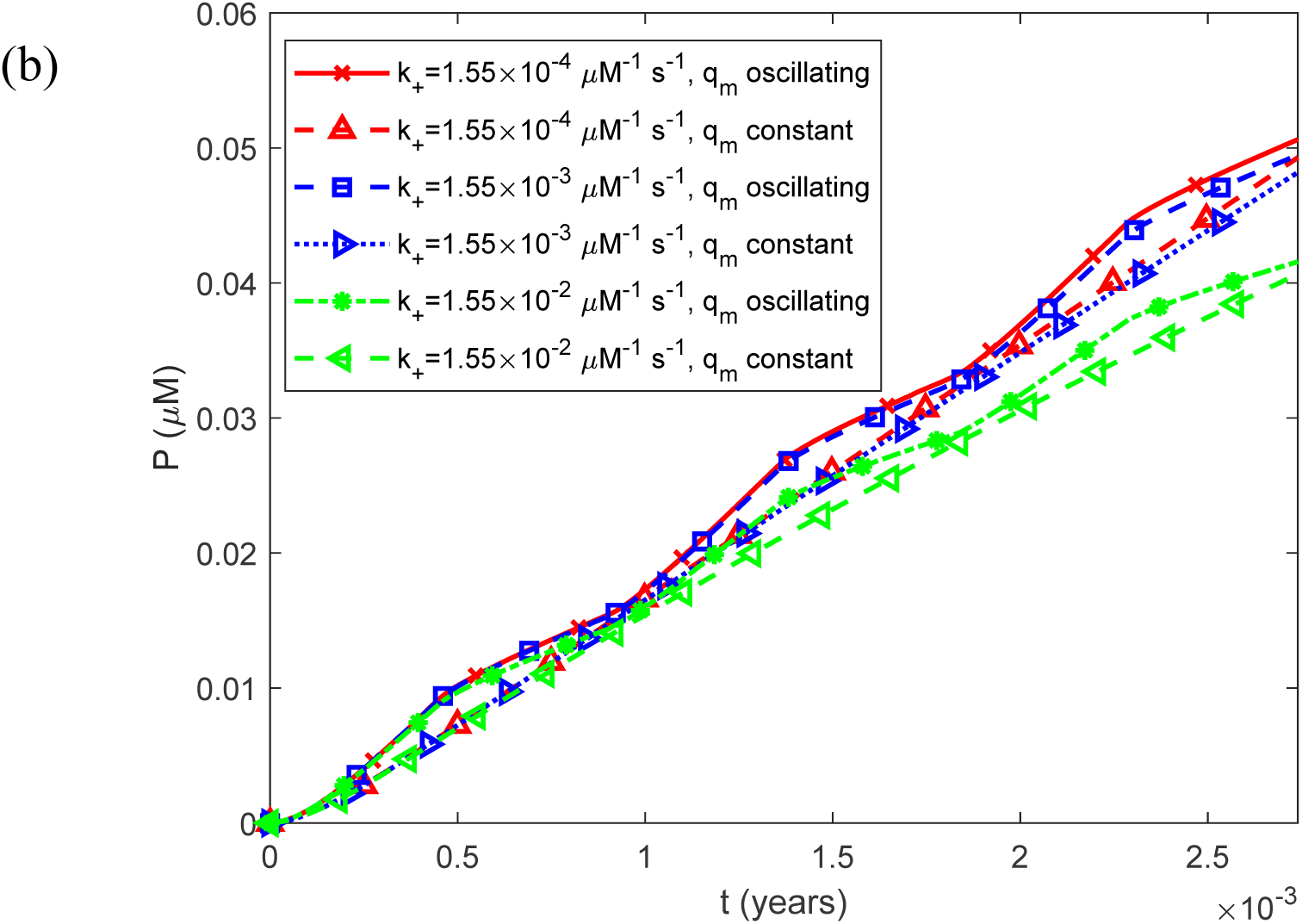
(a) Fibril particle concentration *P*(*t*) , i.e., the molar concentration of fibrils irrespective of length, as a function of time for a constant (time-averaged) monomer release rate of 1.25*q_m_*_0_ . (b) As in (a), but restricted to the time interval [0, 0.0027 years], equivalent to 1 day, illustrating the effect of oscillatory monomer release on the short-term dynamics of the fibril particle concentration. Results are shown for three representative values of the rate constant for IAPP fibril elongation by monomer addition, *k*_+_ .

**Fig. S31.**
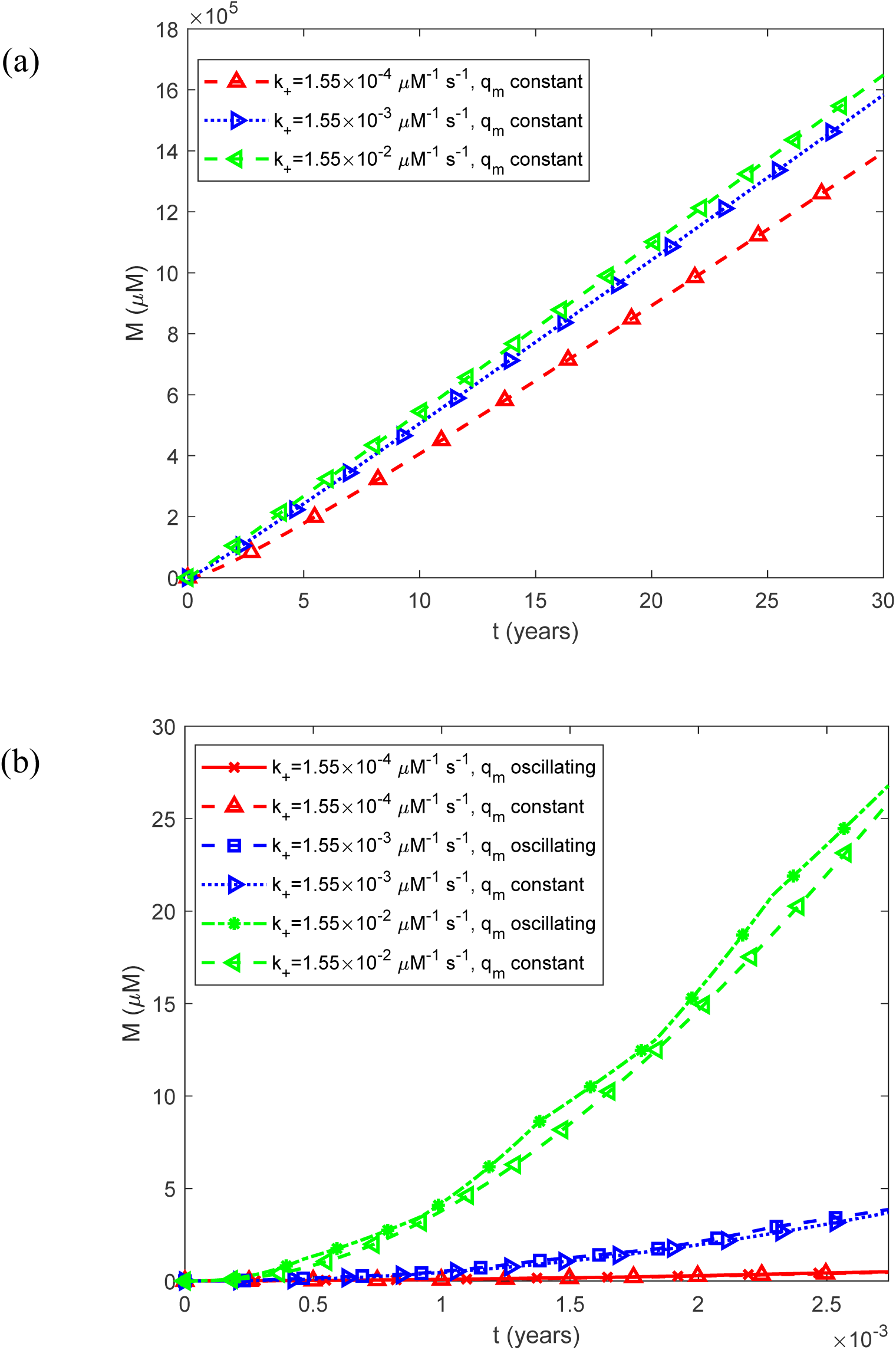
(a) Fibril mass concentration *M* (*t*) , expressed as the molar concentration of IAPP monomer equivalents incorporated into all fibrils, as a function of time for a constant (time-averaged) monomer release rate of 1.25*q_m_*_0_ . (b) As in (a), but restricted to the time interval [0, 0.0027 years], equivalent to 1 day, illustrating the effect of oscillatory monomer release on the short-term dynamics of total fibril mass. Results are shown for three representative values of the rate constant for IAPP fibril elongation by monomer addition, *k*_+_ .

**Fig. S32.**
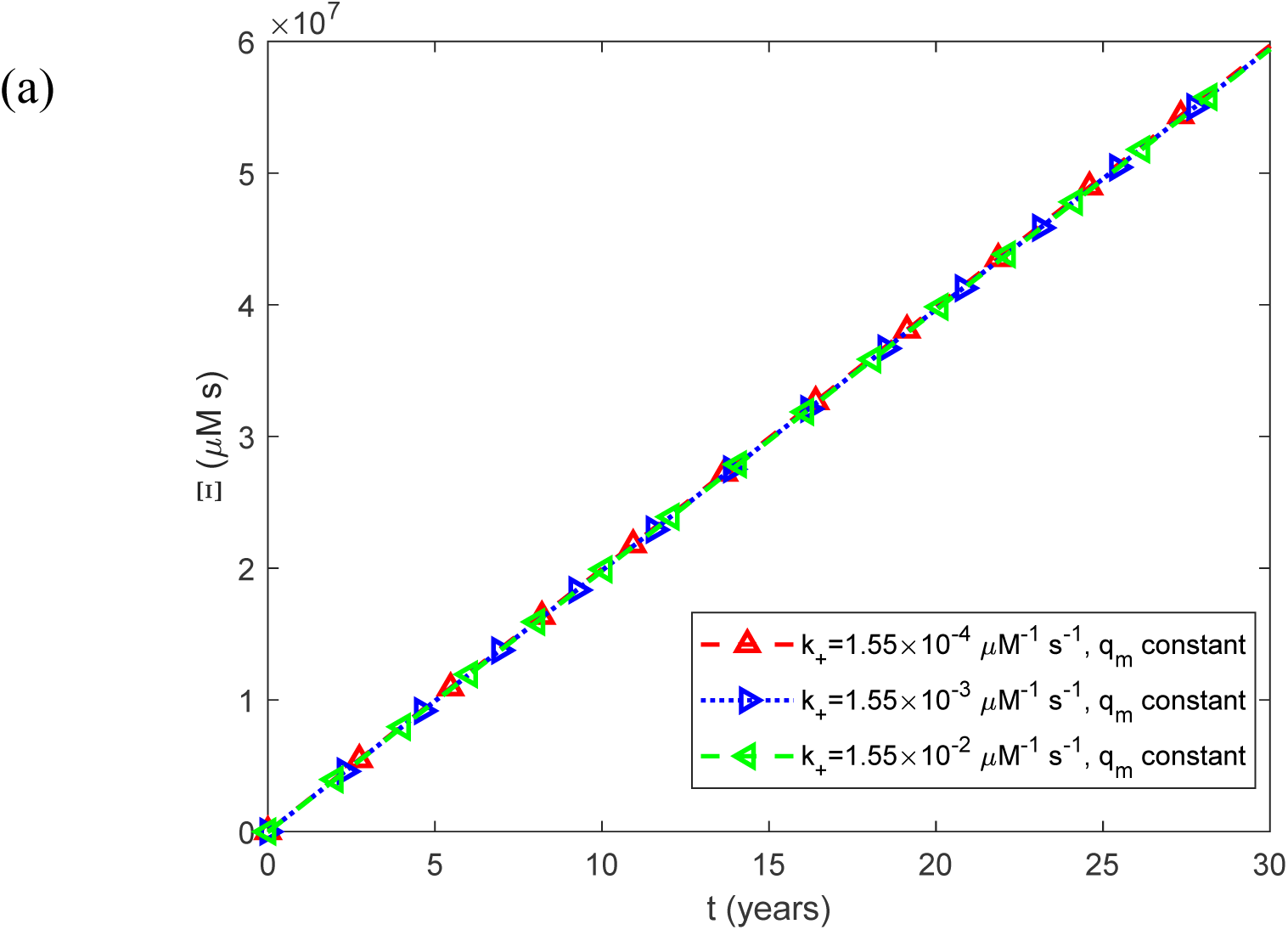
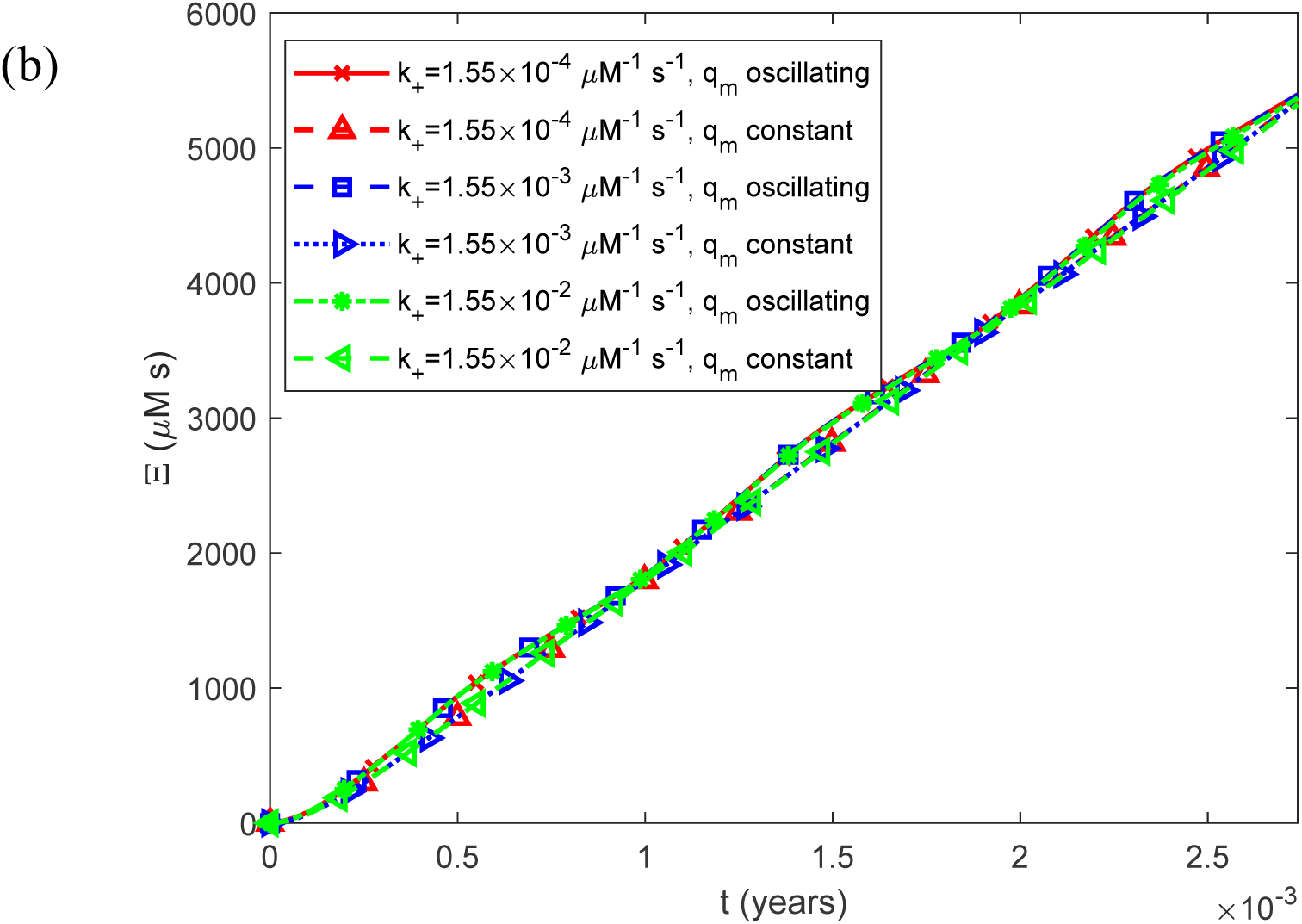
(a) Accumulated cytotoxicity (integrated IAPP oligomer exposure) as a function of time, Ξ(*t*) , for a constant (time-averaged) monomer release rate of 1.25*q_m_*_0_ . (b) As in (a), but restricted to the time interval [0, 0.0027 years], equivalent to 1 day, illustrating the effect of oscillatory monomer release on the short-term dynamics of accumulated cytotoxicity. Results are shown for three representative values of the rate constant for IAPP fibril elongation by monomer addition, *k*_+_ .

**Fig. S33.**
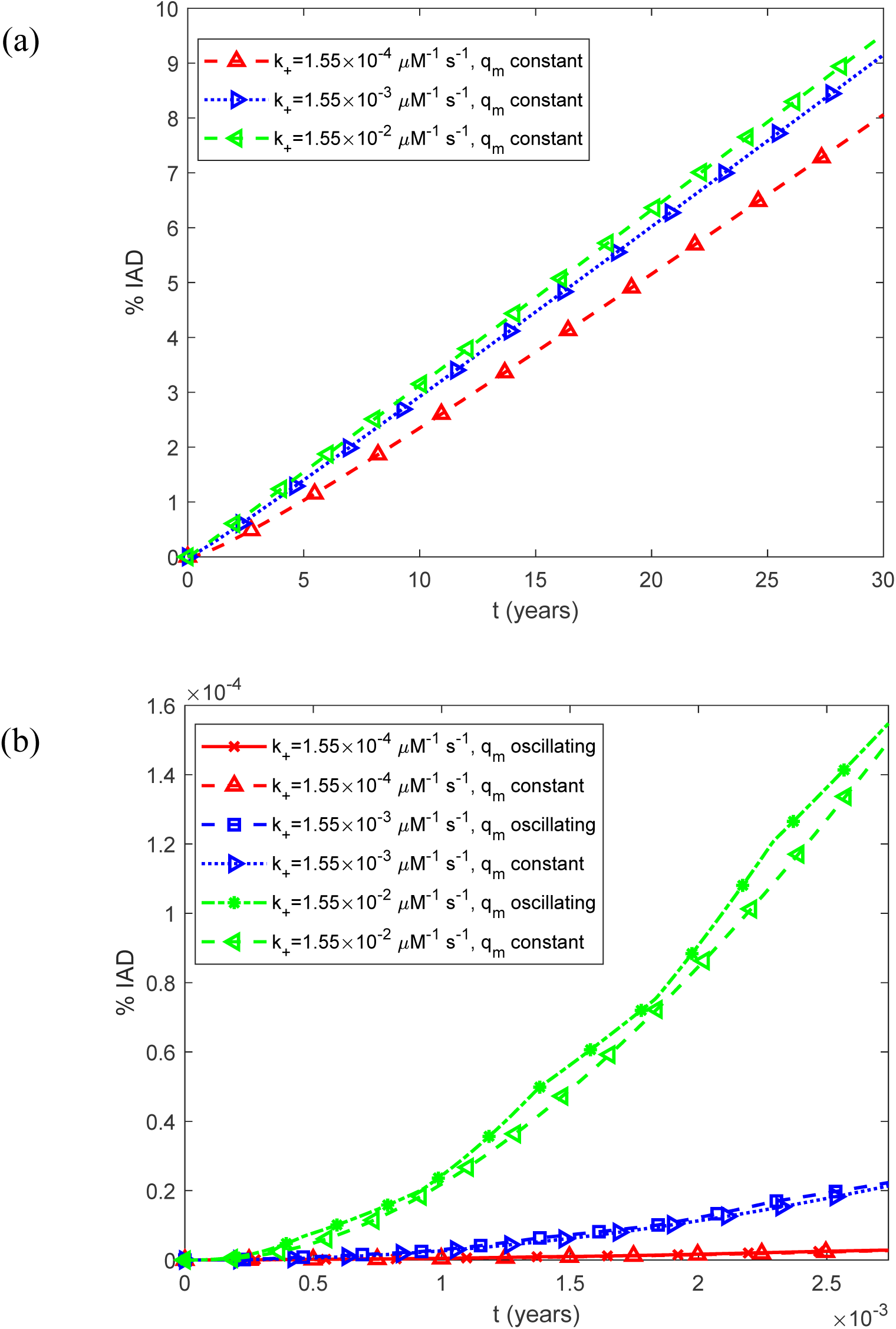
(a) Predicted percentage of the islet occupied by IADs, interpreted as the expected histological area fraction under the stereological assumption described in Section 2.2, as a function of time, %IADs, for a constant (time-averaged) monomer release rate of 1.25*q_m_*_0_ . (b) As in (a), but restricted to the time interval [0, 0.0027 years], equivalent to 1 day, illustrating the effect of oscillatory monomer release on the short-term dynamics of amyloid deposition. Results are shown for three representative values of the rate constant for IAPP fibril elongation by monomer addition, *k*_+_ .

**Fig. S34.**
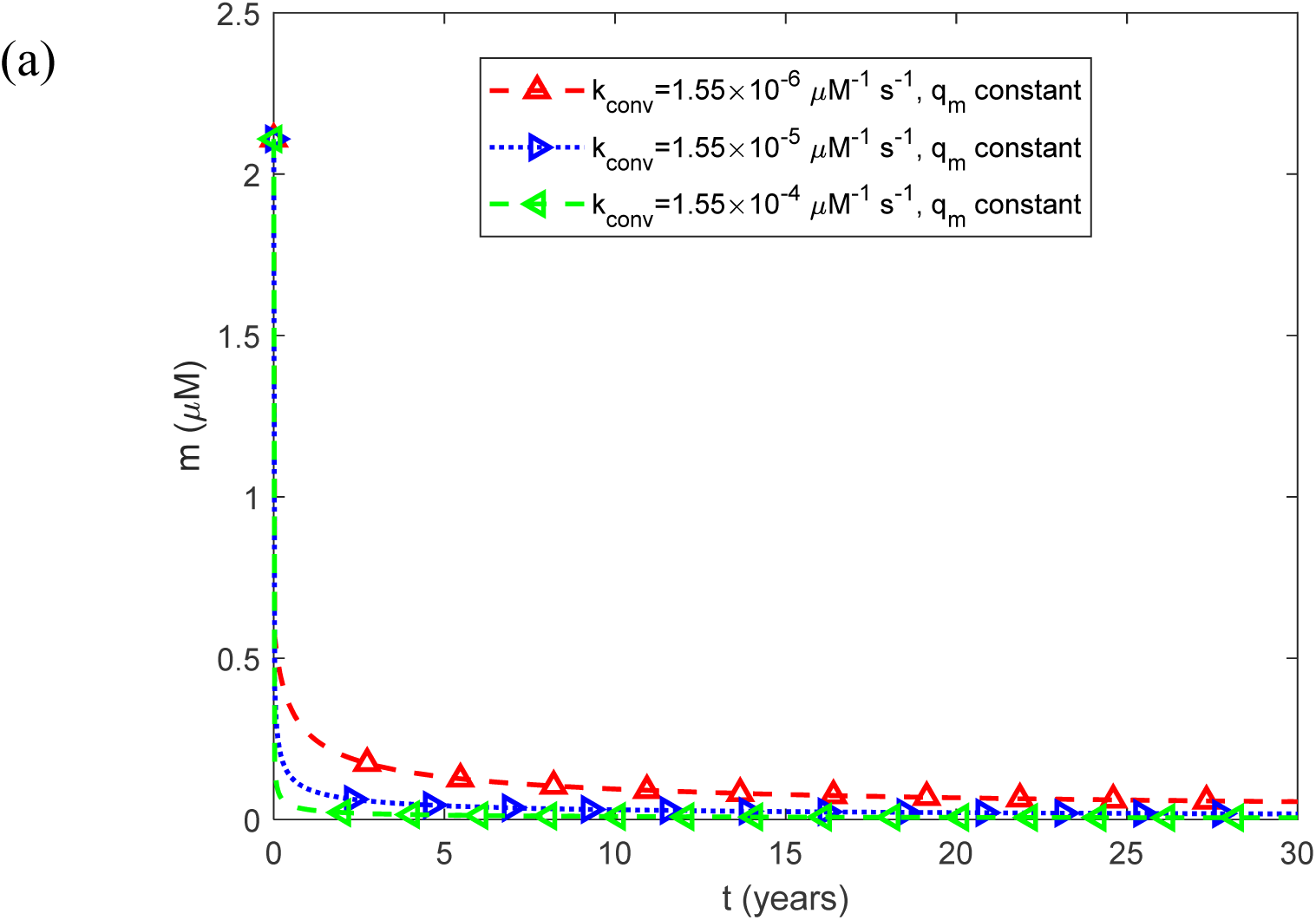
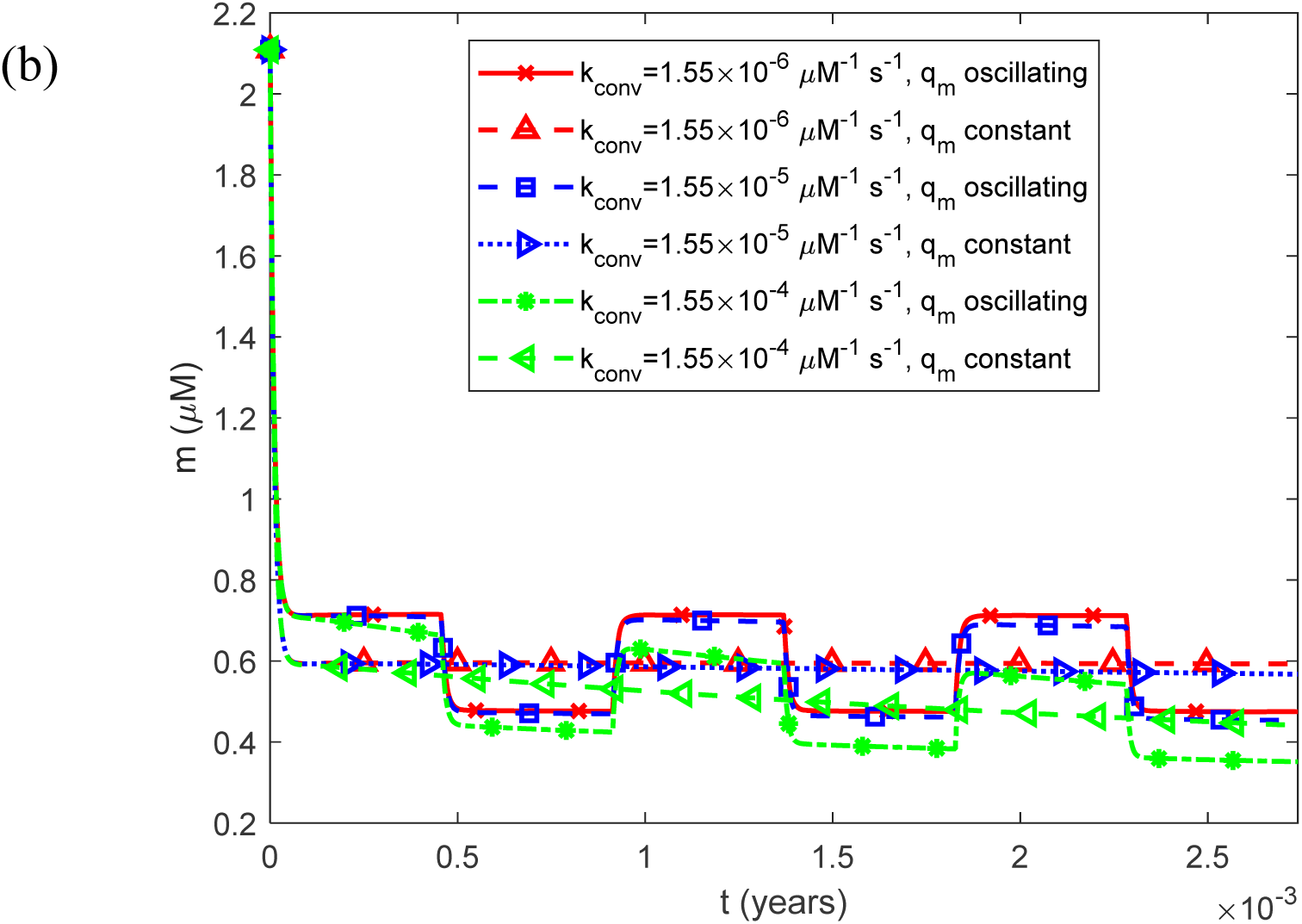
(a) Molar concentration of free IAPP monomers as a function of time, *m*(*t*) , for a constant (time-averaged) monomer release rate of 1.25*q_m_*_0_ . (b) As in (a), but restricted to the time interval [0, 0.0027 years], equivalent to 1 day, illustrating the effect of oscillatory monomer release on the short-term dynamics of the free monomer concentration. Results are shown for three representative values of the rate constant for the conversion of IAPP oligomers into elongation-competent fibrils, *k_conv_* .

**Fig. S35.**
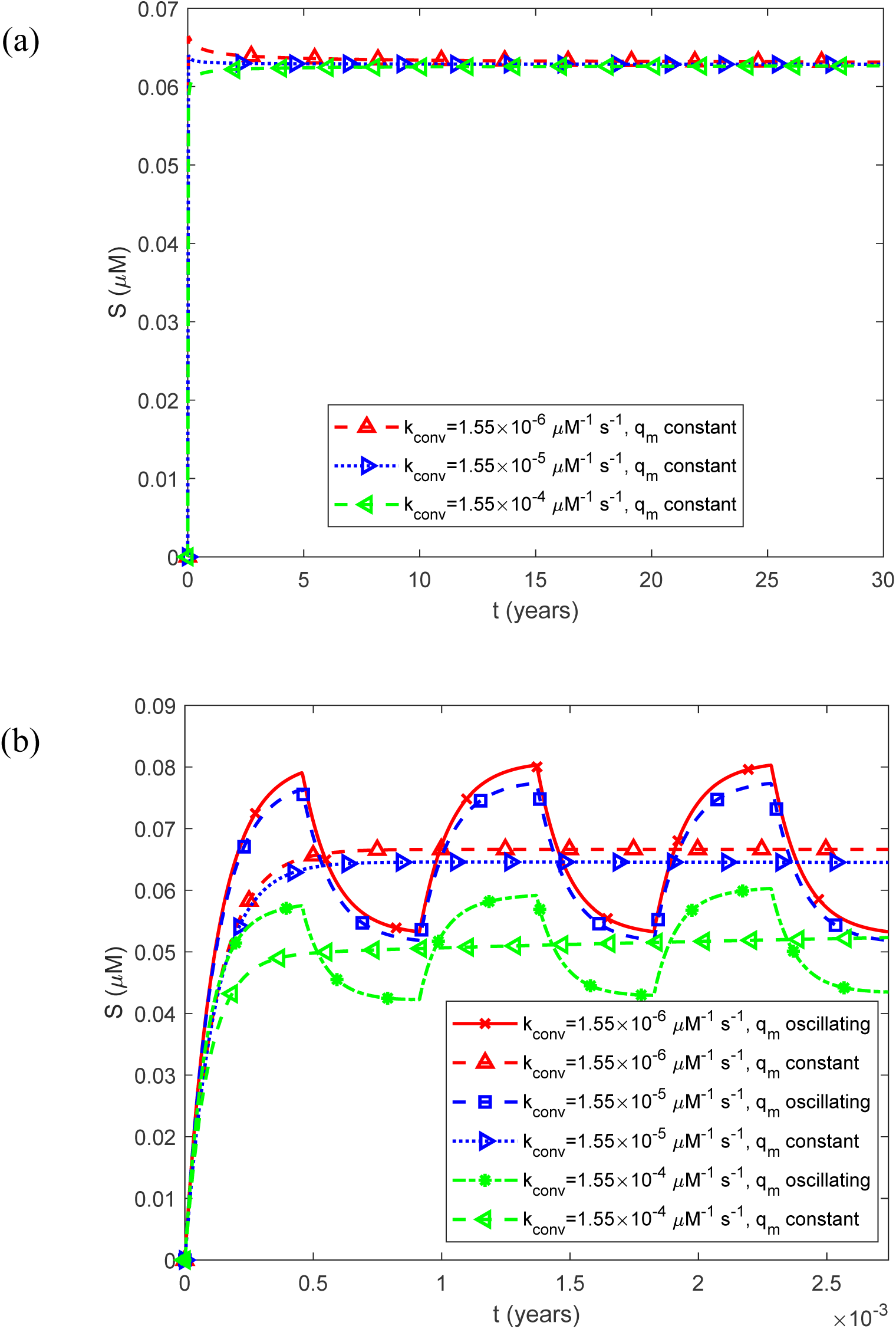
(a) Molar concentration of free IAPP oligomers as a function of time, *S*(*t*) , for a constant (time-averaged) monomer release rate of 1.25*q_m_*_0_ . (b) As in (a), but restricted to the time interval [0, 0.0027 years], equivalent to 1 day, illustrating the effect of oscillatory monomer release on the short-term dynamics of the free oligomer concentration. Results are shown for three representative values of the rate constant for the conversion of IAPP oligomers into elongation-competent fibrils, *k_conv_* .

**Fig. S36.**
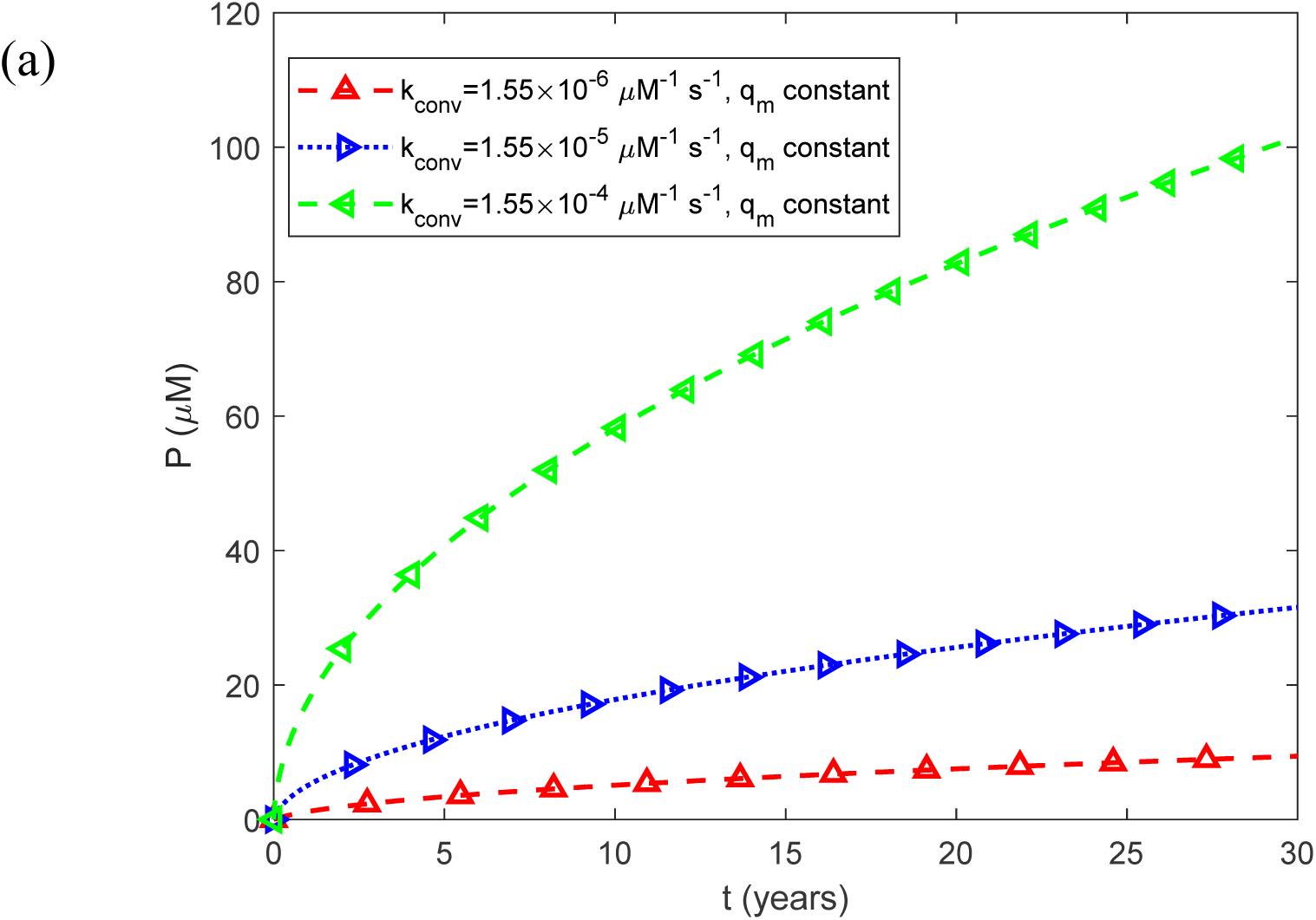
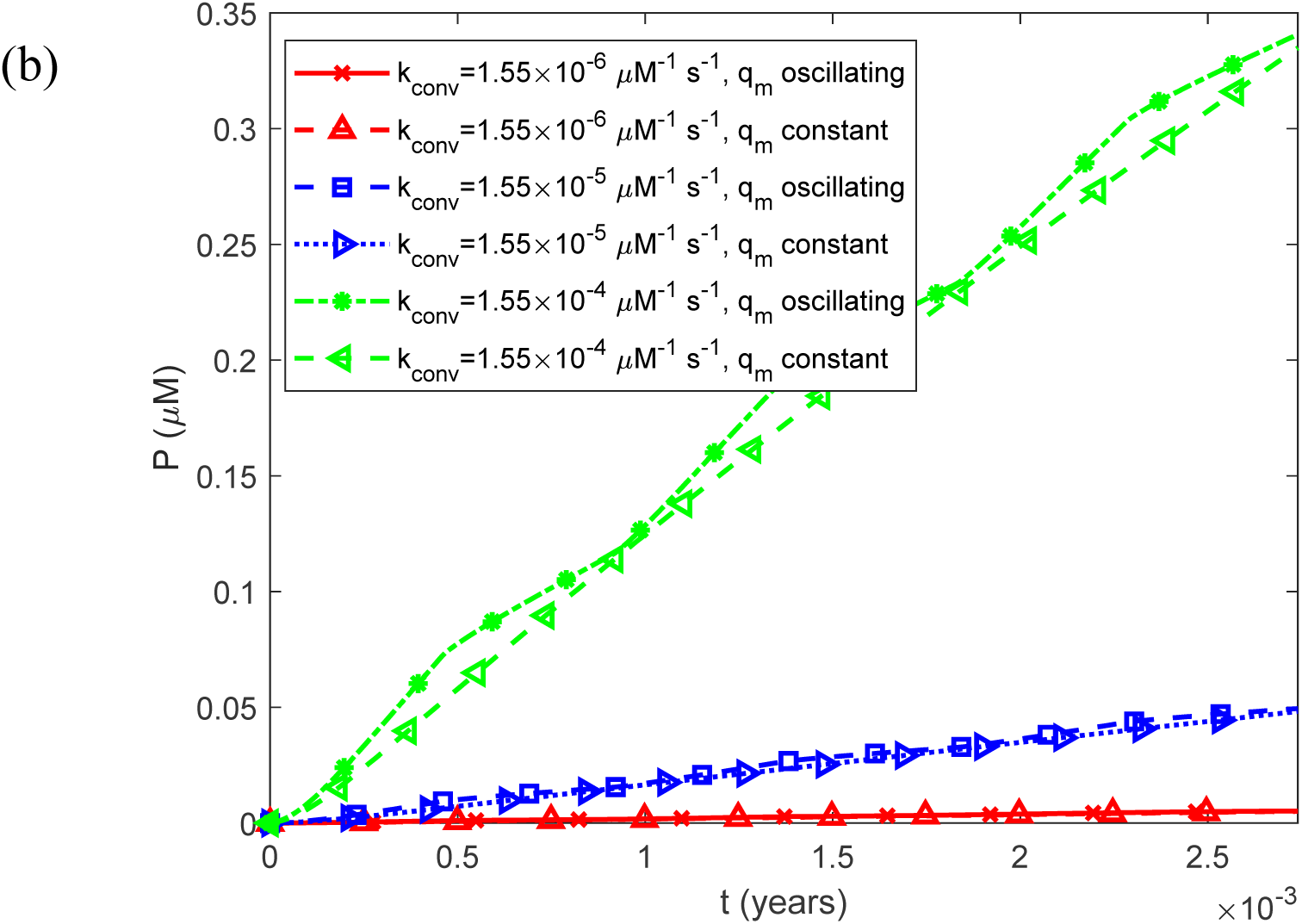
(a) Fibril particle concentration *P*(*t*) , i.e., the molar concentration of fibrils irrespective of length, as a function of time for a constant (time-averaged) monomer release rate of 1.25*q_m_*_0_ . (b) As in (a), but restricted to the time interval [0, 0.0027 years], equivalent to 1 day, illustrating the effect of oscillatory monomer release on the short-term dynamics of the fibril particle concentration. Results are shown for three representative values of the rate constant for the conversion of IAPP oligomers into elongation-competent fibrils, *k_conv_*.

**Fig. S37.**
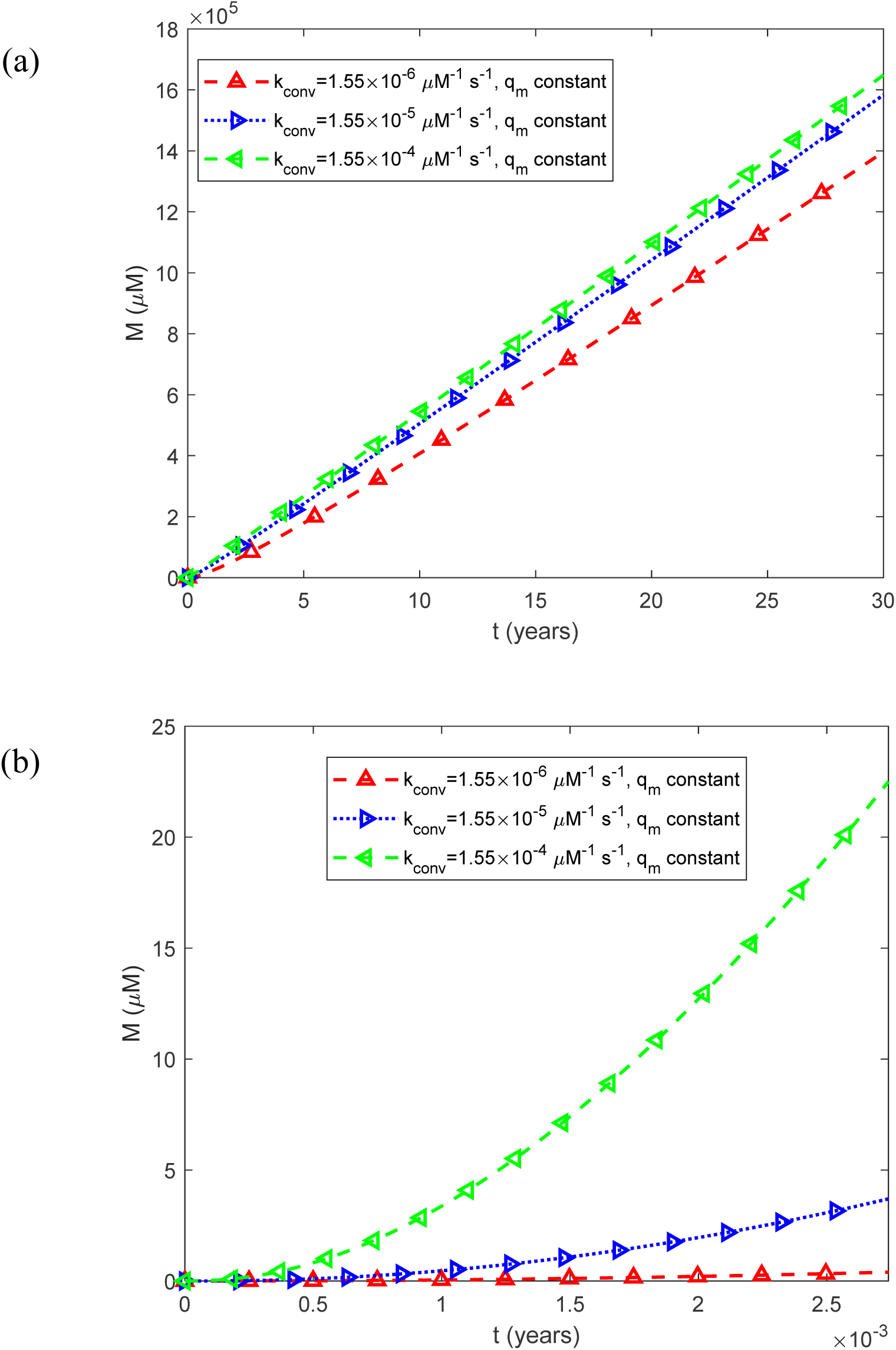
(a) Fibril mass concentration *M* (*t*) , expressed as the molar concentration of IAPP monomer equivalents incorporated into all fibrils, as a function of time for a constant (time-averaged) monomer release rate of 1.25*q_m_*_0_ . (b) As in (a), but restricted to the time interval [0, 0.0027 years], equivalent to 1 day, illustrating the effect of oscillatory monomer release on the short-term dynamics of total fibril mass. Results are shown for three representative values of the rate constant for the conversion of IAPP oligomers into elongation-competent fibrils, *k_conv_* .

**Fig. S38.**
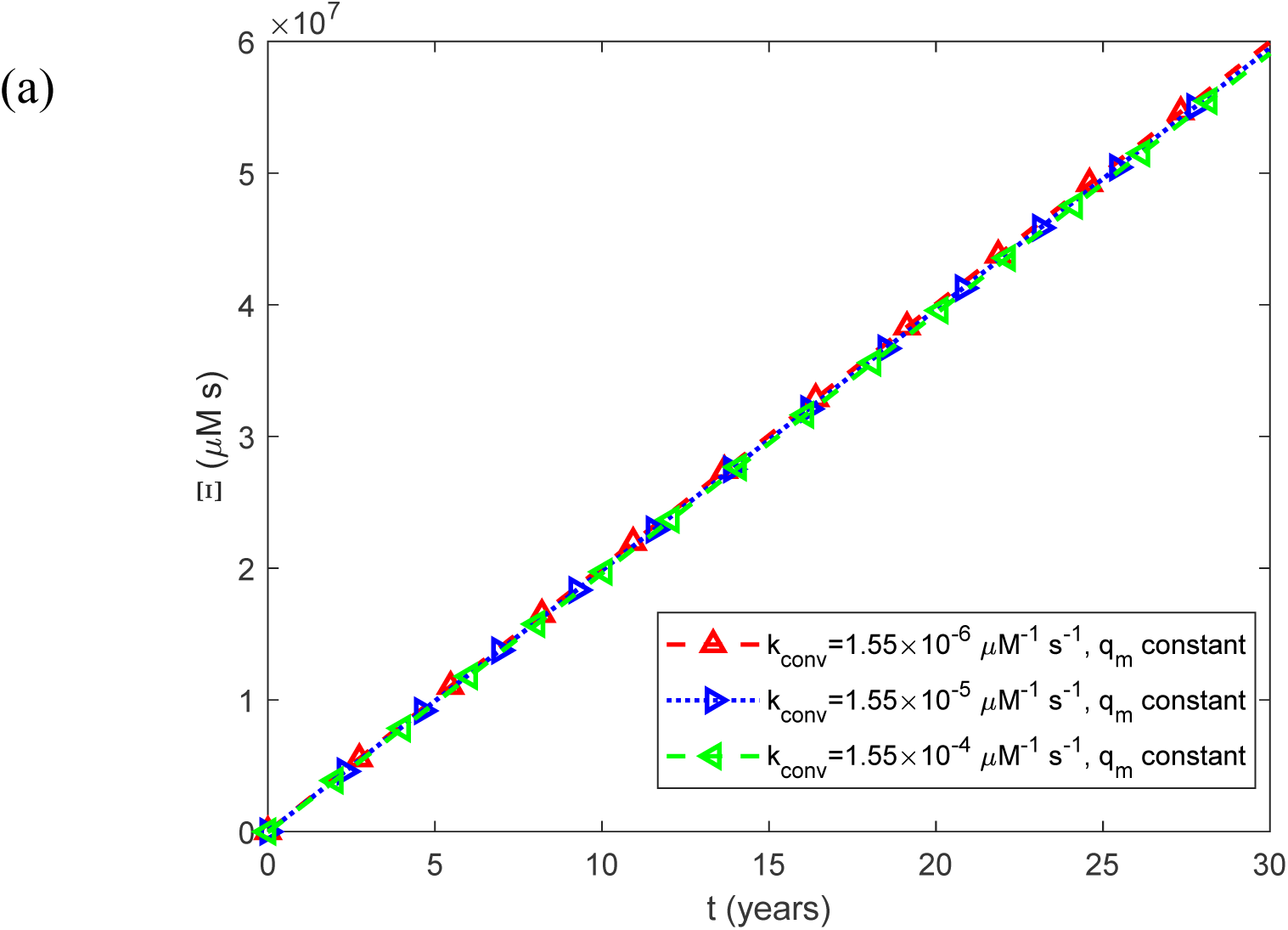
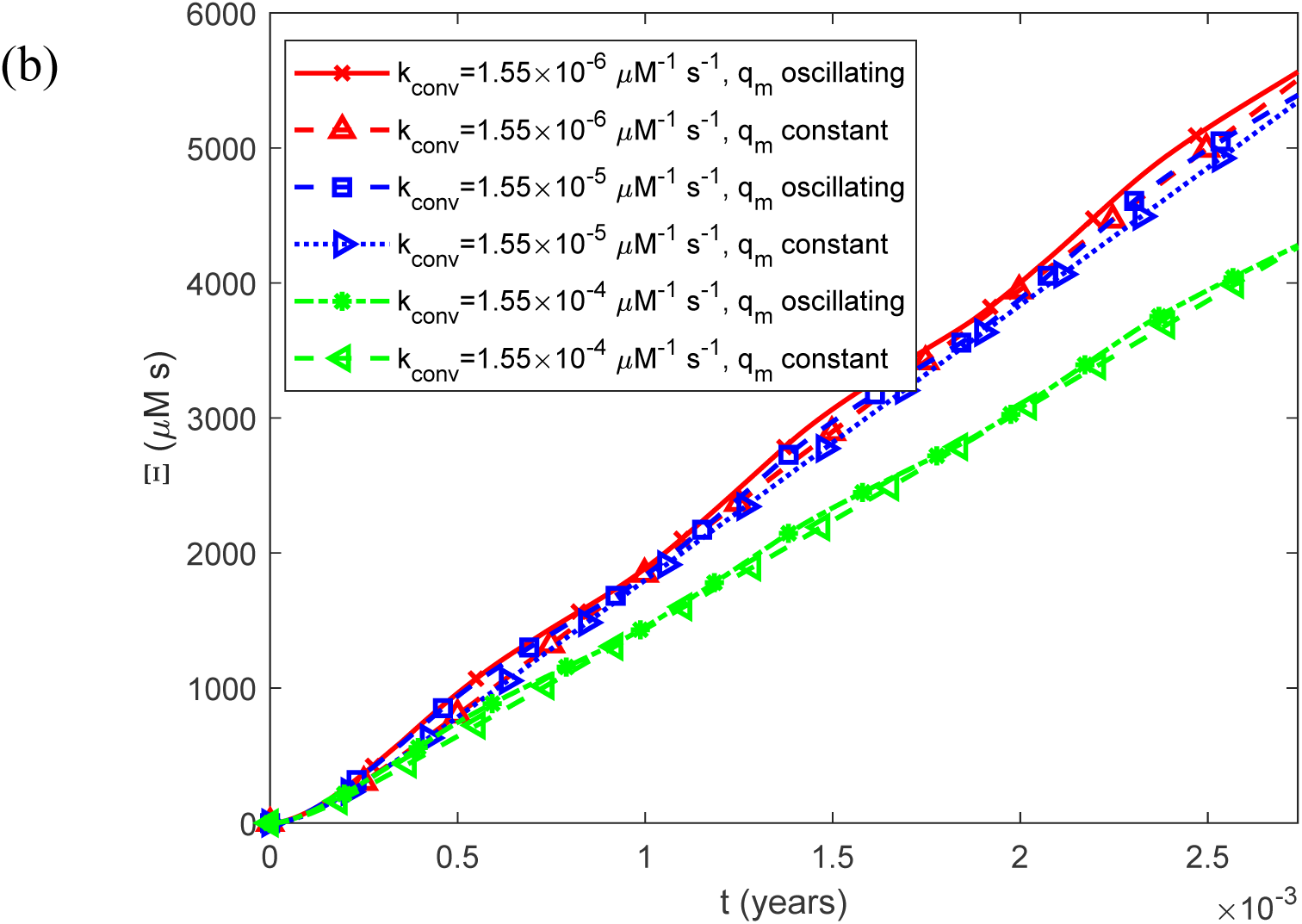
(a) Accumulated cytotoxicity (integrated IAPP oligomer exposure) as a function of time, Ξ(*t*) , for a constant (time-averaged) monomer release rate of 1.25*q_m_*_0_ . (b) As in (a), but restricted to the time interval [0, 0.0027 years], equivalent to 1 day, illustrating the effect of oscillatory monomer release on the short-term dynamics of accumulated cytotoxicity. Results are shown for three representative values of the rate constant for the conversion of IAPP oligomers into elongation-competent fibrils, *k_conv_*.

**Fig. S39.**
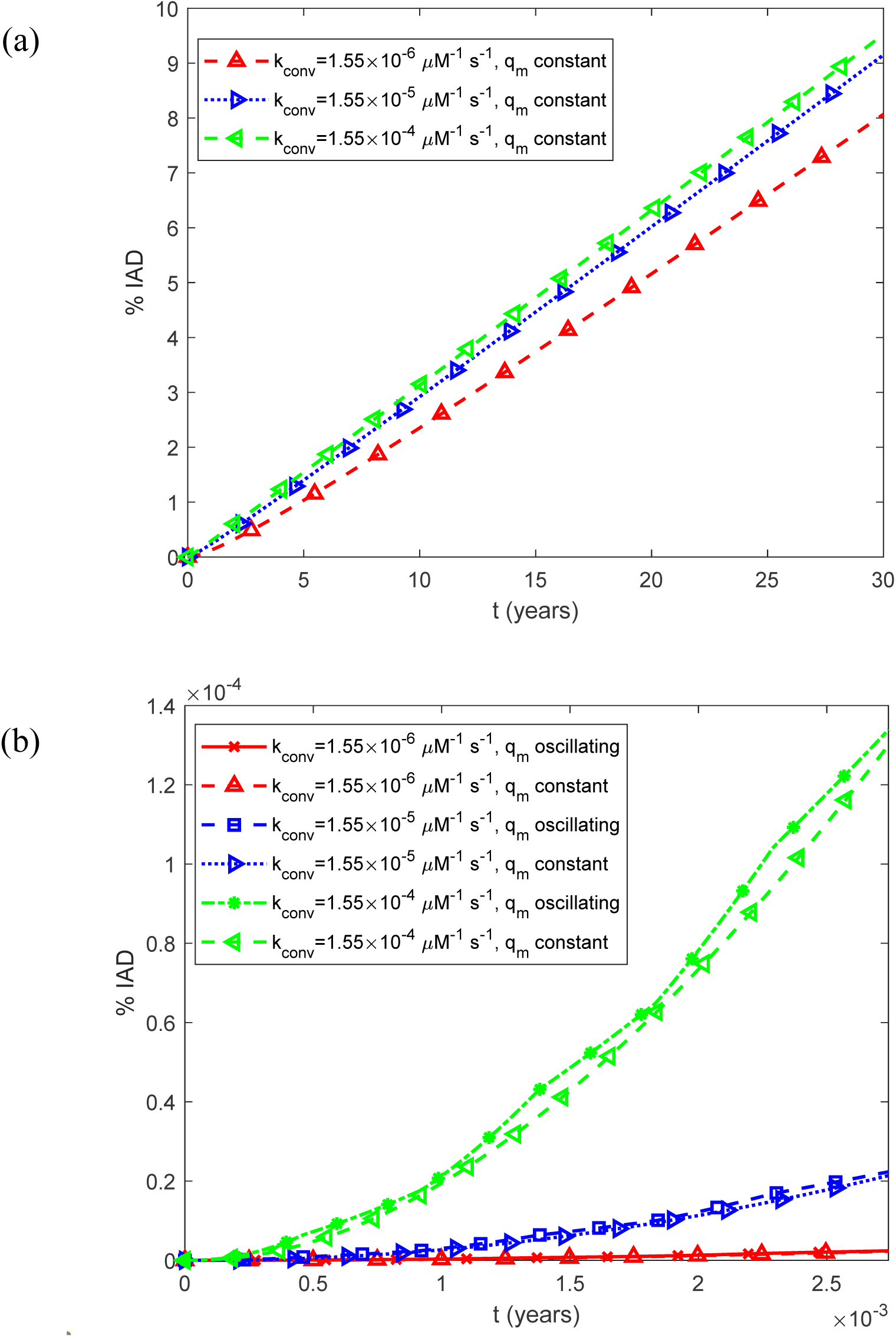
(a) Predicted percentage of the islet occupied by IADs, interpreted as the expected histological area fraction under the sterpeological assumption described in Section 2.2, as a function of time, %IADs, for a constant (time-averaged) monomer release rate of 1.25*q_m_*_0_ . (b) As in (a), but restricted to the time interval [0, 0.0027 years], equivalent to 1 day, illustrating the effect of oscillatory monomer release on the short-term dynamics of amyloid deposition. Results are shown for three representative values of the rate constant for the conversion of IAPP oligomers into elongation-competent fibrils, *k_conv_* .

## References

1. Alcazar, O. & Buchwald, P. (2019) Concentration-Dependency and Time Profile of Insulin Secretion: Dynamic Perifusion Studies With Human and Murine Islets. Frontiers in Endocrinology, 10, 680.

2. Aplin, A.C., Aghazadeh, Y., Mohn, O.G., & Hull-Meichle, R.L. (2024) Role of the Pancreatic Islet Microvasculature in Health and Disease. Journal of Histochemistry & Cytochemistry, 72, 711– 728.

3. Beck, J.V. & Arnold, K.J. (1977) Parameter Estimation in Science and Engineering. New York: Wiley.

4. Benilova, I., Karran, E., & De Strooper, B. (2012) The toxic Aβ oligomer and Alzheimer’s disease: an emperor in need of clothes. Nature Neuroscience, 15, 349–357.

5. Bennett, R.G., Duckworth, W.C., & Hamel, F.G. (2000) Degradation of amylin by insulin-degrading enzyme. Journal of Biological Chemistry, 275, 36621–36625.

6. Bertram, R. & Pernarowski, M. (1998) Glucose diffusion in pancreatic islets of Langerhans. Biophysical Journal, 74, 1722–1731.

7. Brissova, M., Shostak, A., Fligner, C.L., Revetta, F.L., Washington, M.K., Powers, A.C., & Hull, R.L. (2015) Human Islets Have Fewer Blood Vessels than Mouse Islets and the Density of Islet Vascular Structures Is Increased in Type 2 Diabetes. Journal of Histochemistry & Cytochemistry, 63, 637–645.

8. Buchanan, L.E., Dunkelberger, E.B., Tran, H.Q., Cheng, P.-N., Chiu, C.-C., Cao, P., Raleigh, D.P., De Pablo, J.J., Nowick, J.S., & Zanni, M.T. (2013) Mechanism of Iapp amyloid fibril formation involves an intermediate with a transient β-sheet. Proceedings of the National Academy of Sciences, 110, 19285–19290.

9. Buell, A.K. (2019) The growth of amyloid fibrils: rates and mechanisms. Biochemical Journal, 476, 2677–2703.

10. Cavicchi, R.E., King, J., & Ripple, D.C. (2018) Measurement of Average Aggregate Density by Sedimentation and Brownian Motion Analysis. Journal of Pharmaceutical Sciences, 107, 1304– 1312.

11. Clark, A., Charge, S.B.P., Badman, M.K., & De Koning, E.J.P. (1996) Islet amyloid in type 2 (non-insulin-dependent) diabetes. Apmis, 104, 12–18.

12. Clark, A. & Nilsson, M.R. (2004) Islet amyloid: a complication of islet dysfunction or an aetiological factor in Type 2 diabetes? Diabetologia, 47, 157–169.

13. Cottle, L., Gan, W.J., Gilroy, I., Samra, J.S., Gill, A.J., Loudovaris, T., Thomas, H.E., Hawthorne, W.J., Kebede, M.A., & Thorn, P. (2021) Structural and functional polarisation of human pancreatic beta cells in islets from organ donors with and without type 2 diabetes. Diabetologia, 64, 618–629.

14. Crank, J. (1975) The Mathematics of Diffusion. Oxford: Clarendon Press.

15. Dear, A.J., Michaels, T.C.T., Meisl, G., Klenerman, D., Wu, S., Perrett, S., Linse, S., Dobson, C.M., & Knowles, T.P.J. (2020) Kinetic diversity of amyloid oligomers. Proceedings of the National Academy of Sciences, 117, 12087–12094.

16. Defronzo, R.A. (2009) Banting Lecture. From the triumvirate to the ominous octet: a new paradigm for the treatment of type 2 diabetes mellitus. Diabetes, 58, 773–795.

17. Dorrigiv, D., Simeone, K., Communal, L., Kendall-Dupont, J., St-Georges-Robillard, A., Péant, B., Carmona, E., Mes-Masson, A.-M., & Gervais, T. (2021) Microdissected Tissue vs Tissue Slices—A Comparative Study of Tumor Explant Models Cultured On-Chip and Off-Chip. Cancers, 13, 4208.

18. Duckworth, W.C., Bennett, R.G., & Hamel, F.G. (1998) Insulin Degradation: Progress and Potential. Endocrine Reviews, 19, 608–624.

19. Emerging Risk Factors Collaboration (2023) Life expectancy associated with different ages at diagnosis of type 2 diabetes in high-income countries: 23 million person-years of observation. The Lancet Diabetes & Endocrinology, 11, 731–742.

20. Engel, M.F.M., Khemtémourian, L., Kleijer, C.C., Meeldijk, H.J.D., Jacobs, J., Verkleij, A.J., De Kruijff, B., Killian, J.A., & Höppener, J.W.M. (2008) Membrane damage by human islet amyloid polypeptide through fibril growth at the membrane. Proceedings of the National Academy of Sciences, 105, 6033–6038.

21. Eržen, S., Tonin, G., Jurišić Eržen, D., & Klen, J. (2024) Amylin, Another Important Neuroendocrine Hormone for the Treatment of Diabesity. International Journal of Molecular Sciences, 25, 1517.

22. Germanos, M., Gao, A., Taper, M., Yau, B., & Kebede, M.A. (2021) Inside the Insulin Secretory Granule. Metabolites, 11, 515.

23. Gurlo, T., Ryazantsev, S., Huang, C., Yeh, M.W., Reber, H.A., Hines, O.J., O’brien, T.D., Glabe, C.G., & Butler, P.C. (2010) Evidence for proteotoxicity in beta cells in type 2 diabetes: toxic islet amyloid polypeptide oligomers form intracellularly in the secretory pathway. American Journal of Pathology, 176, 861–869.

24. Haataja, L., Gurlo, T., Huang, C.J., & Butler, P.C. (2008) Islet amyloid in type 2 diabetes, and the toxic oligomer hypothesis. Endocrine Reviews, 29, 303–316.

25. Hassan, S., White, K., & Terry, C. (2022) Linking hiapp misfolding and aggregation with type 2 diabetes mellitus: a structural perspective. Bioscience Reports, 42, BSR20211297.

26. Hayden, M.R., Karuparthi, P.R., Manrique, C.M., Lastra, G., Habibi, J., & Sowers, J.R. (2007) Longitudinal ultrastructure study of islet amyloid in the Hip rat model of type 2 diabetes mellitus. Experimental Biology and Medicine, 232, 772–779.

27. Henquin, J.-C. (2019) The challenge of correctly reporting hormones content and secretion in isolated human islets. Molecular Metabolism, 30, 230–239.

28. Hogan, M.F. & Hull, R.L. (2017) The islet endothelial cell: a novel contributor to beta cell secretory dysfunction in diabetes. Diabetologia, 60, 952–959.

29. Hull, R.L., Westermark, G.T., Westermark, P., & Kahn, S.E. (2004) Islet Amyloid: A Critical Entity in the Pathogenesis of Type 2 Diabetes. The Journal of Clinical Endocrinology & Metabolism, 89, 3629–3643.

30. Iashchishyn, I.A., Sulskis, D., Nguyen Ngoc, M., Smirnovas, V., & Morozova-Roche, L.A. (2017) Finke-Watzky Two-Step Nucleation-Autocatalysis Model of S100A9 Amyloid Formation: Protein Misfolding as “Nucleation” Event. Acs Chemical Neuroscience, 8, 2152–2158.

31. Jaikaran, E.T.A.S. & Clark, A. (2001) Islet amyloid and type 2 diabetes: from molecular misfolding to islet pathophysiology. Biochimica et Biophysica Acta (Bba) - Molecular Basis of Disease, 1537, 179–203.

32. Kanatsuka, A., Kou, S., & Makino, H. (2018) Iapp/amylin and β-cell failure: implication of the risk factors of type 2 diabetes. Diabetology International, 9, 143–157.

33. Kuznetsov, A.V. (2024) Numerical and analytical simulation of the growth of amyloid-β plaques. Journal of Biomechanical Engineering, 146, 061004.

34. Kuznetsov, A.V. (2025) A criterion characterizing accumulated neurotoxicity of Aβ oligomers in Alzheimer’s disease. *Proceedings of the Royal Society A: Mathematical*, Physical and Engineering Sciences, 481, 20240652.

35. Kuznetsov, A.V. (2026a) Predicting Biological Age Using an Accumulated Neurotoxicity Biomarker for Amyloid Beta Oligomers. Mathematical Medicine and Biology, 43, 153–176.

36. Kuznetsov, A.V. (2026b) Investigating a Relation between Amyloid Beta Plaque Burden and Accumulated Neurotoxicity Caused by Amyloid Beta Oligomers. Medical & Biological Engineering & Computing, 64, 2747–2768.

37. Kuznetsov, I.A. & Kuznetsov, A.V. (2019) Investigating sensitivity coefficients characterizing the response of a model of tau protein transport in an axon to model parameters. Computer Methods in Biomechanics and Biomedical Engineering, 22, 71–83.

38. Li, X., Zhang, L., Meshinchi, S., Dias-Leme, C., Raffin, D., Johnson, J.D., Treutelaar, M.K., & Burant, C.F. (2006) Islet Microvasculature in Islet Hyperplasia and Failure in a Model of Type 2 Diabetes. Diabetes, 55, 2965–2973.

39. Low, J.T., Mitchell, J.M., Do, O.H., Bax, J., Rawlings, A., Zavortink, M., Morgan, G., Parton, R.G., Gaisano, H.Y., & Thorn, P. (2013) Glucose principally regulates insulin secretion in mouse islets by controlling the numbers of granule fusion events per cell. Diabetologia, 56, 2629–2637.

40. Mclean, C.A., Cherny, R.A., Fraser, F.W., Fuller, S.J., Smith, M.J., Beyreuther, K., Bush, A.I., & Masters, C.L. (1999) Soluble pool of Aβ amyloid as a determinant of severity of neurodegeneration in Alzheimer’s disease. Annals of Neurology, 46, 860–866.

41. Mizgalska, K., Al-Aani, U., Aljaidah, Y., Panek, D., Chaari, A., & Bajda, M. (2025) Inhibition of Human Amylin Aggregation: In Silico and In Vitro Studies. Acs Omega, 10, 52269–52288.

42. Morris, A.M., Watzky, M.A., Agar, J.N., & Finke, R.G. (2008) Fitting neurological protein aggregation kinetic data via a 2-step, Minimal/“Ockham’s Razor” model: The Finke-Watzky mechanism of nucleation followed by autocatalytic surface growth. Biochemistry, 47, 2413–2427.

43. Nedumpully-Govindan, P. & Ding, F. (2015) Inhibition of Iapp aggregation by insulin depends on the insulin oligomeric state regulated by zinc ion concentration. Scientific Reports, 5, 8240.

44. Paulsson, J.F., Andersson, A., Westermark, P., & Westermark, G.T. (2006) Intracellular amyloid-like deposits contain unprocessed pro-islet amyloid polypeptide (proiapp) in beta cells of transgenic mice overexpressing the gene for human Iapp and transplanted human islets. Diabetologia, 49, 1237–1246.

45. Pillay, K. & Govender, P. (2014) Novel insights into amylin aggregation. Biotechnology & Biotechnological Equipment, 28, 123–135.

46. Polonsky, K.S., Given, B.D., Hirsch, L.J., Tillil, H., Shapiro, E.T., Beebe, C., Frank, B.H., Galloway, J.A., & Van Cauter, E. (1988) Abnormal Patterns of Insulin Secretion in Non-Insulin-Dependent Diabetes Mellitus. The New England Journal of Medicine, 318, 1231–1239.

47. Raleigh, D., Zhang, X., Hastoy, B., & Clark, A. (2017) The β-cell assassin: Iapp cytotoxicity. Journal of Molecular Endocrinology, 59, R121–R140.

48. Rivera, J.F., Costes, S., Gurlo, T., Glabe, C.G., & Butler, P.C. (2014) Autophagy defends pancreatic β cells from human islet amyloid polypeptide-induced toxicity. Journal of Clinical Investigation, 124, 3489–3500.

49. Rorsman, P. & Renström, E. (2003) Insulin granule dynamics in pancreatic beta cells. Diabetologia, 46, 1029–1045.

50. Rutter, G.A., Pullen, T.J., Hodson, D.J., & Martinez-Sanchez, A. (2015) Pancreatic β-cell identity, glucose sensing and the control of insulin secretion. Biochemical Journal, 466, 203–218.

51. Sadegh Zadeh, K. & Montas, H.J. (2010) A class of exact solutions for biomacromolecule diffusion– reaction in live cells. Journal of Theoretical Biology, 264, 914–933.

52. Saravanan, M.S., Ryazanov, S., Leonov, A., Nicolai, J., Praest, P., Giese, A., Winter, R., Khemtemourian, L., Griesinger, C., & Killian, J.A. (2019) The small molecule inhibitor anle145c thermodynamically traps human islet amyloid peptide in the form of non-cytotoxic oligomers. Scientific Reports, 9, 19023.

53. Shen, Y., Joachimiak, A., Rosner, M.R., & Tang, W.-J. (2006) Structures of human insulin-degrading enzyme reveal a new substrate recognition mechanism. Nature, 443, 870–874.

54. Shome, G., Mondal, R., Deb, S., Roy, J., Mandal, A.K., & Benito-León, J. (2025) Bridging Pancreatic Amyloidosis and Neurodegeneration: The Emerging Role of Amylin in Diabetic Dementia. International Journal of Molecular Sciences, 26, 5021.

55. Solis-Herrera, C., Triplitt, C., Cersosimo, E., & Defronzo, R.A. (2025) Pathogenesis of Type 2 Diabetes Mellitus. Endotext (Feingold, K.R., Adler, R.A., Ahmed, S.F., et al. eds). South Dartmouth (Ma): MDText.com, Inc. Available at: https://www.ncbi.nlm.nih.gov/books/NBK279115/ (last updated 22 December 2025).

56. Ullsten, S., Bohman, S., Oskarsson, M.E., Nilsson, K.P.R., Westermark, G.T., & Carlsson, P.-O. (2017) Islet amyloid deposits preferentially in the highly functional and most blood-perfused islets. Endocrine Connections, 6, 458–468.

57. Viola, K.L. & Klein, W.L. (2015) Amyloid β oligomers in Alzheimer’s disease pathogenesis, treatment, and diagnosis. Acta Neuropathologica, 129, 183–206.

58. Wang, R., Liu, Y., Liang, Y., Zhou, L., Chen, M.-J., Liu, X.-B., Tan, C.-L., & Chen, Y.-H. (2023) Regional differences in islet amyloid deposition in the residual pancreas with new-onset diabetes secondary to pancreatic ductal adenocarcinoma. World Journal of Gastrointestinal Surgery, 15, 1703–1711.

59. Warren, D., Sitton, J., & Kurouski, D. (2026) Structural and morphological dynamics of “on-path” and “off-path” oligomers of human islet amyloid polypeptide. Protein Science, 35, e70523.

60. Watzky, M.A., Finney, E.E., & Finke, R.G. (2008) Transition-Metal Nanocluster Size vs Formation Time and the Catalytically Effective Nucleus Number: A Mechanism-Based Treatment. Journal of the American Chemical Society, 130, 11959–11969.

61. Weibel, E.R. (1979) Stereological Methods, Vol. 1: Practical Methods for Biological Morphometry. London: Academic Press.

62. Westermark, G.T. & Westermark, P. (2008) Transthyretin and Amyloid in the Islets of Langerhans in Type-2 Diabetes. Experimental Diabetes Research, 2008, 429274.

63. Westermark, G.T., Steiner, D.F., Gebre-Medhin, S., Engström, U., & Westermark, P. (2000) Pro islet amyloid polypeptide (Proiapp) immunoreactivity in the islets of Langerhans. Upsala Journal of Medical Sciences, 105, 97–106.

64. Westermark, P. (2011) Amyloid in the islets of Langerhans: Thoughts and some historical aspects. Upsala Journal of Medical Sciences, 116, 81–89.

65. Westermark, P., Andersson, A., & Westermark, G.T. (2011) Islet amyloid polypeptide, islet amyloid, and diabetes mellitus. Physiological Reviews, 91, 795–826.

66. Zi, Z. (2011) Sensitivity analysis approaches applied to systems biology models. Iet Systems Biology, 5, 336–346.

67. Zraika, S., Hull, R.L., Verchere, C.B., Clark, A., Potter, K.J., Fraser, P.E., Raleigh, D.P., & Kahn, S.E. (2010) Toxic oligomers and islet beta cell death: guilty by association or convicted by circumstantial evidence? Diabetologia, 53, 1046–1056.

